# A Proteome-Wide Gain-of-Function Atlas Mapping the Landscape of Chemotherapy Response

**DOI:** 10.1101/2025.09.02.673780

**Authors:** Kyeong Beom Jo, Mohammed M. Alruwaili, Da-Eun Kim, Yongjun Koh, Kwontae You, Ji-sun Kim, Hyeyeon Kim, Saba Sane, Yanqi Guo, Atina G. Cote, Nidhi Sahni, Frederick P. Roth, David E. Hill, Jung-Hyun Choi, Hunsang Lee, Kenneth A. Matreyek, Kyle K.-H. Farh, Jong-Eun Park, HyunKyung Kim, Andrei V. Bakin, Dae-Kyum Kim

## Abstract

**Background:** Conventional chemotherapies remain the core therapeutic backbone of cancer treatment, yet the rapid emergence of chemoresistance often compromises their clinical efficacy. Although large-scale functional genomics has transformed precision oncology, existing reference maps rely predominantly on loss-of-function paradigms and lack systematic data on gain-of-function (GOF) events that commonly drive clinical resistance.

**Methods:** To address this gap, we engineered a site-specific Bxb1 integrase landing-pad system in HeLa cells, enabling single-copy integration of a human proteome-wide library [18,388 open reading frames (ORFs)] at the AAVS1 locus under a uniform EF1α promoter. We systematically profiled cellular fitness across 16 chemotherapeutics. We integrated this GOF perturbation screen with bulk and single-cell RNA sequencing, FACS-based lysosomal functional screening, clinical cohort survival analyses, and preclinical xenograft modeling.

**Results:** Stratified modeling of over 230,000 gene-drug interactions defined distinct functional categories that are strongly enriched for clinical cancer biomarkers and prognostic relevance. Global network analysis and organelle-specific lysosomal functional screening uncovered that chemotherapeutic agents, most prominently the nucleoside analog TAS-102 (Lonsurf), converge upon a shared survival hub centered on the *BCL-2* family-associated autophagy-endolysosomal network. Mechanistically, TAS-102 triggered lysosomal activation as an adaptive survival response. Co-treatment with the BCL-2 family inhibitor ABT-263 (Navitoclax) attenuated this TAS-102-induced lysosomal response and synergistically reversed TAS-102 resistance across colorectal and pancreatic cancer models *in vitro* and in colorectal cell line- and patient-derived xenograft (CDX, PDX) models *in vivo*.

**Conclusions:** Our study establishes an open-access, proteome-wide GOF reference atlas that systematically maps chemotherapeutic responses, recapitulates canonical resistance biology, and nominates rational drug combinations. This systems-genomics resource bridges massive-scale perturbation mapping with actionable target discovery, providing a translational framework to overcome therapeutic resistance and reprogram cancer drug responses.

## Background

Precision oncology aims to match individual tumor profiles with targeted therapeutic interventions to maximize clinical response [1–3]. However, conventional chemotherapies remain the core therapeutic backbone for cancer treatment worldwide, offering curative potential or significant disease control for millions of patients [4–6]. Despite their widespread efficacy, standard chemotherapeutic regimens have limited clinical utility because of a narrow therapeutic window and the rapid emergence of chemoresistance [5,6]. Rational drug combinations, particularly pairing conventional chemotherapeutics with targeted agents, are among the most effective strategies for restoring therapeutic sensitivity and overcoming resistance [7,8]. Developing these rational combinations relies fundamentally on mapping how cellular networks respond to chemical perturbation across diverse therapeutic modalities.

Although targeted agents are rationally designed against specific molecular drivers and frequently combined with standard-of-care chemotherapies, identifying the genetic determinants that govern chemoresistance across the human proteome remains a major challenge [9–11]. Because aberrant protein expression broadly shapes tumor phenotypes [12], systematically evaluating how cancer cells respond to genetic perturbations across diverse therapeutic stresses is essential to reveal drug activity and guide rational combination strategies [13]. In particular, gain-of-function (GOF) alterations, such as gene amplification and transcriptional upregulation, frequently drive clinical resistance, making systematic overexpression mapping vital for uncovering dosage-sufficient drivers that loss-of-function (LOF) screens cannot capture [14–16].

However, despite the critical need for systematic GOF mapping, existing large-scale functional genomics initiatives, such as the Dependency Map [17], are intrinsically anchored in LOF paradigms, leaving GOF resistance mechanisms largely unmapped. Meanwhile, previous GOF efforts have largely been restricted to isolated drug contexts [18,19], limited expression library sizes [20], or non-cancerous cell models [21,22] using CRISPRa and lentivirus systems [23,24], failing to provide a unified reference framework that captures global therapeutic response networks. Consequently, it remains unresolved whether distinct chemotherapeutic classes trigger drug-specific responses or converge on shared, systems-level resistance networks.

Here, we present a human proteome-wide GOF atlas mapping cellular responses across 16 chemotherapeutic agents. Utilizing Bxb1-mediated single-copy ORF integration at the AAVS1 locus in HeLa cells [25–29], we established a standardized platform that enables directly comparable drug response measurements across genes and treatments. By anchoring this functional screen with bulk transcriptomics, single-cell RNA sequencing, and clinical cohort benchmarking, we demonstrated the robust translational validity of our framework. Global network analysis coupled with high-throughput lysosomal functional screening revealed that diverse chemotherapeutic stresses converge upon a central survival hub governed by the *BCL-2* family-associated autophagy-endolysosomal network. Targeting this vulnerability by pairing TAS-102 (Lonsurf) with the BCL-2 family inhibitor ABT-263 (Navitoclax) abrogated adaptive lysosomal activation and effectively reversed drug resistance across colorectal and pancreatic cancer models *in vitro* and in colorectal cell line-derived and patient-derived xenograft (PDX) models *in vivo*. Our study establishes a foundational GOF reference map that directly translates proteome-scale perturbation profiling into resistance-associated prognostic biomarkers and actionable combination strategies to overcome therapeutic resistance in cancer.

## Methods

Detailed experimental protocols, reagent catalogs, primer and antibody tables, and bioinformatic parameters are provided in **Additional file 1: Supplementary Methods**.

### Cell culture, single-integration validation, and HeLa ORFeome pool construction

T-REx-HeLa cells engineered with a single Bxb1 recombinase landing pad at the safe-harbor AAVS1 locus driven by a constitutive EF1α promoter (HeLa EF1α cells; generously provided by Dr. Frederick P. Roth) [30], alongside human colorectal (HT-29, HCT 116 *p53* WT/KO, and RKO *p53* WT/KO) and pancreatic (MIA PaCa-2 and hTERT-HPNE) cell lines were cultured as recommended by ATCC. Single-copy site-specific integration at the AAVS1 landing pad was validated by co-transfecting HeLa EF1α cells with the Bxb1 integrase vector (pCAG-NLS-HA-coBxb) followed by an equimolar (1:1) mixture of promoterless attB-EGFP and attB-mCherry donor plasmids and quantifying mutually exclusive single-positive (EGFP⁺/mCherry⁻ and EGFP⁻/mCherry⁺) versus double-positive events by flow cytometry two weeks post-transfection [26,27].

To construct the proteome-wide GOF library, sequence-verified entry clones from the human ORFeome v9.1 [25,31] and CCSB Delta (Δ) collections (18,388 unique ORFs) were pooled at equimolar ratios and transferred into the pDEST-HC-REC destination vector via 5-day en masse Gateway LR cloning, followed by electroporation into 10-β electrocompetent *E. coli* to achieve ∼1,000× library coverage [29,32]. HeLa EF1α cells seeded across twelve 10 cm dishes (1.3 × 10^6^ cells/dish) were sequentially transfected with 10 µg of pCAG-NLS-HA-coBxb and 10 µg of the ORFeome library using Lipofectamine 3000, following the manufacturer’s protocol. Cells were pre-sorted for mCherry⁺ expression 48 h later and post-sorted on day 5 post-transfection for mCherry⁺/mTagBFP2⁻ cells on a Sony MA900 sorter, expanded, and cryopreserved at 3 × 10^6^ cells per vial.

### Chemotherapeutic screening, sequencing prep, and high-saturation retesting (HSR)

Half-maximal inhibitory concentrations (IC_50_) at 72 h were determined in HeLa cells using AlamarBlue viability assays [20]. We profiled 16 chemotherapeutics spanning four mechanistic classes [33] (**Fig. 1A**): antimetabolites [5-FU (0.1925 µM), TFT (0.75 µM), EdU (0.875 µM), FdU (25 nM), 6-MP (2 µM), and TAS-102 (0.8 µM)], DNA cross-linking agents [MMC (14 nM), CDDP (0.333 µM), CBDCA (3.37 µM), and BCNU (7 µM)], DNA strand-break agents [DOX (5 nM), BLM (3.3 µM), VP-16 (0.625 µM), and CPT-11 (0.8 µM)], and microtubule inhibitors [PTX (2 nM) and VBL (0.3 nM)]. Screens were conducted across five independent experimental batches, each incorporating untreated baseline controls and batch-matched DMSO vehicle controls (n = 3 biological replicates per drug; duplicates for 6-MP, CPT-11, and VBL due to insufficient ORF recovery in one replicate, 5 baseline + 15 DMSO + 45 drug-treated = totaling 65 samples). For each condition, ≥3 × 10^6^ HeLa ORFeome cells (150× coverage) were treated at the respective IC₅₀ dose for 72 h, passaged at 3 × 10^6^ cells into fresh drug-containing medium for an additional 48 h, and allowed to recover in drug-free medium for 3 days before genomic DNA (gDNA) extraction (Qiagen Blood and Cell Culture DNA Midi Kit).

**Fig. 1.**
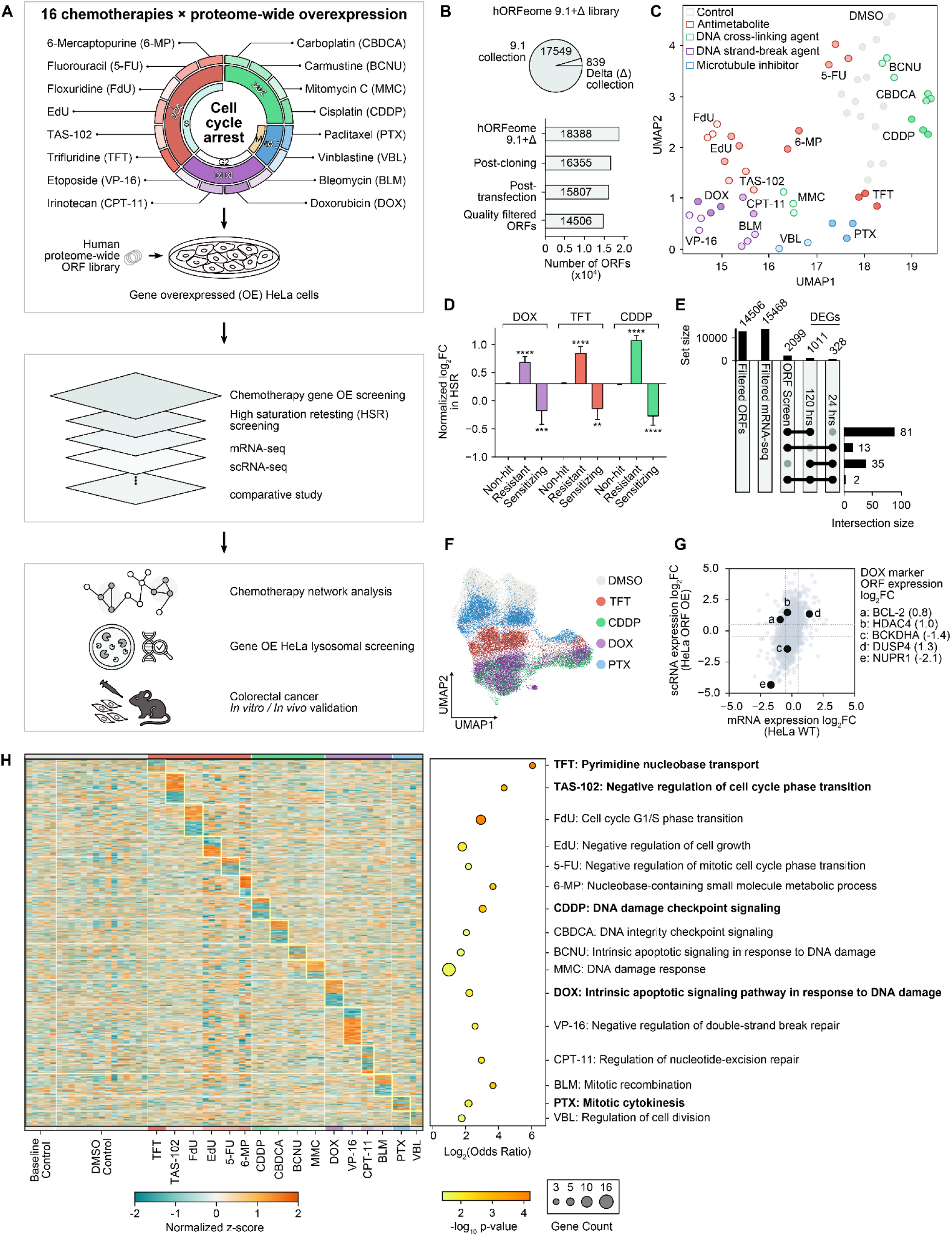
Proteome-wide GOF screening framework and multiscale response atlas across 16 chemotherapeutic modalities. **(A)** Conceptual schematic of the proteome-wide ORF overexpression screening pipeline in HeLa cells across 16 chemotherapeutic agents. **(B)** Library representation and sequencing recovery metrics across sequential experimental stages of the hORFeome 9.1+Δ collection. **(C)** 2D UMAP projection of 16 chemotherapeutic agents clustered by ORF profiles. **(D)** Screening fidelity validation demonstrating normalized log_2_FC distributions from high-saturation retesting (HSR) across non-hit, resistant, and sensitizing ORF candidates. **(E)** UpSet plot showing overlap of differentially expressed genes (DEGs) between the doxorubicin drug response screen and the HeLa wild-type transcriptomic screen at 24 and 120 hours of doxorubicin (DOX) treatment. **(F)** UMAP embedding of single-cell transcriptomic profiles derived from scRNA-seq of HeLa ORFeome cells under the indicated chemotherapy treatments. **(G)** Quadrant scatter plot comparing single-cell RNA expression in HeLa ORFeome cells against bulk mRNA expression in wild-type HeLa cells under DOX treatment, highlighting key representative doxorubicin marker genes. **(H)** Global scale and biological validity of the proteome-wide GOF chemoresistance atlas. Left: heatmap displaying the full dimension of the functional dataset. Right: dot plot demonstrates high biological fidelity of the dataset in capturing canonical drug-specific responses. The bar graph in panel D shows the mean ± Standard Error of the Mean (SEM). Statistical analysis was performed by one-way ANOVA with Dunnett’s multiple comparisons test against the non-hit control group; ** *p* < 0.01, *** *p* < 0.001, **** *p* < 0.0001.

Integrated full-length ORFs were selectively amplified from 18 µg of gDNA per sample via two-step nested PCR using Q5 High-Fidelity Master Mix with primers targeting the landing-pad locus (PCR1) and flanking Gateway attB sites containing 5′ NNNNN diversity pads (PCR2). Amplicons were sheared to 250-300 bp (Covaris E220), ligated to TruSeq adaptors (KAPA HyperPrep), and sequenced on an Illumina NovaSeq 6000 (≥3 million paired-end reads/sample). To validate primary screening fidelity and reproducibility via High-Saturation Retesting (HSR), 2,217 ORFs selected by primary *t*-score rankings across eight functional baskets were cherry-picked using an automated robotic platform (S&P Robotics BM3-BC), pooled via en masse LR cloning into HeLa EF1α cells, and re-screened at >1,000× coverage under DOX, TFT, and CDDP at matching IC_50_ doses. Primary resistant (log_2_ fold change (log_2_FC) ≥ 1.0, *p* < 0.01) and sensitizing (log_2_FC ≤ −1.0, *p* < 0.01) hits were benchmarked against Non-hit controls (|log_2_FC| ≤ 1.0, p ≥ 0.01) using ordinary one-way ANOVA followed by Dunnett’s test in SciPy.

### ORFeome bioinformatic pipeline, longitudinal baseline, and atlas visualization

FASTQ reads were adapter- and quality-trimmed using Cutadapt and Trimmomatic, aligned to the 18,388 ORF reference sequences (hORFeome v9.1 + Δ) using Bowtie2 (v2.5.2), and quantified with Samtools and Pysam (MAPQ ≥ 30). Raw read counts were normalized by Trimmed Mean of M-values (TMM) in edgeR (v4.0.16), and 14,506 ORFs exhibiting a median log₂TMM > −1 across baseline samples were retained as quality-filtered ORFs. After 0.001% winsorization (DescTools), upper-quantile normalization (edgeR), and linear batch correction across experimental batches and sequencing runs (limma v3.58.1), differential representation (log_2_FC) between drug-treated replicates and batch-matched DMSO controls was evaluated via 10,000 permutations. Two-tailed *p*-values from t-statistics, log₂ mean differences, and z-transformed rank-sum tests were combined into an integrated *p*-value using Liptak–Stouffer’s z-method, with Benjamini–Hochberg false discovery rate (FDR) estimation (statsmodels). Sample clustering and replicate reproducibility were assessed using UMAP (umap-learn v0.5.7), 2D PCA (scikit-learn, random_state = 77), and pairwise Pearson correlation (r) analysis (two-sample Student’s *t*-test between intra-replicate and inter-treatment pairs).

Longitudinal library uniformity from the input plasmid pool through post-transfection days 3, 7, 9, 12, 14, and 24 was quantified using counts per million (CPM), represented in Lorenz curves, and Gini coefficients. Baseline essentiality dropout from D9 to D24 was classified into unchanged, gradual decrease, and complete dropout (0 CPM at D24) groups based on reproducible depletion across PCR1 (log_2_FC ≤ −1.5) and PCR2 (log_2_FC ≤ −1.0) series. Reporter silencing ratios were defined as log₂(mCherry⁺BFP⁻ / mCherry⁻BFP⁻), where log_2_(E/S) < −1 denoted silenced ORFs, and correlated against log₁₀-transformed ORF length (bp) using Pearson’s r and OLS regression hexbin plots. For global heatmap visualization across all 65 samples, ORFs significantly altered (|log_2_FC| ≥ 0.5, integrated *p* < 0.05) in 1–4 drugs were assigned to their primary drug cluster by maximum row-wise *z*-score, ordered via Ward’s hierarchical clustering in SciPy, standardized by column- and row-wise *z*-scores, and aligned with drug-cluster pathway enrichments computed via GSEApy (v1.1.9; Enrichr API) against GO, Reactome, and KEGG databases.

### Bulk mRNA-seq and single-cell RNA-seq profiling

For bulk transcriptomics, wild-type HeLa cells treated with 5 nM DOX for 24 h or 120 h were profiled using the KAPA mRNA HyperPrep Kit on a NovaSeq 6000 and processed via the nf-core/rnaseq pipeline (v3.18.0; Trim Galore, SortMeRNA, STAR v2.7.10a alignment to GRCh38.p14, RSEM v1.3.1 quantification, and edgeR exactTest). For pooled single-cell RNA-seq, HeLa ORFeome cells treated with DMSO, TFT, CDDP, DOX, or PTX were captured on the 10x Genomics Chromium 5′ v2 platform (∼10,000 cells/channel) and aligned to GRCh38 using Cell Ranger (v6.0.2). Using Scanpy (v1.9.1), cells with <2,000 counts, <500 genes, >10% mitochondrial reads, or Scrublet doublet scores >0.4 were excluded. Counts were normalized to 10,000 per cell, log-transformed [ln(x + 1)], embedded via PCA (top 50 PCs across 2,000 highly variable genes) and UMAP, and clustered via the Leiden algorithm (resolution = 0.5) with Wilcoxon rank-sum marker detection.

### ORF functional categorization, intrinsic gene properties, and clinical benchmarking

Differentially represented ORFs (|log_2_FC| ≥ 0.5 and integrated *p* < 0.05) among the 14,506 quality-filtered ORFs were sequentially prioritized into five mutually exclusive categories: (i) Core ORFs (197 ORFs altered in ≥7 chemotherapeutics), (ii) Categorical ORFs (1,072 class-specific ORFs defined by differential median log₂FC between a given drug class and remaining classes). The remaining ORFs were classified into (iii) Peripheral ORFs (3,832 single-drug-specific ORFs), (iv) Multidrug ORFs (6,556 ORFs altered in 2–6 drugs across multiple classes), and (v) Non-respondent ORFs (2,849 ORFs). Intrinsic gene properties were compiled from hORFeome sequences (ORF length and GC content), GTEx v6 (abundance and tissue broadness) [34], ProteinHistorian (evolutionary age) [35], D^2^P^2^ (intrinsic disorder) [36], DAVID (GO terms, PubMed counts, UniProt tissue upregulation, and OMIM terms) [37–39], and CORUM v4.1 (protein complex membership) [40]. Clinical biomarker enrichment was evaluated against TheMarker database [41] across verified, non-silenced ORFs.

Baseline proliferation scores were computed from batch-wise log₂FC (DMSO versus baseline), standardized to *z*-scores per batch, and aggregated via inverse *z*-score transformation. Proliferation and drug-response *z*-scores were benchmarked against MCF-7 lentiviral OE screens (364 shared ORFs) [22], length-stratified lentiviral screens across 1-kb intervals [22,42], and NALM-6 CRISPR-Cas9 KO screens across eight shared chemotherapeutics [43]. Clinical prognostic and predictive performance was evaluated using precision–recall at rank k curves and log-rank Kaplan–Meier survival analyses across Human Protein Atlas (HPA v24.0) [44] and curated TCGA treatment cohorts [45–47].

### Drug-response similarity network, SAFE neighborhood enrichment, and pathway convergence analysis

An ORF–ORF drug-response similarity network was constructed across all 11,595 drug-responsive ORFs (|log_2_FC| ≥ 0.5, integrated *p* < 0.05 in ≥1 drug) by computing pairwise-complete Pearson correlations (r) across signed categorical response profiles (minimum 8 shared measured drugs) and connecting each ORF to its k = 10 most correlated neighbors (11,595 nodes; 100,347 edges). Node coordinates were generated using a deterministic Fruchterman–Reingold layout (60 iterations, PCG64 seed = 0), aligned by SVD, and radially equalized onto a circular disc. Correlation-rank neighborhood enrichment modeled on SAFE was performed using each node’s 100 most correlated neighbors across GO terms containing 10–600 library ORFs (one-sided hypergeometric *p* < 0.001), benchmarked against 3,000 size-matched random gene sets with within-domain Šidák and across-domain Benjamini–Hochberg adjustments.

Network-free pathway convergence was tested across 14 pre-specified apoptotic, autophagic, and endolysosomal gene sets across all 16 drugs in both directions (444 one-sided hypergeometric tests), with headline enrichments benchmarked against 20,000 random gene sets matched to the combined growth *z*-score decile distribution. The gene–drug bipartite network encompassed the 15 Core ORFs (≥7 drugs) comprising nine apoptotic/*BCL-2* family annotated Core ORFs together with the six Core axis ORFs (MARCHF1, HEBP2, FUNDC2, BCL2L2, UGCG, and DEDD2; 113 interactions). Directional concordance across the 33 responsive subunits of the obligate ESCRT-0/I/II/III, HOPS, and V-ATPase complexes under TAS-102 was evaluated using a one-sided exact binomial sign test.

### Proteome-wide LysoTracker FACS screen and PPI network analysis

HeLa ORFeome cells treated with vehicle (DMSO) for 72 h in biological triplicate (3 × 10^6^ cells/dish) were stained with 50 nM LysoTracker Green for 20 min at 37°C and sorted on a Sony MA900 sorter into mCherry⁺ populations exhibiting high (top 5%) and low (bottom 5%) lysosomal activity based on green fluorescence (GFP channel) alongside pre-sort controls. Sorted populations were expanded, sequenced, and analyzed via the ORFeome bioinformatic pipeline to identify lysosomal activators and repressors (|log_2_FC| ≥ 0.5, integrated *p* < 0.05), followed by DAVID pathway analysis and TCGA Pan-Cancer survival stratification via cBioPortal [46,47] (comparing detectable versus undetectable tumors for GHSR and median-split High versus Low groups for CCDC86 using cancer-type-stratified log-rank and Cox proportional hazards ratio (HR) models with 95% confidence interval (CI)). Overlapping hits present in at least two of the three datasets (LysoTracker high 5%, LysoTracker low 5%, and TAS-102 screen) were mapped onto the STRING database (confidence score ≥ 0.7) in Cytoscape (v3.10.4), and the largest connected subnetwork was annotated with Therapeutic Target Database (TTD) drug targets and multi-condition log₂FC pie charts using Omics Visualizer.

### *In vitro* functional validation and *in vivo* xenograft models

For immunoblotting, HeLa (0.8 µM), RKO (0.6 µM), and HCT 116, MIA PaCa-2, and HT-29 (2 µM) cells were treated with TAS-102 or DMSO for 24 h to assess *BCL-2* expression. HeLa and HCT 116 cells treated with DMSO, TAS-102 (1 µM), ABT-263 (0.5 µM for HCT 116 or 1 µM for HeLa), or their combination for 48 h were lysed, resolved by 8**–**15% SDS-PAGE, transferred to PVDF membranes, and probed with primary antibodies against *BCL-2*, TFEB, SQSTM1/p62, LAMP1, LC3B, and β-actin. In HCT 116 cells treated with DMSO, 2 µM TAS-102, 0.5 µM ABT-263, or their combination for 24 h, nuclear TFEB translocation (anti-TFEB immunofluorescence normalized to Hoechst 33342 nuclear counts across n = 5) and lysosomal proteolysis (10 µg/mL DQ Red-BSA for 24 h) were quantified by confocal microscopy (Zeiss LSM800) and ImageJ (v1.53K). Luminal lysosomal pH (n = 5/group) was measured in 96-well plates using 1 µM LysoSensor Yellow/Blue DND-160 (329/440 nm and 380/540 nm) calibrated against pH 4.5–7.0 nigericin standards. Proliferation with 100 nM Bafilomycin A1 (BafA1) and 2 µM TAS-102 in HCT 116 cells was quantified over 48 h (n = 6 replicates). Combinatorial cytotoxicity across TAS-102 (0–50 µM, 24 h) and ABT-263 (0–5,000 nM, 96 h) matrices was measured by 0.5% methylene blue staining (650 nm) and evaluated using Bliss independence (Bliss) [48], Highest Single Agent (HSA) [49], and Zero Interaction Potency (ZIP) [50] synergy models.

All animal studies were approved by the RPCCC IACUC. For cell line-derived xenografts (CDX), female SCID/CB17 mice (6–8 weeks old) were injected subcutaneously with 1 × 10^6^ HCT 116 cells in 1:1 DPBS: Matrigel. For patient-derived xenografts (PDX), colorectal tumor fragments (CRC PDX-03-26-4p; 2 × 2 × 2 mm^3^) were implanted subcutaneously into the left flank of male SCID/CB17 mice. Once tumors reached 50–100 mm^3^, mice were randomized into four groups (n = 10–11/group for CDX and n = 6–8/group for PDX): (1) vehicle, (2) ABT-263 (50 mg/kg), (3) TAS-102 (50 mg/kg), and (4) combination (50 mg/kg TAS-102 administered 4 h before 50 mg/kg ABT-263), delivered via oral gavage on a 5-day-on, 2-day-off schedule. Tumor volumes [(length × width^2^)/2] and body weights were monitored twice weekly, and endpoint complete blood counts (CBC) were analyzed using a Heska HemaTrue Analyzer. Statistical comparisons were performed in GraphPad Prism 9 using two-way ANOVA with Tukey’s multiple comparisons test (with TAS-102 and ABT-263 as main-effect factors for single-timepoint 2 × 2 factorial assays, and treatment group and time as factors for longitudinal proliferation and tumor growth curves), one-way ANOVA with Tukey’s test for CBC parameters, or Welch’s two-tailed *t*-test for endpoint tumor weights.

### Use of Large Language Model (LLM)

During the preparation of this manuscript, the authors used large language models (ChatGPT, Gemini, and Claude) solely for language editing, grammar checking, and verification of text consistency. The authors reviewed and edited all content and took full responsibility for the integrity of the manuscript.

## Results

### Systematic proteome-wide mapping of chemoresistance landscape using a site-specific ORF overexpression system

We performed a proteome-wide ORF screen using a panel of 16 chemotherapeutic agents selected to span diverse cytotoxic mechanisms and cell cycle arrest pathways [33,51,52] (**Fig. 1A**). We grouped these 16 chemotherapeutic compounds into four major functional categories representing distinct rate-limiting lethality axes and cell cycle targets. Based on the mechanisms of action, we tested i) 6 antimetabolites [fluorouracil (5-FU), trifluridine (TFT), TAS-102, EdU, floxuridine (FdU) and 6-mercaptopurine (6-MP)] that target pyrimidine/purine synthesis, thymidylate synthase activity, or S-phase progression kinetics, ii) 4 DNA cross-linking and alkylating agents [mitomycin C (MMC), cisplatin (CDDP), carboplatin (CBDCA) and carmustine (BCNU)] that induce intra- and interstrand DNA cross-links and covalent alkylation, iii) 4 DNA strand-break inducing agents [doxorubicin (DOX), bleomycin (BLM), etoposide (VP-16) and irinotecan (CPT-11)] that target topoisomerase I/II cleavage complexes or generate reactive oxygen species and iv) 2 microtubule inhibitors [paclitaxel (PTX) and vinblastine (VBL)] that disrupt mitotic spindle assembly and cytokinesis. This comprehensive panel was designed to capture both broad, multi-drug survival hubs and class-specific response modifiers across fundamental cytotoxic mechanisms.

To execute this multi-drug screen with proteome-wide coverage and strict quantitative uniformity, we engineered a site-specific recombination architecture based on a Bxb1 landing pad donor vector [26] integrated into the *AAVS1* locus of HeLa cells (**BOGO**, **<u>B</u>**xb1-landing pad human **<u>O</u>**RFeome-integrated system for a proteome-wide **<u>G</u>**ene **<u>O</u>**verexpression; **Fig. 1B, Additional file 2: Fig. S1A**). Insertion of the human ORFeome library (hORFeome 9.1 [25] + Δ collections, totaling 18,388 unique ORFs) yielded 16,355 post-cloning constructs, from which 15,807 were successfully integrated into HeLa cells (HeLa ORFeome cells; **Fig. 1B)**. We applied baseline quality filtering to prevent low-abundance ORFs from introducing stochastic noise. This established a high-confidence pool of 14,506 ORFs for downstream functional profiling (**Fig. 1B, Additional file 3: Table S1**). Unsupervised UMAP dimensionality reduction cleanly segregated the 16 chemotherapeutics into distinct clusters aligned with their primary mechanisms of action, while DMSO vehicle controls from all five batches clustered tightly along the UMAP1 axis (**Fig. 1C**). Principal component analysis (PCA) across five independent experimental and sequencing batches confirmed minimal batch variation and high technical reproducibility, with experimental samples broadly distributed and intermingled across the principal component space (**Additional file 2: Fig. S1B, left**). To verify that drug treatments elicited true mechanism-driven biological responses rather than non-specific noise, we benchmarked the functional reactions of canonical drug-gene control pairs (**Additional file 2: Fig. S1B, right**). Overexpressing direct molecular targets or metabolic modifiers produced expected, highly specific phenotypic shifts: *TYMS* overexpression selectively conferred resistance to thymidylate synthase inhibitors FdU, TFT, and TAS-102 [53,54], *TK1* and *TPMT* altered sensitivity to purine/nucleoside analogs like TAS-102 and 6-MP [55,56]. Similarly, *MGMT* protected against alkylating damage [57], *NQO1* modulated bioreductive agent response [58], and *TUBB* increased cellular sensitivity to microtubule-targeting agents PTX and VBL [59] (**Additional file 2: Fig. S1B, right**). To systematically evaluate technical reproducibility and signal specificity across the dataset, we analyzed pairwise Pearson correlation coefficients across all 45 sequencing samples (**Additional file 2: Fig. S1C, right**). Intra-replicate samples from independent experimental batches exhibited consistently high correlation coefficients, whereas inter-treatment samples across different chemotherapeutic classes displayed markedly lower, drug-specific correlations (**Additional file 2: Fig. S1C, left**).

We further verified screening fidelity through high-saturation retesting (HSR), a targeted, high-coverage approach evaluating a curated sub-library of 2,217 ORFs at >1,000× coverage across three representative treatment settings spanning major DNA-damaging and antimetabolite classes (DOX, CDDP, and TFT; **Fig. 1A**, **Fig. 1D, Additional file 2: Fig. S1D; Additional file 3: Table S3**). HSR recovered 98.5% of the 2,217 ORFs and 96.9% of the 96 primary chemotherapy response ORFs that had high quantitative accuracy (**Fig. 1D, Additional file 2: Fig. S1D**). Specific candidates across treatments displayed log_2_FC > 1 differential fold changes (**Additional file 2: Fig. S1D**), confirming exceptional signal reproducibility between primary and retest screens. *MAGEB2* conferred robust resistance to DOX [60], *GCNT2* modulated altered glycosylation states to enhance DOX sensitivity [61], *TXNDC9* acted as a protective resistance factor against TFT-induced oxidative stress [62], the nucleoside transporter *SLC28A1* mediated enhanced TFT sensitivity [63], *TBPL1* drove resistance under cisplatin pressure [64], and *GPR21* marked opposing sensitivity-conferring modulation [65] (**Additional file 2: Fig. S1D**). These representative genes highlighted balanced functional axes spanning diverse drug classes.

To benchmark against standard transcriptomics, we analyzed longitudinal bulk mRNA sequencing of wild-type (WT) HeLa cells treated with or without doxorubicin. These transcriptomic profiles were collected at 24 hours (h) to capture acute transcriptional shifts and at 120 h to match the cumulative drug-exposure duration of our overexpression platform (**Fig. 1E; Additional file 3: Table S4**). UpSet plot analysis showed minimal overlap between chemo-responsive genes and differentially expressed genes identified by bulk mRNA-seq, with less than 5% overlap at 120 h and less than 1% at 24 h (**Fig. 1E**). This limited overlap may reflect that bulk transcriptomics measures reactive, downstream transcriptional shifts in HeLa WT cells (**Additional file 2: Fig. S1E**), whereas HeLa GOF screening identifies more survival and resistance genes under the same selective therapeutic pressure (**Fig. 1E**). To bridge genomic-level survival hits with downstream expression dynamics, we performed single-cell RNA sequencing (scRNA-seq) on HeLa ORFeome cells treated with one of the representative chemotherapeutics from each mechanism of action (TFT, CDDP, DOX, and PTX; **Fig. 1F; Additional file 3: Table S5**). When comparing fold-change dynamics under DOX stress, endogenous mRNA expression in HeLa WT cells showed only modest spread near baseline, while ORF expression driven by the constitutive promoter in the GOF system exhibited a markedly broader transcriptomic distribution (**Fig. 1G**). This approach identified key resistance and sensitization markers such as *BCL-2* [66], *HDAC4* [67], *DUSP4* [68], *NUPR1* [69], and *BCKDHA* [70], of which *HDAC4* and *BCKDHA* were captured exclusively through gene overexpression (**Fig. 1G**). These drivers were organized into cluster-specific functional modules, including cell cycle and DNA repair, pro-survival signaling (Wnt and NF-κB), and programmed cell death regulation (autophagy and apoptosis; **Additional file 2: Fig. S1F**). Together, these findings demonstrate how systematic GOF profiling can identify functional resistance and sensitization drivers that are not readily detected by conventional transcriptomic profiling of drug-treated WT cells, providing a complementary approach for mapping drug response networks across diverse chemotherapeutic agents. Comprehensive quality control metrics evaluating cellular integration efficiency, longitudinal library representation, and baseline expression dynamics are detailed in **Additional file 1: Supplementary Note**, **Additional file 2: Fig. S1G–J**, and **Additional file 3: Table S2**.

To map functional response landscapes across all 16 chemotherapeutic treatments, we performed proteome-wide GOF profiling across 14,506 ORFs in 5 untreated (“baseline”), 15 DMSO controls, and 45 drug-treated conditions across 16 chemotherapeutic modalities (**Fig. 1H, left**). This dataset yielded approximately 0.9 million sample-level functional data points, synthesizing a high-confidence matrix of over 230,000 unique gene-drug interaction pairs mapped across four major therapeutic classes (**Fig. 1H, left**). Systematic pathway enrichment analysis (**Additional file 3: Table S6**) demonstrated that these differential ORFs map directly onto the canonical mechanisms of action unique to each chemotherapeutic class (**Fig. 1H, right**). Antimetabolites (TFT and 6-MP) primarily block nucleobase transport and metabolic salvage kinetics, whereas fluoropyrimidines and cell cycle agents (FdU, TAS-102, 5-FU, and EdU) are governed by G1/S transition overrides (**Fig. 1H, right**). Platinum agents (CDDP and CBDCA) and alkylators (MMC) selectively enrich for DNA damage checkpoint and interstrand cross-link repair; topoisomerase inhibitors and strand-break inducers (DOX, BCNU, VP-16, CPT-11, and BLM) converge on double-strand break repair and apoptotic threshold regulation; and microtubule agents (PTX and VBL) map tightly to mitotic spindle checkpoint and cytokinesis networks (**Fig. 1H, right**). These results establish the technical accuracy, functional specificity, and deep biological validity of our proteome-wide GOF reference atlas.

### The systematic atlas identifies functional biological drivers and potential clinical prognostic biomarkers

We categorized 14,506 quality-filtered ORFs according to their differential representation patterns, resulting in five functional groups: Core, Categorical, Multidrug, Peripheral, and Non-respondent (**Fig. 2A, Additional file 2: Fig. S2A**). Examining their cumulative distribution demonstrated that unique drug response ORFs expanded progressively with the inclusion of additional drugs (**Additional file 2: Fig. S2B**), exceeding expectations from random permutation models. A histogram of interaction counts revealed a sharp decline as the breadth of drug-response sharing increased, with only a small subset modulating responses across up to 13 agents (**Additional file 2: Fig. S2C**). Modeling the distribution of drug-response counts per ORF using a mixture of gamma and normal distributions further illustrated this transition from a right-skewed baseline of narrow-spectrum interactions (gamma component) to a broader tail of multidrug-regulating ORFs (normal component) (**Additional file 2: Fig. S2D**). Accordingly, ORFs modulating responses across ≥ 7 drugs (at least half of the maximum observed breadth of 13 agents) were designated as Core ORFs, whereas ORFs specific to a single chemotherapeutic agent were defined as Peripheral ORFs (**Additional file 2: Fig. S2E**), and those affecting 2 to 6 drugs across different mechanisms of action were classified as Multidrug ORFs. Within Categorical ORFs, drug response ORFs for DNA strand break agents were most prevalent (**Additional file 2: Fig. S2F**), while Multidrug ORFs exhibited defined breakdowns across drug combinations (**Additional file 2: Fig. S2G**). To maintain a uniform discovery threshold across all 16 chemotherapeutic agents despite differences in replicate number and ORF detection across conditions, we prioritized permutation-derived integrated *p*-values (|log_2_FC| ≥ 0.5 and *p* < 0.05) for pan-atlas categorization while reporting FDR *q*-values for all gene-drug pairs (**Additional file 2: Fig. S2H; Additional file 3: Table S1**).

**Fig. 2.**
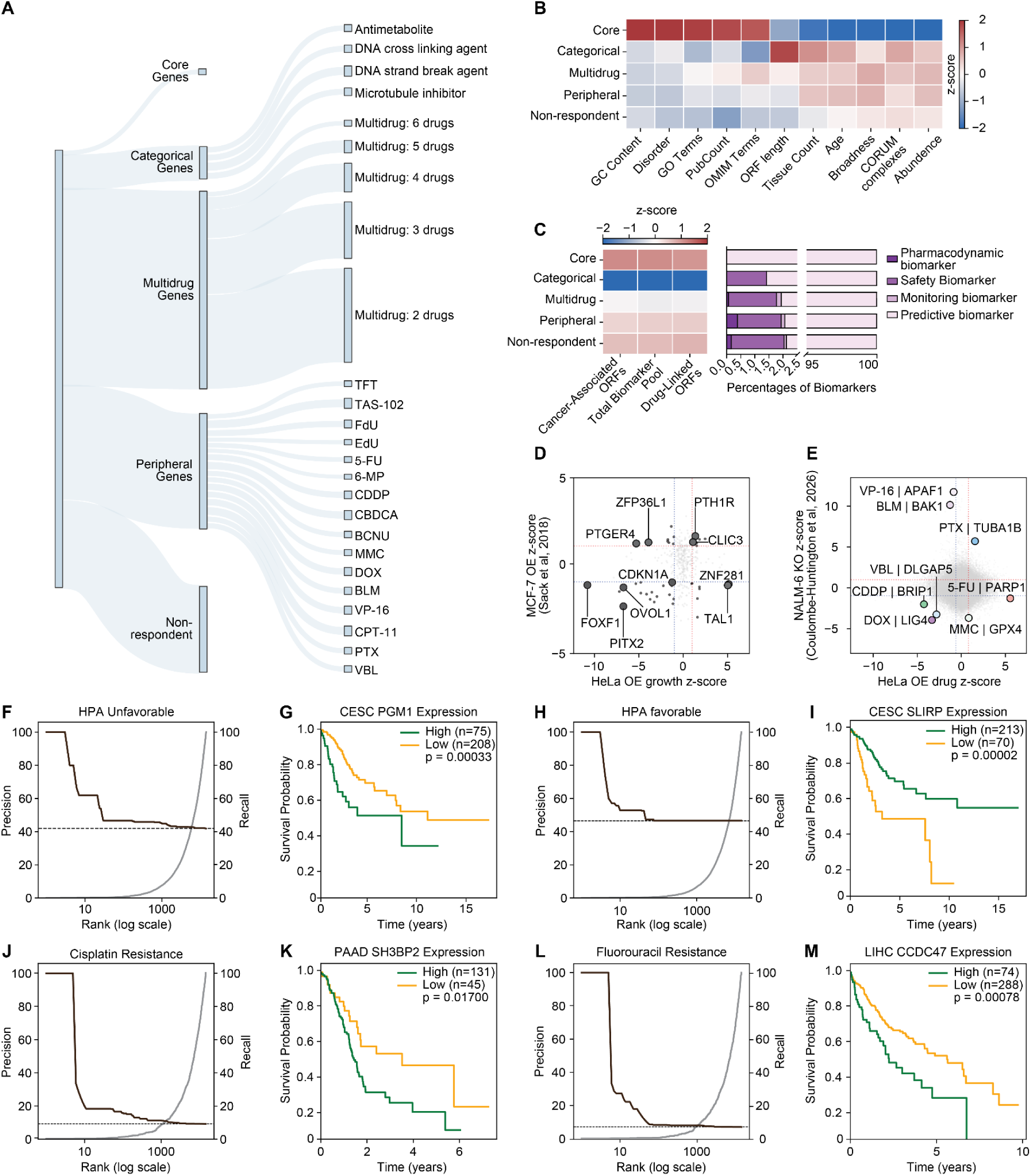
Biological and clinical relevance of the GOF atlas for cancer proliferation and chemotherapy response. **(A)** Sankey diagram categorizing ORF numbers into drug response groups across 16 chemotherapeutic agents. **(B)** Heatmap of *z*-scores for intrinsic ORF characteristics across categories. **(C)** Left: Heatmap showing overlap between ORFs and cancer biomarkers from TheMarker database; Right: Distribution of biomarker types across ORF categories. **(D)** Quadrant scatter plot comparing cell proliferation effects measured by HeLa overexpression (OE) growth *z*-scores against MCF-7 OE data from Sack et al., 2018. **(E)** Quadrant scatter plot mapping drug response interactions by benchmarking HeLa ORFeome drug *z*-scores against NALM-6 knockout (KO) data across 8 chemotherapeutic agents from Coulombe-Huntington J et al. 2026. **(F, G)** Precision-recall at rank *k* curve ranking ORFs by cell proliferation effects, comparing top upregulators with unfavorable prognosis from the HPA dataset **(F)**, paired with the corresponding representative Kaplan-Meier survival curve for an unfavorable prognostic marker in cervical cancer **(G)**. **(H, I)** Precision-recall at rank *k* curve ranking ORFs by cell proliferation effects, comparing top downregulators with favorable prognosis from the HPA dataset **(H)**, paired with the corresponding representative Kaplan-Meier survival curve for a favorable prognostic marker in cervical cancer **(I)**. **(J, K)** Precision-recall at rank *k* curve evaluating cisplatin resistance drivers from the ORFeome screen **(J)**, paired with the corresponding representative Kaplan-Meier survival curve for a resistance-associated candidate linked to reduced survival in pancreatic cancer **(K)**. **(L, M)** Precision-recall at rank *k* curve evaluating 5-FU resistance drivers from the ORFeome screen **(L)**, paired with the corresponding representative Kaplan-Meier survival curve for a resistance-associated candidate linked to reduced survival in liver cancer (LIHC) **(M)**. Survival curve comparisons were performed using the log-rank test.

We next investigated whether the defined ORF categories exhibit distinct intrinsic features underlying their differential drug response profiles. Core ORFs demonstrated the most pronounced divergence across multiple molecular features, showing elevated *z*-scores for GC content, predicted protein disorder [36], Gene Ontology (GO) term enrichment, publication frequency (PubCount), and associations with Mendelian diseases (OMIM terms) [39] (**Fig. 2B, Additional file 3: Table S7**). Conversely, features such as tissue-specific upregulation [38], evolutionary age [35], expression broadness [34], expression abundance [34], and protein complex membership (CORUM database) [40] were markedly reduced in Core ORFs (**Fig. 2B**). Sequence length evaluation revealed that Core ORFs tended to be shorter, whereas Categorical ORFs exhibited longer sequences. These observations suggest that ORFs with lower baseline expression may have greater phenotypic impact when artificially overexpressed in GOF screens, highlighting Core ORFs as a distinct subset of central, multi-drug responsive genes across cell cycle-arrest chemotherapies that gain functional significance upon upregulation.

To evaluate translational relevance, we assessed the overlap of categorized ORFs with cancer-specific biomarkers curated in TheMarker database [41] (**Additional file 3: Tables S8 and S9**). Core ORFs showed elevated *z*-scores across all metrics, including the percentage of cancer-associated ORFs, contribution to the total biomarker pool, and drug-linked biomarker proportions, confirming strong enrichment for clinically validated markers (**Fig. 2C, left**) and a predominance of predictive biomarker traits (**Fig. 2C, right**). Furthermore, cross-platform benchmarking demonstrated strong gene-level agreement between our HeLa screen and an external MCF-7 overexpression dataset (364 ORF overlaps, **Additional file 3: Tables S10 and S11**) [22]. Key proliferation drivers such as *PTH1R* and *CLIC3* displayed concordant growth-promoting phenotypes (**Fig. 2D; top-right quadrant**), whereas growth-inhibitory candidates including *CDKN1A*, *FOXF1*, and *PITX2* consistently suppressed growth across both cell lines (**Fig. 2D; bottom-left quadrant**), while also highlighting lineage-specific proliferation drivers in MCF-7 (*PTGER4, ZFP36L1*) and HeLa (*TAL1, ZNF281*). We next compared the relationship between ORF length and absolute fold-change magnitude (|log_2_FC|) across 1-kb length intervals against published lentivirus-based screening platforms [22,42]. Our platform maintained a consistently higher phenotypic dynamic range across all transcript sizes, including large ORFs between 4 and 9 kb (**Additional file 2: Fig. S2I**). Because plasmid-based Bxb1 recombination bypasses lentiviral packaging and reverse-transcription size constraints, our single-copy integration system minimizes the length-dependent signal attenuation typical of lentiviral screens, enabling robust functional profiling across the full size spectrum of the human proteome.

To determine whether our GOF screening platform recovers conserved target biology and complements LOF paradigms, we benchmarked our drug response profile against published genome-wide knockout (KO) screens in human pre-B acute lymphoblastic leukemia NALM-6 cells [43], across eight overlapping chemotherapeutic agents. To prioritize candidate gene-drug interactions, *z*-scores were independently calculated within each drug treatment from both platforms, and top candidates with the longest distance from the origin were prioritized in each quadrant (**Fig. 2E, Additional file 3: Table S12**). Among these candidates, *GPX4* showed a reciprocal resistance profile under MMC, consistent with its established role in suppressing lipid peroxidation and ferroptosis [71], while core apoptotic machinery such as *BAK1* (BLM) and *APAF1* (VP-16) [71,72] mapped to expected pro-apoptotic sensitivity axes. Similarly, *PARP1* overexpression conferred resistance to 5-FU, reflecting PARP1-mediated DNA damage repair [73], as well as *TUBA1B* (PTX) for resistance [74]. In contrast, candidate interactions in other quadrants, such as *BRIP1* (CDDP) [75], likely reflect complex dosage sensitivity, physiological expression effects, or context-dependent network dynamics. These findings demonstrate that integrating OE and KO profiles can recapitulate established biological relationships and may provide a complementary strategy for identifying previously uncharacterized drug-gene interactions.

We next evaluated whether ORF proliferation regulators could serve as clinical prognostic biomarkers. To test whether ranking ORFs by their baseline proliferation score recovers clinically relevant genes, we benchmarked these functional fitness ranks against patient prognostic indicators curated in the Human Protein Atlas (HPA) [44] and TCGA [47] databases (**Additional file 3: Tables S13 and S14**). Using precision-recall at rank *k* analysis [76], ranking ORFs in descending order of proliferation score yielded high precision for unfavorable prognostic markers (**Fig. 2F**). Illustrating this relationship in cervical squamous cell carcinoma and endocervical adenocarcinoma (CESC), overexpression of Phosphoglucomutase 1 (*PGM1*), a growth upregulator in our screen (rank #798, *z* = 1.85), was strongly associated with reduced overall survival probability (**Fig. 2G**). ORFs representing growth downregulators were enriched for favorable prognostic markers (**Fig. 2H**). In line with this trend, overexpression of SRA Stem-Loop Interacting RNA Binding Protein (*SLIRP*), a growth downregulator in our screen (rank #569, *z* = −1.92), correlated with improved patient outcomes in CESC (**Fig. 2I**). These precision-recall and survival analyses underscore the clinical validity of our baseline functional screening framework, demonstrating its capacity to prioritize prognostic cell proliferation drivers in human cancer.

Finally, we evaluated the ability of our platform to identify established drug resistance mechanisms by ranking drug response ORFs against curated TCGA resistance datasets [45]. Evaluated by the Precision-recall at rank *k* curve, top-ranked drug response ORFs displayed high precision for identifying confirmed resistance drivers for both CDDP (**Fig. 2J**) and 5-FU (**Fig. 2L**). Clinically, overexpression of Src homology 3 domain-binding protein 2 (*SH3BP2*), a top cisplatin drug response ORF, was strongly associated with poor overall survival in pancreatic adenocarcinoma (PAAD; **Fig. 2K**). Similarly, Coiled-coil domain-containing protein 47 (*CCDC47*), identified as a 5-FU drug response ORF, correlated with adverse prognosis in liver hepatocellular carcinoma (LIHC) (**Fig. 2M**), with both candidates displaying robust drug-specific profiles in our screening data (**Additional file 2: Fig. S2J**). These findings validate that the proliferation-based ranking recovers established resistance drivers for both agents, with the top-ranked candidates from each screen also showing prognostic associations in patient cohorts.

### Network analysis of the atlas points to the *BCL-2* family-associated autophagy-endolysosomal network on TAS-102

To investigate how the functional ORF categories organize into broader biological modules across treatments, we constructed a global drug-response similarity network encompassing all 11,595 drug-responsive ORFs connected by 100,347 pairwise-complete correlation edges across the 16-agent profile (**Fig. 3A, Additional file 3: Table S15**). Within this similarity landscape, single-agent-specific (Peripheral) ORFs segregated into 16 discrete peripheral clusters corresponding to their respective treatments, whereas Core (n = 197) and Multidrug (n = 6,556) ORFs occupied the central territory bridging multiple drug classes (**Fig. 3A**). Across all 16 chemotherapeutics, Core ORFs consistently exhibited the highest frequency of resistance-conferring perturbations, reaching 47% of measured Core ORFs under the nucleoside analog TAS-102 (**Additional file 2: Fig. S3A**).

**Fig. 3.**
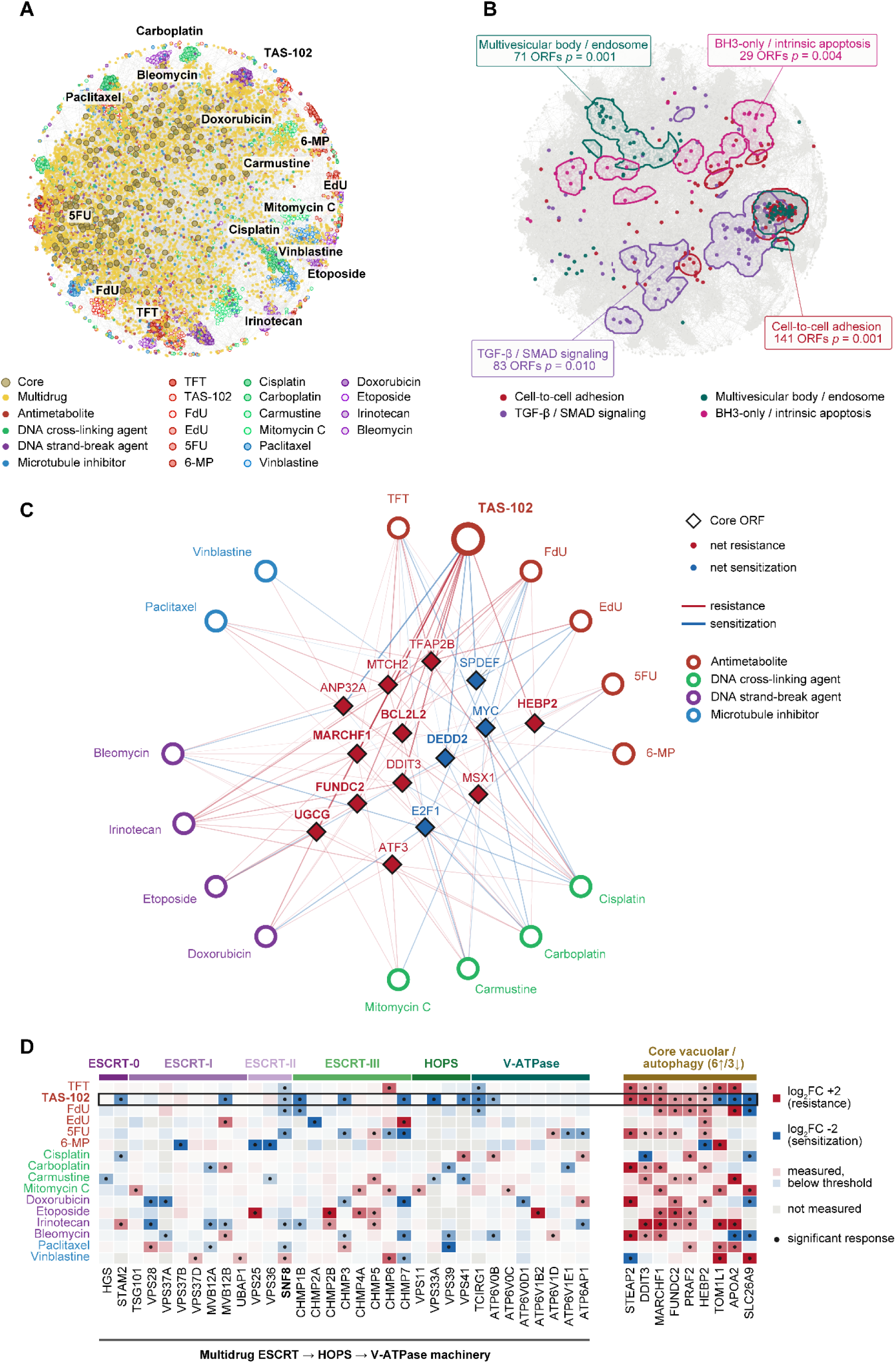
Global drug-response network architecture and hierarchical convergence of the *BCL-2* family-associated autophagy-endolysosomal axis on TAS-102. **(A)** Global ORF-ORF drug-response similarity network connected by pairwise-complete Pearson correlation edges calculated using 16 drug response profiles. Periphery labels indicate clusters formed by single-agent-specific ORFs. **(B)** Functional neighborhood enrichment mapped onto the similarity network layout from (A), where the local neighborhood of each ORF is defined by its top 100 most strongly correlated ORFs across the 16-drug profiles. Colored nodes and density contours highlight four spatially concentrated functional domains: Multivesicular body/endosome, BH3-only/intrinsic apoptosis, Cell-to-cell adhesion, and TGF-β/SMAD signaling. **(C)** Gene-drug bipartite network of the 15 Core ORFs associated with apoptosis, *BCL-2* family [*BCL2L2* (known as *BCL-w*)], and endolysosomal axis (113 gene-drug interactions across the 16 chemotherapeutics). Each Core ORF node is positioned at the response-weighted centroid of its interacting agents and colored by its net response direction across treatments (red, net resistance; blue, net sensitization). Outer circular nodes represent the 16 chemotherapeutic agents colored by mechanism of action, and edges indicate individual significant responses (red lines, resistance; blue lines, sensitization; edges connected to TAS-102 are highlighted). **(D)** Heatmap of log_2_FC drug response profiles across all 16 drugs for the 33 responsive subunits of the endolysosomal cascade spanning ESCRT-0, ESCRT-I, ESCRT-II, ESCRT-III, HOPS, and V-ATPase (left, Multidrug ESCRT → HOPS → V-ATPase machinery; *SNF8* is highlighted in bold as the sole Core ORF), contrasted with TAS-102-responsive Core vacuolar and autophagic ORFs (right, 6 resistance and 3 sensitization hits in TAS-102). Heatmap colors represent log_2_FC direction and magnitude, black dots denote statistically significant responses (|log_2_FC| ≥ 0.5 and integrated *p* < 0.05), and the TAS-102 row is outlined in black (10/10 sensitization across the endolysosomal cascade, sign-test *p* = 0.001; 23/33 subunits directionally sensitized, binomial *p* = 0.0175).

Mapping functional neighborhood enrichment onto this network against size-matched random gene sets revealed localized clustering of ORFs associated with *BCL-2* homology (BH) BH3-only [77] and intrinsic apoptosis, multivesicular body and endosome, cell-to-cell adhesion, and TGF-β/SMAD signaling (**Fig. 3B, Additional file 3: Table S16**). To evaluate these pathways without network-level clique amplification, we systematically tested 14 pre-specified apoptotic, autophagic, and endolysosomal gene sets across all 16 agents and both response directions (**Additional file 2: Fig. S3B**). Across the 842 ORFs annotated to this combined *BCL-2*, autophagy, and lysosome axis, every chemotherapeutic agent drove subsets of axis genes in both resistance and sensitization directions. TAS-102 engaged the largest total number of axis regulators (81 resistance and 62 sensitization hits) and was the only agent to engage all six Core ORFs of the axis (*MARCHF1*, *HEBP2*, *FUNDC2*, *BCL2L2*, *UGCG*, and *DEDD2*; **Additional file 2: Fig. S3C**). Unlike transcription factors, these six Core axis ORFs encode direct mitochondrial, autophagic, and endolysosomal effectors that drive unidirectional drug responses across treatments without altering baseline cell proliferation.

Dissecting this convergence across functional tiers revealed a clear hierarchical division of labor between Core apoptotic regulators and Multidrug endolysosomal complexes (**Fig. 3C–D**). We next constructed a gene-drug bipartite network of the 15 Core ORFs comprising nine apoptotic signaling Core ORFs together with the six Core axis ORFs (113 gene-drug interactions; **Fig. 3C**). Within this network, TAS-102 connected to all six Core axis hubs, including the anti-apoptotic target *BCL2L2*, mitochondrial *HEBP2*, mitophagy receptor *FUNDC2*, endolysosomal E3 ligase *MARCHF1*, and glucosylceramide synthase *UGCG* in the resistance direction, paired with pro-apoptotic *DEDD2* in the sensitization direction (**Fig. 3C**). Consistently, the canonical *BCL-2* family exhibited significant enrichment for TAS-102 resistance (3.63-fold, *p* = 0.0042), which remained robust after controlling for baseline proliferation bias (**Additional file 2: Fig. S3B, D**).

Conversely, consistent with the depletion of multi-subunit complexes among Core ORFs (**Fig. 2B**), the obligate multi-subunit machinery governing MVB biogenesis, endolysosomal fusion, and luminal acidification resided almost exclusively within the Multidrug category. Across the 33 responsive subunits spanning ESCRT-0, ESCRT-I, ESCRT-II, ESCRT-III, HOPS, and the V-ATPase proton pump, the ESCRT-II subunit SNF8 was the sole Core ORF (**Fig. 3D, Additional file 3: Table S17**). Whereas these subunits showed mixed directional responses across other chemotherapeutics, every significant subunit hit under TAS-102 acted exclusively as a sensitizer (10/10 sensitizers versus 0 resisters, sign-test *p* = 0.0010; **Fig. 3D**). These sensitizing hits include ESCRT-0 (*STAM2* with log_2_FC of −1.41), ESCRT-I (*MVB12B*, −1.54), ESCRT-II (*SNF8*, −0.80), ESCRT-III (*CHMP1B*, −1.77; *CHMP3*, −1.59; *CHMP7*, −1.80), HOPS (*VPS33A*, −2.08; *VPS41*, −1.65), and V-ATPase (*TCIRG1*, −1.65; *ATP6V0B*, −0.96). Across the entire gene enrichment grid, sensitization of the 33-subunit ESCRT, HOPS, and V-ATPase cascade under TAS-102 ranked as the top endolysosomal enrichment (4.92-fold with *p* = 1.87 × 10^-5^; **Additional file 2: Fig. S3B**). This enrichment survived proliferation-bias control testing (*p* < 0.0001, median set growth *z* = 0.2; **Additional file 2: Fig. S3D**), alongside broader autophagy sensitization (1.73-fold, *p* = 0.014; **Additional file 2: Fig. S3D**). However, these opposing directional readouts highlight an interpretive boundary of survival-based overexpression screens. Whereas the nine core vacuolar and autophagic ORFs split bidirectionally under TAS-102 (6 resistance versus 3 sensitization hits; **Fig. 3D**), all 10 significant subunits of obligate multi-subunit complexes acted as sensitizers. Because single-subunit overexpression can disrupt complex stoichiometry to produce a dominant-negative, LOF phenotype, this uniform sensitization implies that intact endolysosomal function supports survival under TAS-102, necessitating a secondary organelle-specific screen to directly quantify lysosomal activity.

### Proteome-wide lysosome functional screening maps lysosomal modulators driving drug resistance

Building on the *BCL-2* family and the autophagy-lysosome pathway as candidate mediators of TAS-102 response (**Fig. 3**), we sought to systematically identify lysosomal modulators by integrating the overexpression platform with lysosome functional screening using LysoTracker-based fluorescence-activated cell sorting (FACS; **Fig. 4A**). To isolate intrinsic, gene-driven lysosomal remodeling without confounding secondary cytotoxicity from drug exposure, HeLa cells expressing the ORFeome 9.1 + Δ library were stained with LysoTracker under DMSO conditions. Cells were sorted into populations exhibiting high and low lysosomal activity, defined by the high and low 5% of green fluorescence intensity (**Fig. 4A, Additional file 3: Table S18**), denoted as lysosomal activators and repressors, respectively (|log_2_FC| ≥ 0.5 and integrated *p* < 0.05). Unsupervised UMAP dimensionality reduction demonstrated clear spatial segregation between the high- and low-fluorescence populations relative to presort baselines, with tight co-clustering across independent replicates reflecting distinct phenotypic lysosomal states (**Additional file 2: Fig. S4A**).

**Fig. 4.**
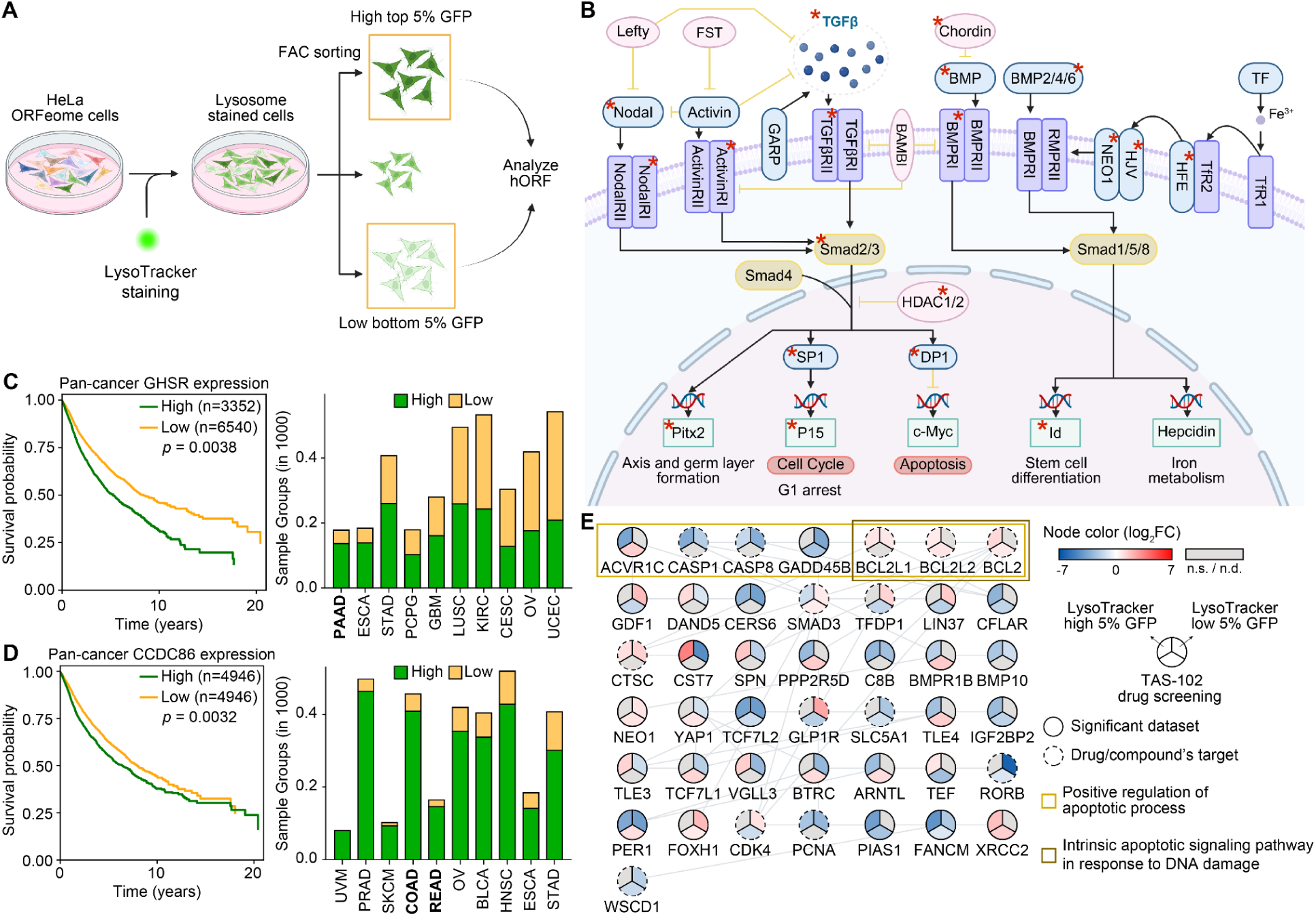
Lysosomal screening and interactomics map the lysosomal axis as a survival hub for target discovery. **(A)** Schematic workflow of the high-throughput LysoTracker-based ORFeome overexpression screen with FACS sorting. HeLa ORFeome cells stained with LysoTracker were subjected to fluorescence-activated cell sorting (FACS) to isolate the top 5% (high fluorescence, lysosomal activation) and bottom 5% (low fluorescence, lysosomal repression) populations for hORF identification. **(B)** Diagram of the TGF-β signaling pathway highlighting lysosomal modulating ORFs (marked with red asterisks) identified in the LysoTracker-based ORFeome screening. **(C–D)** Kaplan-Meier overall survival curves depicting patient survival stratified by **(C)** *GHSR* (detectable [High] vs. undetectable [Low]) and **(D)** *CCDC86* (median split [High vs. Low]) expression across the TCGA PanCancer Atlas, accompanied by stacked bar graphs showing patient cohort distributions across individual cancer studies (denoted by TCGA study abbreviations). **(E)** Functional sub-network interactome mapping lysosomal modulators and TAS-102 screening metrics, where nodes are rendered as pie charts indicating multi-dimensional log_2_FC values [including TAS-102 overexpression response, as well as upper and lower LysoTracker fluorescence populations; color-coded from blue for negative log_2_FC to red for positive log_2_FC; gray denotes not significant (n.s.) or not detected (n.d.)]. Solid node outlines indicate significant screening datasets, and dashed node outlines designate known drug or compound targets. Edges denote STRING functional protein-protein interactions with a confidence score ≥ 0.7, with yellow and brown boxes framing GO enrichment terms. Statistical significance for survival curve comparisons was determined using the cancer-type-stratified log-rank test.

Pathway enrichment analysis of these identified lysosomal modulators demonstrated a mechanistically coherent convergence upon TGF-β signaling networks (**Fig. 4B**), highlighting an upstream regulatory axis governing cellular stress, autophagy, and lysosomal biogenesis. To further evaluate drug-responsive genes associated with lysosomal activity, we intersected our lysosomal modulators with TAS-102-specific response ORFs using an UpSet plot analysis, identifying 109 unique ORFs as lysosomal activators, 78 as lysosomal repressors, and 91 intersecting candidates that link lysosomal reprogramming to TAS-102 responses (**Additional file 2: Fig. S4B**). Among the top-tier lysosomal activators, we prioritized *GHSR* and *CCDC86* as potential novel markers based on three stringent criteria: (i) strong activation magnitude log_2_FC > 1, (ii) functional novelty, and (iii) a pronounced TAS-102 resistance phenotype. In multi-cohort survival analyses across the TCGA PanCancer Atlas (n = 9892) [47], patients with detectable (High, n = 3352) GHSR transcript exhibited significantly shorter overall survival than those with undetectable (Low, n = 6540) GHSR (median 62.8 vs. 96.2 months; stratified log-rank *p* = 0.0038; stratified Cox HR = 1.12, 95% CI 1.04–1.22, *p* = 0.0039; **Fig. 4C, left panel**), with GHSR most frequently detected in pancreatic tumors (PAAD, 77.0%; 1st, **Fig. 4C, right panel**). CCDC86, which was quantifiable across all tumors, ranked among the highest-expressing cohorts in colon (COAD, 89.7% high; 4th) and rectal adenocarcinoma (READ, 89.0% high; 5th) (**Fig. 4D, right panel**), where elevated expression was significantly associated with poorer overall survival after tumor-type stratification (both high/low n = 4946 respectively, stratified log-rank *p* = 0.0032; stratified Cox HR = 1.15, 95% CI 1.05–1.26, *p* = 0.0032; **Fig. 4D, left panel**). Because TAS-102 is approved for previously treated metastatic colorectal cancer and is under clinical investigation in pancreatic cancer [78,79], and induced *BCL-2* protein expression in these models (**Additional file 2: Fig. S4C**), we focused subsequent functional validation on colorectal and pancreatic models.

To integrate these multi-dimensional datasets into a cohesive functional architecture, we constructed a STRING protein-protein interaction (PPI) network mapping lysosomal modulators alongside TAS-102 screening metrics (**Fig. 4E**). Stringent network filtering was applied by requiring a minimum STRING interaction confidence score of 0.7 and selecting nodes shared across at least two out of the three screening datasets (high 5% LysoTracker and low 5% LysoTracker fluorescence, and TAS-102 drug screening response). Individual gene nodes are rendered as multi-segmented pie charts that capture parallel log_2_FC values across all three screening conditions, with dashed outlines explicitly designating known drug or compound targets (**Fig. 4E**). GO enrichment analysis identified two primary functional clusters: positive regulation of the apoptotic process (yellow box) and the intrinsic apoptotic signaling pathway in response to DNA damage (brown box) (**Fig. 4E**). These functional modules prominently feature anti-apoptotic *BCL-2* family members (*BCL2L1*, *BCL2L2*, and *BCL-2*) alongside other pathway members, all explicitly designated for the established BCL-2 family inhibitor ABT-263, which simultaneously targets *BCL-2*, *BCL2L1*, and *BCL2L2* [80,81] (**Fig. 4E**). This functional screening and network interactome isolate the *BCL-2* family as a primary actionable target in colorectal and pancreatic cancers, providing the rationale for combinatorial therapeutic strategies with ABT-263 and TAS-102 aimed at overcoming lysosomal chemoresistance.

### Lysosomal modulation mediates TAS-102 resistance that is synergistically reversed by the BCL-2 family inhibitor ABT-263 *in vitro* and *in vivo*

We next investigated whether pharmacological inhibition of *BCL-2* family members could reverse lysosomal-mediated chemoresistance to TAS-102. Immunoblotting across HeLa, HCT 116, RKO, HT-29, and MIA PaCa-2 cancer cell lines confirmed that TAS-102 exposure robustly induced *BCL-2* protein expression across HCT 116, RKO, and MIA PaCa-2, whereas baseline *BCL-2* levels remained low in HeLa and undetectable in HT-29 cells (**Additional file 2: Fig. S4C**). Assessment of autophagic flux and lysosomal markers revealed that TAS-102 treatment induced notable upregulation of LAMP1 and TFEB, indicating expansion of the lysosomal compartment and activation of master lysosomal transcriptional programs. Concurrently, TAS-102 enhanced autophagic flux, as evidenced by increased conversion of LC3B-I to LC3B-II alongside progressive degradation of p62. Co-treatment with ABT-263 effectively suppressed TAS-102-induced TFEB and LAMP1 upregulation while restoring p62 levels (**Additional file 2: Fig. S4D**), demonstrating that *BCL-2* family protein inhibition attenuates adaptive autophagic and lysosomal responses triggered by TAS-102 stress.

At the cellular level, confocal immunofluorescence confirmed that TAS-102 treatment promoted striking nuclear translocation of TFEB relative to vehicle controls (**Additional file 2: Fig. S4E**). Quantification of normalized nuclear TFEB fluorescence intensity demonstrated a significant increase upon TAS-102 exposure, which was blunted by ABT-263 single treatment and further suppressed under combination therapy (**Fig. 5A**). To evaluate lysosomal proteolytic activity, we performed functional assays using DQ-BSA, where cleavage of self-quenched fluorophores inside active lysosomes yields bright red fluorescence (**Additional file 2: Fig. S4F**). TAS-102 treatment substantially enhanced proteolysis, whereas co-treatment with ABT-263 markedly reduced lysosomal hydrolytic activity (**Fig. 5B**, **Additional file 2: Fig. S4F**). Assessment of lysosomal pH revealed that TAS-102 promoted lysosomal acidification (pH 5.15 to 4.80), whereas the combination reversed this shift beyond baseline (pH 5.30), indicating a net loss of lysosomal acidification rather than a simple return to the untreated state (**Fig. 5C**). ABT-263 alone produced a small but statistically significant acidic shift of uncertain biological relevance, consistent with a mild stress response in the absence of the TAS-102-driven TFEB program. Consistent with this, co-treatment of colorectal cancer cells with TAS-102 and the vacuolar H⁺-ATPase inhibitor BafA1 significantly impaired cell proliferation compared with single-agent treatments (**Fig. 5D**), indicating that functional lysosomal acidification contributes to TAS-102 survival and that the combination phenocopies V-ATPase inhibition.

**Fig. 5.**
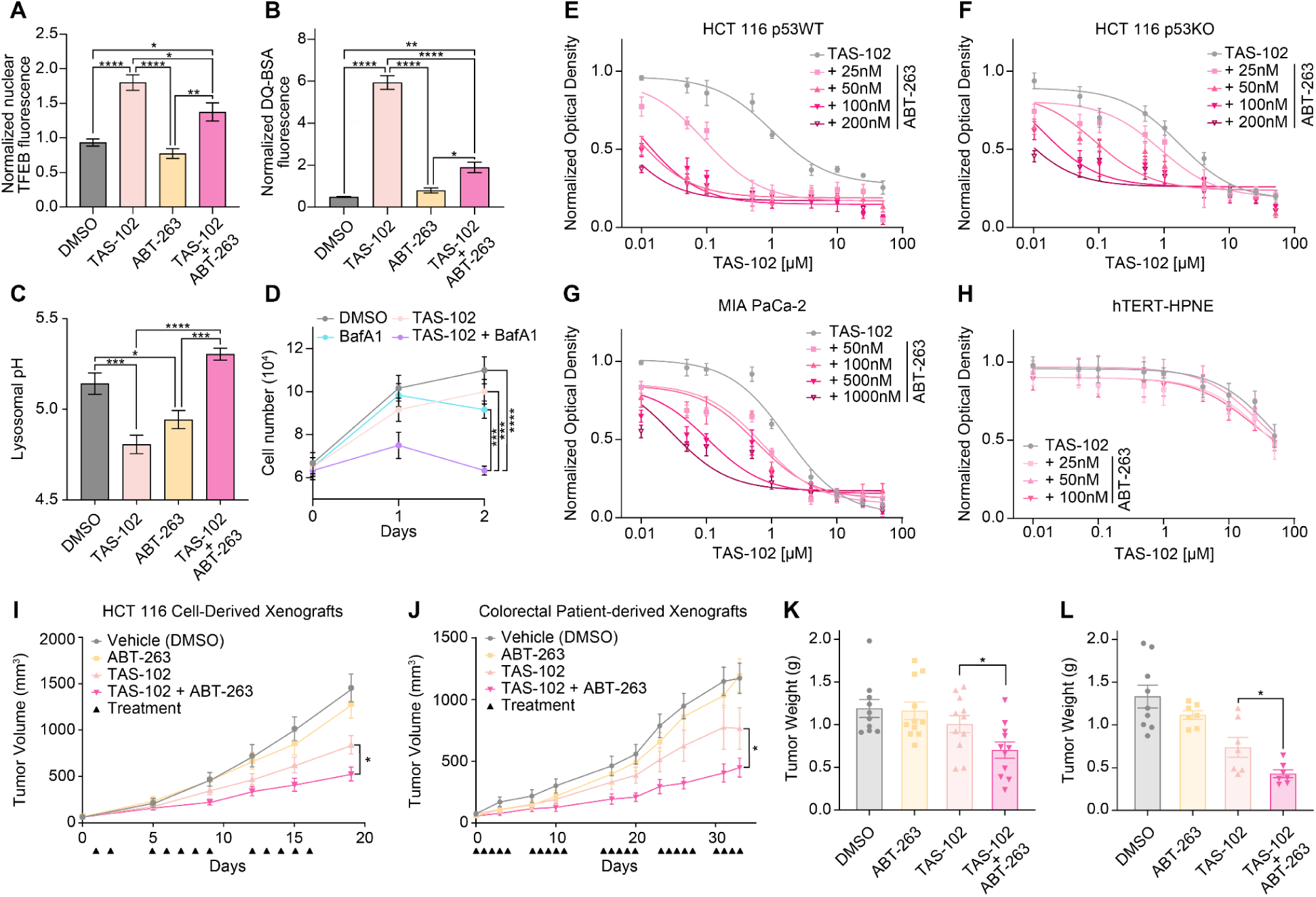
Lysosomal modulation mediates TAS-102 resistance that is synergistically reversed by the BCL-2 family inhibitor ABT-263. **(A)** Quantification of normalized nuclear TFEB fluorescence across vehicle (DMSO), TAS-102, ABT-263, and combination (TAS-102 + ABT-263) treatments in HCT 116. **(B)** Quantification of normalized DQ-BSA fluorescence intensity assessing lysosomal proteolysis in HCT 116 cells. **(C)** Measurement of lysosomal pH alterations across treatment conditions. **(D)** Cell proliferation assay of HCT 116 treated with vehicle, TAS-102, BafA1, or combination treatment over 2 days. **(E–G)** Dose-response proliferation curves evaluating synergistic growth inhibition of TAS-102 across increasing concentrations of ABT-263 in **(E)** HCT 116 *p53* WT, **(F)** HCT 116 *p53* KO, and **(G)** MIA PaCa-2 pancreatic cancer cells. **(H)** Cytotoxicity assay in non-transformed hTERT-HPNE human pancreatic ductal epithelial cells. **(I–J)** Longitudinal tumor volume (mm^3^) progression in **(I)** HCT 116 CDX and **(J)** colorectal PDX mouse models following treatment with Vehicle (DMSO), ABT-263, TAS-102, or combination therapy. Arrowheads denote treatment administration days. **(K–L)** Endpoint tumor weight comparisons for **(K)** HCT 116 CDX and **(L)** colorectal PDX models. *In vitro* data (**A–H**) are presented as mean ± standard deviation (SD). In vivo tumor growth and weight data (**I–L**) are presented as mean ± SEM. Statistical significance for panels **A–D** and **I–J** was evaluated using two-way ANOVA with Tukey’s multiple comparisons test (with TAS-102 and ABT-263 as factors for panels **A–C**, and treatment and time as factors for panels **D** and **I–J**). Statistical significance for tumor weights (**K–L**) was evaluated using Welch’s two-tailed *t*-test; * *p* < 0.05, ** *p* < 0.01, *** *p* < 0.001, **** *p* < 0.0001.

To establish the broader therapeutic synergy between TAS-102 and ABT-263 across colorectal and pancreatic cancers, we performed dose-response proliferation assays across increasing concentrations of ABT-263. Although literature-curated STRING PPI networks link the TAS-102 target TYMS to the *BCL-2* family and autophagy machinery (*BECN1*, *ATG14*) canonically through *TP53* (**Additional file 2: Fig. S5A**), our primary screen captured this survival axis in *p53*-compromised HeLa cells, suggesting that TAS-102 can trigger adaptive *BCL-2* family and lysosomal signaling in *p53*-compromised settings. Using a sequential regimen of 24 h TAS-102 pretreatment followed by ABT-263 exposure (**Additional file 2: Fig. S5B**), we observed robust dose-dependent growth inhibition across isogenic HCT 116 *p53* WT (Fig. 5E) and HCT 116 *p53* KO (**Fig. 5F**) models, as well as *p53*-mutant MIA PaCa-2 (**Fig. 5G**). Combinatorial synergy was quantitatively confirmed by Bliss [48], HSA [49], and ZIP [50] synergy modeling (**Additional file 3: Tables S19 and S20**). Baseline single-agent sensitivity to ABT-263 across additional colorectal cancer models (HCT 116, RKO *p53* WT and KO, and HT-29) confirmed concentration-dependent responses across all lines (**Additional file 2: Fig. S5C**). Higher, micromolar concentrations of ABT-263 (1–5 µM) were required to elicit combinatorial growth inhibition in RKO *p53* WT and KO models (**Additional file 2: Fig. S5D–E**). Across the tested lines, comparable combinatorial responses were observed between *p53* WT and *p53* KO isogenic backgrounds, consistent with the *p53*-independent activity reported for ABT-263 [81]. Importantly, combination treatment showed minimal cytotoxicity and lacked synergistic activity in non-transformed hTERT-HPNE human pancreatic ductal epithelial cells (**Fig. 5H**), establishing a favorable *in vitro* therapeutic window [80].

We next translated these findings in vivo using cell line-derived xenograft (CDX) and patient-derived xenograft (PDX) mouse models using a sequential dosing schedule in which TAS-102 was administered 4 h before ABT-263 to allow adaptive *BCL-2* family and lysosomal priming before *BCL-2* family blockade. Longitudinal tumor volume tracking in HCT 116 CDX models revealed that while single-agent TAS-102 or ABT-263 provided moderate growth delay, combination therapy significantly suppressed tumor progression (**Fig. 5I**). Superior anti-tumor activity was mirrored in the colorectal PDX model, where combination therapy resulted in marked tumor growth inhibition relative to control or single-agent groups (**Fig. 5J**). Endpoint tumor mass evaluation confirmed significant reductions in tumor weight in both HCT 116 CDX (**Fig. 5K**) and colorectal PDX models (**Fig. 5L**). The combination regimen was well tolerated with no overt systemic toxicity *in vivo*, as stable body weight profiles were maintained throughout the treatment schedule in both CDX and PDX models (**Additional file 2: Fig. S5F, J**). Complete blood count (CBC) analyses revealed no significant differences across treatment groups for white blood cells or red blood cells in either model (**Additional file 2: Fig. S5G–H, K–L**). As expected for *BCL-2* family targeted therapies, no hematologic abnormalities were observed apart from a modest, transient reduction in platelet counts in the combination group relative to vehicle controls in PDX and to ABT-263 in CDX (**Additional file 2: Fig. S5I, M**). Together, these results demonstrate that lysosomal adaptation contributes to TAS-102 chemoresistance and that combinatorial targeting with ABT-263 attenuates this lysosomal response, synergistically restores sensitivity and suppresses tumor growth *in vitro* and *in vivo*.

## Discussion

In this study, we established a standardized, human proteome-wide GOF atlas that systematically maps cellular survival and sensitization dynamics across 16 chemotherapeutics spanning four major cytotoxic classes. Our platform ensures quantitative uniformity across perturbations and directly complements LOF (KO) screens and conventional transcriptomics. Stratified modeling of over 230,000 gene-drug interactions revealed a hierarchical organization of drug response drivers, wherein a distinct subset of ORF categories governs broad multidrug resistance and is highly enriched for clinically validated predictive and prognostic biomarkers. We uncovered that diverse chemotherapeutic stresses, most prominently the nucleoside analog TAS-102, converge upon a central survival hub orchestrated by the *BCL-2* family-associated autophagy-endolysosome axis. TAS-102 triggered *BCL-2* upregulation, nuclear translocation of TFEB, lysosomal acidification, and enhanced autophagic proteolysis, which were attenuated alongside synergistic tumor suppression by co-treatment with the BCL-2 family inhibitor ABT-263 in colorectal CDX and PDX models.

Our network and functional interactome analyses identified multi-paralog anti-apoptotic modules comprising *BCL-2*, *BCL2L1*, and *BCL2L2* as potential TAS-102 resistance modules, providing an a priori rationale for selecting the broad-spectrum inhibitor ABT-263 over *BCL-2* selective agents (**Fig. 3C**, **Fig. 4E**). Here, whereas *BCL-2* and *BCL2L2* crossed the primary survival cutoff under TAS-102, *BCL2L1* scored directly as a significant lysosomal modulator in the lysosomal screen (**Fig. 4E**) and exhibited a concordant, sub-threshold positive shift (log_2_FC = 0.25) across antimetabolites under single-gene overexpression. Within this family, pro-apoptotic members (BAX, BBC3, PMAIP1) also scored in the resistance direction. Because single-gene overexpression does not by itself license productive BAX oligomerization, and because BAX–BAK activation can be diverted toward mtDNA efflux rather than canonical apoptosis in apoptosome-deficient settings [82], our interpretation rests on the anti-apoptotic members validated at the protein level (**Additional file 2: Fig. S4C–D**). Consistent with functional lymphoma modeling and recent high-density genome-wide CRISPRa screens showing that selective *BCL-2* inhibition with venetoclax is readily bypassed via compensatory transcriptional upregulation of *BCL2L1* or *MCL1* [83], simultaneous blockade of multiple *BCL-2* family paralogs using the broad-spectrum inhibitor ABT-263 was required to attenuate adaptive lysosomal survival signaling alongside canonical anti-apoptotic blockade under TAS-102 stress. This paralog dependency also explains the distinct sensitivity profiles observed across our colorectal cancer models. While HCT 116 and RKO cells both share microsatellite instability-high (MSI-H) status, they diverged markedly in their responsiveness to combination therapy across both *p53* WT and KO backgrounds. Beyond morphological differences between epithelial HCT 116 and mesenchymal-like RKO cells, specific somatic driver mutations such as *KRAS* and *BRAF* may govern this therapeutic window. *KRAS*-mutant models (HCT 116 and MIA PaCa-2) exhibit potent synergy at nanomolar ABT-263 concentrations (25–200 nM in HCT 116 and 50–1,000 nM in MIA PaCa-2) due to *KRAS*-driven reliance on the *BCL-2* family and lysosomal survival axis (**Fig. 5E–G**). Conversely, *BRAF* V600E-mutant RKO cells required micromolar ABT-263 concentrations (1–5 µM) to achieve growth suppression (**Additional file 2: Fig. S5D–E**), likely because *BRAF* V600E promotes apoptosis evasion through downstream upregulation of *MCL-1*, an anti-apoptotic paralog not targeted by ABT-263, thereby reducing cellular dependency on *BCL-2* family modulation [84]. Beyond the *BCL-2* family, our integrated screens also uncovered signaling pathways governing chemoresistance and lysosomal biogenesis, including TGF-β signaling, alongside novel resistance-associated lysosomal activators such as *GHSR* and *CCDC86* (**Fig. 4B–D, Additional file 2: Fig. S4B**). Further mechanistic investigation of these identified drivers will help elucidate their precise roles in lysosomal reprogramming and evaluate their potential as actionable therapeutic targets.

From a clinical perspective, although TAS-102 monotherapy is well tolerated and provides meaningful disease stabilization (58%) in heavily pretreated metastatic colorectal cancer patients, objective tumor regression remains exceptionally rare, with a 0% objective response rate and median progression-free survival of ∼2.1 months [78]. Pairing TAS-102 with a BCL-2 family inhibitor offers a rational strategy to convert this cytostatic disease stabilization into active tumor regression. In our *in vivo* CDX and PDX studies, combination therapy significantly suppressed tumor growth while maintaining stable body weights and normal leukocyte and erythrocyte counts, accompanied by a modest, transient reduction in platelet counts (**Additional file 2: Fig. S5F–M**) that reflects the known on-target inhibition of *BCL2L1* in circulating platelets [85]. Several clinical frameworks support the translational feasibility and safety of this regimen. First, in real-world clinical practice, TAS-102 monotherapy is administered on an intermittent 28-day schedule incorporating a 14-day rest period and induces negligible severe thrombocytopenia (0% grade 3–4), minimizing overlapping megakaryocytic toxicity [78]. Second, clinical trials of navitoclax have established that a 7-day low-dose lead-in titration (150 mg) or weight-adjusted low-dose combination schedules effectively mitigate severe thrombocytopenia in patients [85]. Third, next-generation platelet-sparing BCL-XL PROTAC degraders such as DT2216, which recruit the VHL E3 ligase that is minimally expressed in human platelets, have demonstrated clinical safety, rapid platelet recovery without bleeding, and strong synergy with conventional chemotherapies in phase 1 solid tumor trials [86].

Despite the breadth of our resource, several methodological limitations should be considered. First, regarding our gene overexpression architecture, Bxb1 recombination bypasses lentiviral packaging limits and preserves stronger functional sensitivity for longer ORFs (**Additional file 2: Fig. S2I**) compared to lentiviral screens. However, initial cellular integration efficiency still exhibited an inverse correlation with ORF length (*r* = −0.477; **Additional file 2: Fig. S1J**), potentially reflecting translation and selection bias introduced by the downstream IRES-mCherry reporter cassette for longer transcripts. Furthermore, insufficient ORF recovery in one replicate restricted 6-MP, CPT-11, and VBL to duplicate rather than triplicate profiling (45 total drug-treated samples across 16 agents). In addition, EdU exhibited a lower total number of detected ORFs compared to other treatments, which reduced statistical power in these conditions and necessitated prioritizing integrated *p*-values over FDR *q*-values for pan-atlas comparisons (**Additional file 2: Fig. S2H**). Nevertheless, we report q-values across all 232,096 gene-drug pairs (**Additional file 3: Table S1**), and our primary mechanistic hits under TAS-102, including *BCL-2* (*q* = 0.151), *BCL2L2* (*q* = 0.159), and 5/6 Core axis regulators (*q* = 0.115 to 0.167), satisfy an exploratory false-discovery threshold of *q*-values < 0.20. The 33-subunit ESCRT, HOPS, and V-ATPase cascade maintains significant directional sensitization even without significance thresholding (23/33 subunits with negative log_2_FC, one-sided binomial *p* = 0.0175; **Additional file 3: Table S17**). Likewise, HSR screening was only conducted in three representative chemotherapeutics (DOX, CDDP, and TFT) rather than all 16 agents. Second, because the hORFeome library comprises canonical protein-coding sequences expressed as single copies under a constitutive EF1α promoter at the AAVS1 locus, our platform does not capture non-coding RNAs, alternative splice isoforms, or variable focal copy-number amplifications observed in clinical tumors. It also does not capture phenotypes requiring stoichiometric co-expression of multi-subunit protein complexes, consistent with the lower representation of CORUM complex members among Core ORFs (**Fig. 2B**). Third, while pooled scRNA-seq bridged our phenotypic hits with downstream transcriptional modules (**Fig. 1F–G**), profiling the full 18,388 ORF library in a pooled format resulted in modest per-ORF cell coverage (∼2x detection depth), limiting the resolution for de novo reconstruction of individual gene regulatory networks. Coupling targeted sub-library retesting (such as our >1,000x HSR framework) with direct Perturb-seq capture [87,88] will substantially deepen single-cell resolution to uncover rare resistant subpopulations. Fourth, our primary screens were conducted in a single proof-of-principle HeLa cell line at a fixed endpoint, which does not fully capture lineage-specific dependencies, dynamic resistance evolution, or tumor microenvironmental and immune interactions. Integrating our gene overexpression architecture into longitudinal time-course screens, *in vivo* models [89], patient-derived organoids [90], and expanded multi-patient PDX cohorts across diverse cancer lineages [91] will distinguish early survival drivers from long-term adaptive mechanisms and further expand the translational depth of our framework.

Beyond our proof-of-concept validation of the TAS-102 and ABT-263 combination, this proteome-wide GOF atlas provides an immediate engine for systematic drug repurposing and rational combination design [92]. Because clinical inhibitors intrinsically target overactive or upregulated proteins, the resistance and sensitization drivers mapped across our atlas represent actionable vulnerabilities that can be matched with existing targeted agents to sensitize tumors to standard-of-care chemotherapies, substantially bypassing the time and cost of *de novo* drug discovery. By complementing LOF reference maps with a standardized GOF dimension, our open-access resource establishes a foundational systems-genomics blueprint to decode therapeutic resistance and rationally reprogram drug responses in cancer.

## Conclusions

In summary, our study establishes a standardized, open-access human proteome-wide gain-of-function atlas that delineates the genetic architecture of chemotherapy response across 16 cytotoxic agents. By integrating single-copy ORF perturbation screening with single-cell and bulk transcriptomics, clinical survival cohorts, and organelle-level functional profiling, we identified clinically relevant resistance-associated and prognostic biomarkers and uncovered the *BCL-2* family-associated autophagy-endolysosomal network as a central, targetable survival hub under TAS-102 therapy. Preclinical validation in colorectal and pancreatic cancer models in vitro and in colorectal cell line- and patient-derived xenografts in vivo demonstrated that pairing TAS-102 with the BCL-2 family inhibitor ABT-263 attenuates TAS-102-induced lysosomal reprogramming and synergistically overcomes drug resistance. Altogether, this systems-genomics resource provides a translational framework to accelerate biomarker discovery and guide the rational design of combination therapies for refractory cancers.

## List of abbreviations

5-FU: Fluorouracil (5-fluorouracil)
6-MP: 6-Mercaptopurine
AAVS1: Adeno-associated virus integration site 1
ANOVA: Analysis of variance
ATCC: American Type Culture Collection
BafA1: Bafilomycin A1
BCNU: Carmustine [1,3-bis(2-chloroethyl)-1-nitrosourea]
BH: BCL-2 Homology
BLCA: Bladder urothelial carcinoma
BLM: Bleomycin
BOGO: Bxb1-landing pad human ORFeome-integrated system for a proteome-wide Gene Overexpression
BSA: Bovine serum albumin
CBC: Complete blood count
CBDCA: Carboplatin
CCSB: Center for Cancer Systems Biology
CDDP: Cisplatin
CDX: Cell line-derived xenograft
CESC: Cervical squamous cell carcinoma and endocervical adenocarcinoma
CI: Confidence interval
COAD: Colon adenocarcinoma
CORUM: Comprehensive Resource of Mammalian Protein Complexes
CPM: Counts per million
CPT-11: Irinotecan
CRC: Colorectal cancer
CRISPRa: Clustered regularly interspaced short palindromic repeats activation
DAVID: Database for Annotation, Visualization and Integrated Discovery
DEG: Differentially expressed gene
DMSO: Dimethyl sulfoxide
DOX: Doxorubicin
DPBS: Dulbecco’s phosphate-buffered saline
DQ-BSA: Dye-quenched bovine serum albumin
EdU: 5-Ethynyl-2′-deoxyuridine
EF1α: Elongation factor 1-alpha
EGFP: Enhanced green fluorescent protein
ESCA: Esophageal carcinoma
ESCRT: Endosomal sorting complexes required for transport
FACS: Fluorescence-activated cell sorting
FDR: False discovery rate
FdU: Floxuridine (5-fluoro-2′-deoxyuridine)
GBM: Glioblastoma multiforme
gDNA: Genomic DNA
GFP: Green fluorescent protein
GO: Gene Ontology
GOF: Gain-of-function
GTEx: Genotype-Tissue Expression
HNSC: Head and neck squamous cell carcinoma
HOPS: Homotypic fusion and vacuole protein sorting
HPA: Human Protein Atlas
HR: Hazard ratio
HSA: Highest Single Agent
HSR: High-saturation retesting
IACUC: Institutional Animal Care and Use Committee
IC_50_: Half-maximal inhibitory concentration
IRB: Institutional Review Board
IRES: Internal ribosome entry site
KEGG: Kyoto Encyclopedia of Genes and Genomes
KIRC: Kidney renal clear cell carcinoma
KO: Knockout
LIHC: Liver hepatocellular carcinoma
LOF: Loss-of-function
log_2_FC: Log_2_ fold change
LUSC: Lung squamous cell carcinoma
MAPQ: Mapping quality
MMC: Mitomycin C
MSI-H: Microsatellite instability-high
MVB: Multivesicular body
NCBI: National Center for Biotechnology Information
OE: Overexpression
OLS: Ordinary least squares
OMIM: Online Mendelian Inheritance in Man
ORF: Open reading frame
OV: Ovarian serous cystadenocarcinoma
PAAD: Pancreatic adenocarcinoma
PCA: Principal component analysis
PCPG: Pheochromocytoma and paraganglioma
PCR: Polymerase chain reaction
PDX: Patient-derived xenograft
PPI: Protein-protein interaction
PRAD: Prostate adenocarcinoma
PROTAC: Proteolysis-targeting chimera
PTX: Paclitaxel
PVDF: Polyvinylidene difluoride
READ: Rectal adenocarcinoma
RPCCC: Roswell Park Comprehensive Cancer Center
SAFE: Spatial Analysis of Functional Enrichment
scRNA-seq: Single-cell RNA sequencing
SD: Standard deviation
SDS-PAGE: Sodium dodecyl sulfate–polyacrylamide gel electrophoresis
SEM: Standard error of the mean
SKCM: Skin cutaneous melanoma
SRA: Sequence Read Archive
STAD: Stomach adenocarcinoma
STRING: Search Tool for the Retrieval of Interacting Genes/Proteins
SVD: Singular value decomposition
TAS-102: Trifluridine/tipiracil
TCGA: The Cancer Genome Atlas
TFT: Trifluridine
TGF-β: Transforming growth factor beta
TMM: Trimmed mean of M-values
TTD: Therapeutic Target Database
UCEC: Uterine corpus endometrial carcinoma
UMAP: Uniform Manifold Approximation and Projection
UVM: Uveal melanoma
V-ATPase: Vacuolar-type H+-ATPase
VBL: Vinblastine
VHL: von Hippel–Lindau
VP-16: Etoposide
WT: Wild-type
ZIP: Zero Interaction Potency

## Supplementary Information

**Additional file 1.**

- **File format:** Portable Document Format (.pdf)
- **Title of data:** Supplementary Note and Supplementary Methods.
- **Description of data:** Detailed quality control and longitudinal baseline characterization of the engineered HeLa ORFeome pool (Supplementary Note), followed by comprehensive experimental protocols, chemotherapeutic IC_50_ tables, nested PCR primer sequences, antibody specifications, bioinformatic analysis parameters, and supplementary references (Supplementary Methods).

## Additional file 2

- **File format:** Portable Document Format (.pdf)
- **Title of data:** Supplementary Figures S1–S5.
- **Description of data: Fig. S1.** Technical validation and quality control of the proteome-wide GOF screening platform. **Fig. S2.** Methodological workflow, distribution modeling, and functional characterization of screened ORFs. **Fig. S3.** Chemotherapy response and ORF GO enrichment dynamics in the *BCL-2* family-associated autophagy-endolysosomal axis. **Fig. S4.** Characterization of lysosomal modulation. **Fig. S5.** Synergistic efficacy and safety profile of TAS-102 and ABT-263 combination.

## Additional file 3

- **File format:** Microsoft Excel spreadsheet (.xlsx)
- **Title of data:** Supplementary Tables S1–S20.
- **Description of data: Summary.** Overview and column descriptions for Supplementary Tables S1–S20. **Table S1.** Proteome-wide GOF chemotherapy screen matrix across 14,506 quality-filtered ORFs and 16 chemotherapeutic agents. **Table S2.** Longitudinal baseline sequencing read counts normalized to counts per million (CPM). **Table S3.** High-saturation retesting (HSR) sub-library screen results across 2,217 ORFs. **Table S4.** Doxorubicin bulk mRNA transcriptomic analysis in wild-type HeLa cells. **Table S5.** Pooled single-cell RNA-seq differential expression profiles in HeLa ORFeome cells. **Table S6.** Pathway enrichment analysis of drug-responsive ORFs across 16 chemotherapeutics. **Table S7.** Intrinsic molecular and evolutionary gene properties across 14,506 quality-filtered ORFs. **Table S8.** Multi-tiered clinical biomarker overlap summary across ORF categories. **Table S9.** Clinical cancer biomarker annotations from TheMarker database. **Table S10.** Baseline cell proliferation *z*-scores across 14,506 quality-filtered ORFs. **Table S11.** Cross-platform benchmarking datasets comparing HeLa OE against MCF-7 OE proliferation. **Table S12.** Cross-modality benchmarking comparing HeLa OE and NALM-6 CRISPR-Cas9 KO drug responses. **Table S13.** HPA prognostic and TCGA drug-resistance datasets used for precision–recall benchmarking. **Table S14.** Patient survival analysis datasets across HPA/TCGA cohorts (CESC, PAAD, and LIHC). **Table S15.** Global drug-response similarity network coordinates for 11,595 drug-responsive ORFs. **Table S16.** SAFE neighborhood enrichment domains mapped onto the global drug-response similarity network. **Table S17.** Endolysosomal cascade and Core vacuolar/autophagic ORF drug-response profiles. **Table S18.** Proteome-wide LysoTracker Green FACS screening dataset. **Table S19.** Combination drug synergy score summary (Bliss, HSA, and ZIP) for TAS-102 plus ABT-263. **Table S20.** Dosage-pairing cell viability and point-wise synergy matrices across tested cell lines.

## Declarations

### Ethics approval and consent to participate

Animal studies were conducted following the protocol and guidelines approved by the Institutional Animal Care and Use Committee (IACUC) at Roswell Park Comprehensive Cancer Center. The colorectal PDX model (CRC PDX-03-26-4p) was established from a de-identified patient tumor specimen originally collected with written informed consent under a protocol approved by the Institutional Review Board (IRB) at RPCCC. No new human participants were recruited for this study. Animal experiments are reported in accordance with the ARRIVE 2.0 guidelines.

## Consent for publication

Not applicable.

## Availability of data and materials

The sequencing data generated in this study have been deposited in the NCBI database under BioProject SRA accession number PRJNA1353565. The record remains in private status and can be reviewed through the secure link https://dataview.ncbi.nlm.nih.gov/object/PRJNA1353565?reviewer=dc3jinggo7a0jli08lpi00boks until release. The public release of data for all sequencing datasets is set for Oct 27th, 2029, or upon publication, whichever occurs first.

The code and ORFeome reference file used to perform the analyses reported in this study are available on GitHub at the following URL: https://github.com/Harry77j/BOGO_analysis and archived on Zenodo at the following URL: https://doi.org/10.5281/zenodo.23964204 for access. All other data generated or analyzed during this study are included in this published article and its supplementary information files. Plasmids and engineered HeLa landing-pad cell lines generated in this study are available from the corresponding authors upon reasonable request.

## Competing interests

**M.M.A.**, **A.V.B.**, and **D.-K.K.** are inventors on a patent (EP4709390A1) relating to this study [93]. K.Y. and K.K.-H.F. are employees of Illumina Inc. The remaining authors declare that they have no competing interests.

## Funding

This research was supported by Aune Foundation-Endowed Chair Award via Montreal General Hospital Foundation, and the Research Institute of the McGill University Health Centre (**D.-K.K.**), NIH R21CA259719 and DoD BC220542 (**A.V.B.**), National Research Foundation (NRF) funded by the Korean government (NRF-2021M3A9I4024447, **J.-E.P.**; 2022R1A2C2010940 and RS-2025-02303891 (Ministry of Science and ICT(MSIT)), **H.K.K.**) and Chungbuk National University Glocal30 project 2025 (**J.-H.C.**).

## Authors’ contributions

**K.B.J.** and **D.-K.K.** conceptualized and designed the study. **K.B.J.** performed the *in vitro* experiments, processed the data, analyzed the results, drafted the original manuscript, managed the project, and prepared the figures. **Y.G.** and **M.M.A.** performed *in vitro* and *in vivo* experiments. **D.-E.K.** conducted *in vitro* experiments and molecular assays. **Y.K.** processed data and analyzed results. **H.Y.K.** and **J.-S.K.** carried out data curation, figure editing, and critical manuscript review. **S.S.** assisted with data processing and manuscript editing. **K.Y.** provided critical resources and contributed ideas for experimental optimization. **A.G.C.**, **N.S., F.P.R.**, **K.A.M.**, **D.E.H.**, **J.-H.C.**, **K.K.-H.F.**, and **H.L.** contributed to conceptualization, provided resources, experimental advice, manuscript editing, and technical support. **J.-E.P.**, **H.K.K.**, **A.V.B.**, and **D.-K.K.** supervised the project, provided conceptual guidance, and oversaw project administration. All authors read and approved the final manuscript.

## Acknowledgements

We thank Dr. Bert Vogelstein (Johns Hopkins University) and Dr. Alessandro Carugo (MD

Anderson Cancer Center) for generously providing the isogenic HCT 116 and RKO cell lines, respectively. We also thank the Flow & Immune Analysis Shared Resource (FIASR) and the Genomics Shared Resource (GSR) at Roswell Park Comprehensive Cancer Center for technical assistance with flow cytometry sorting and high-throughput sequencing.

---

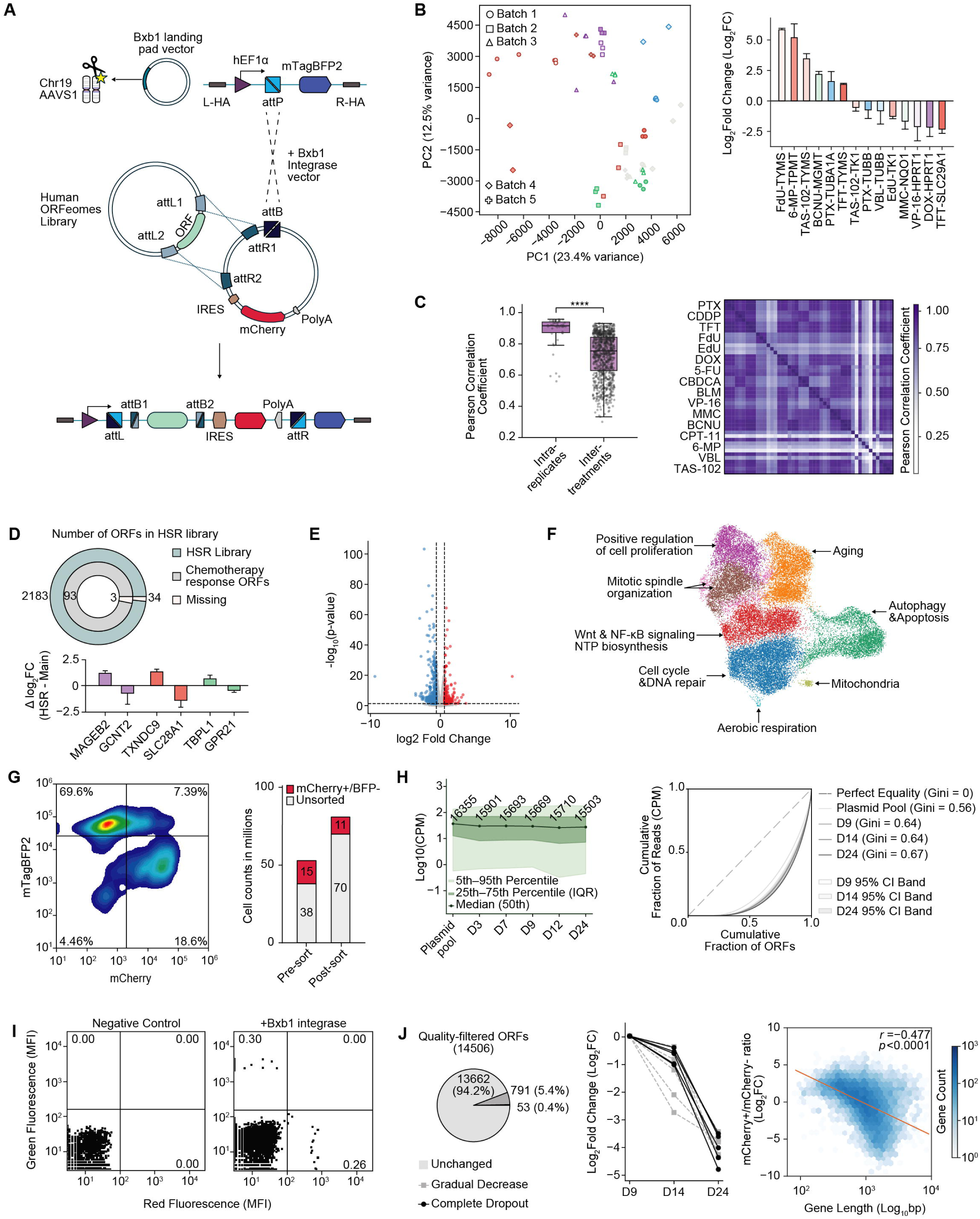

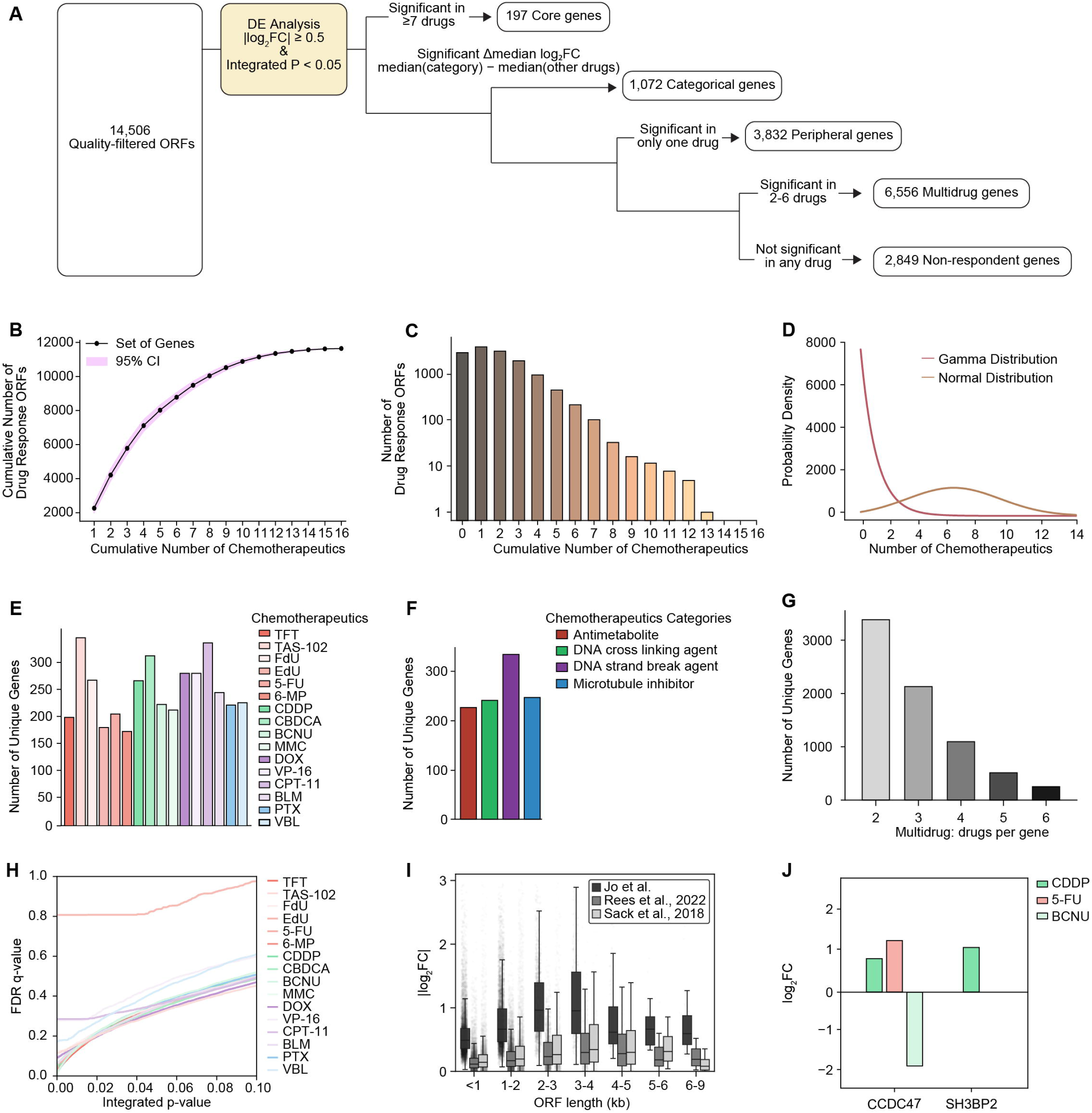

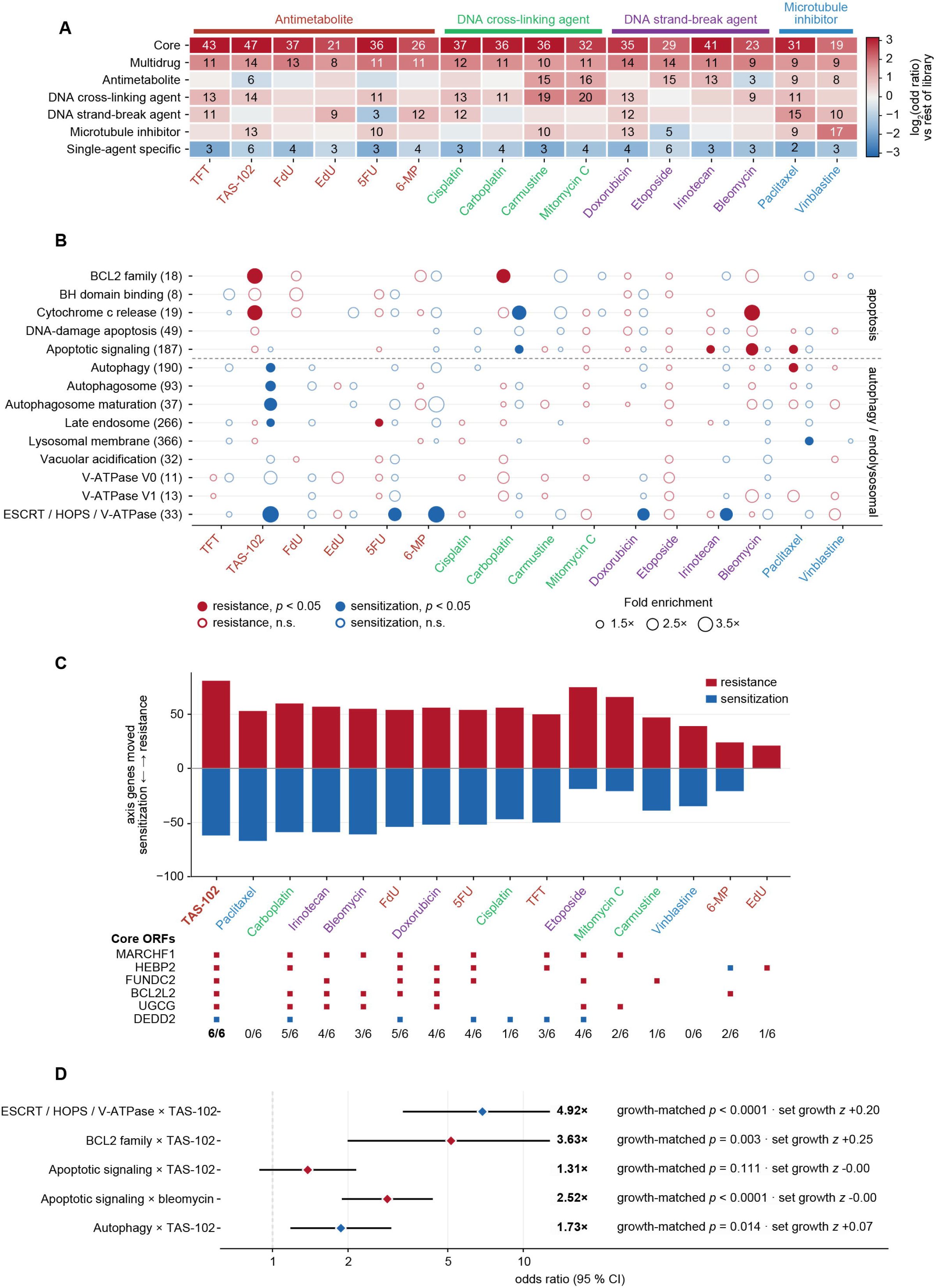

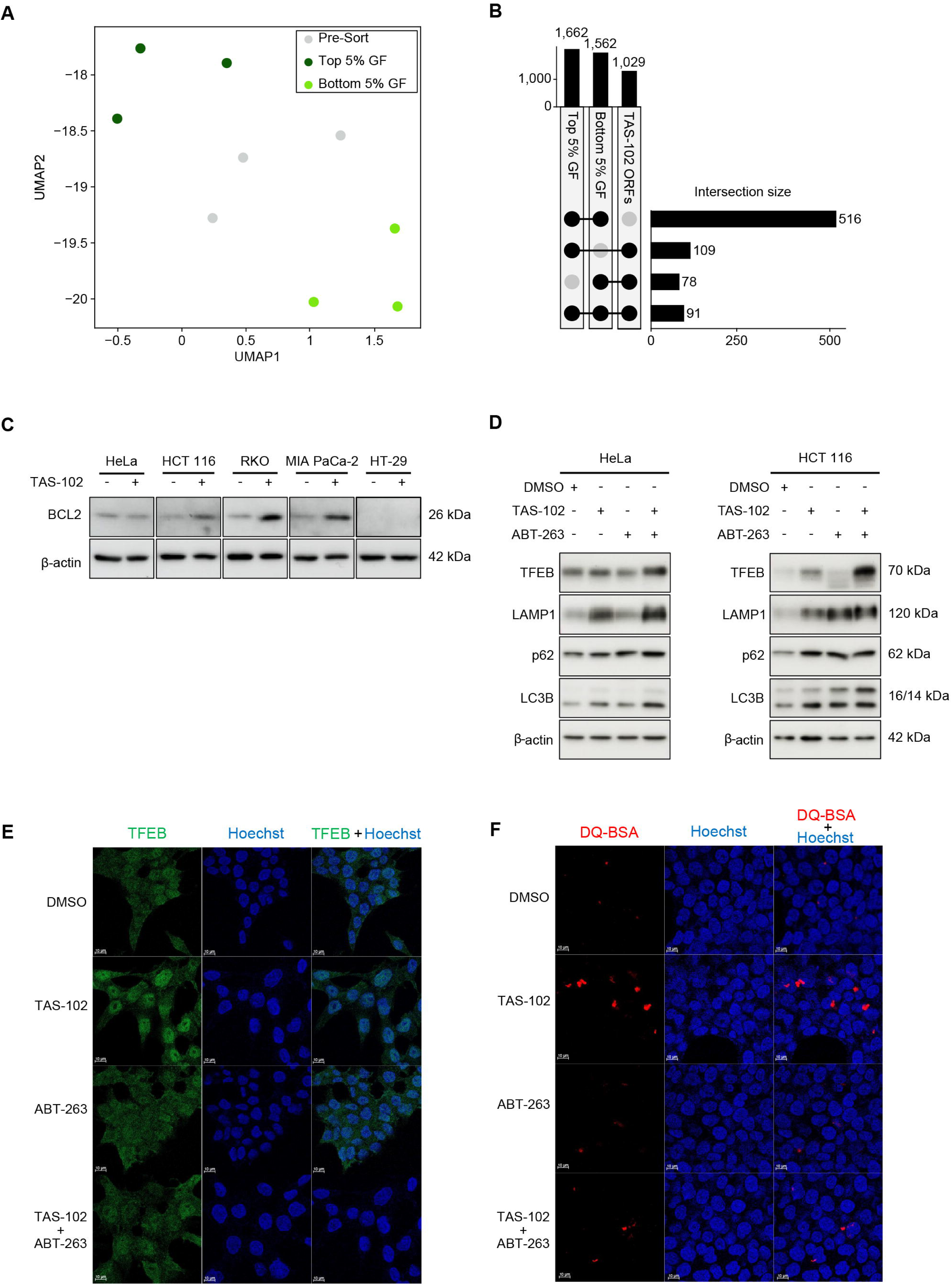

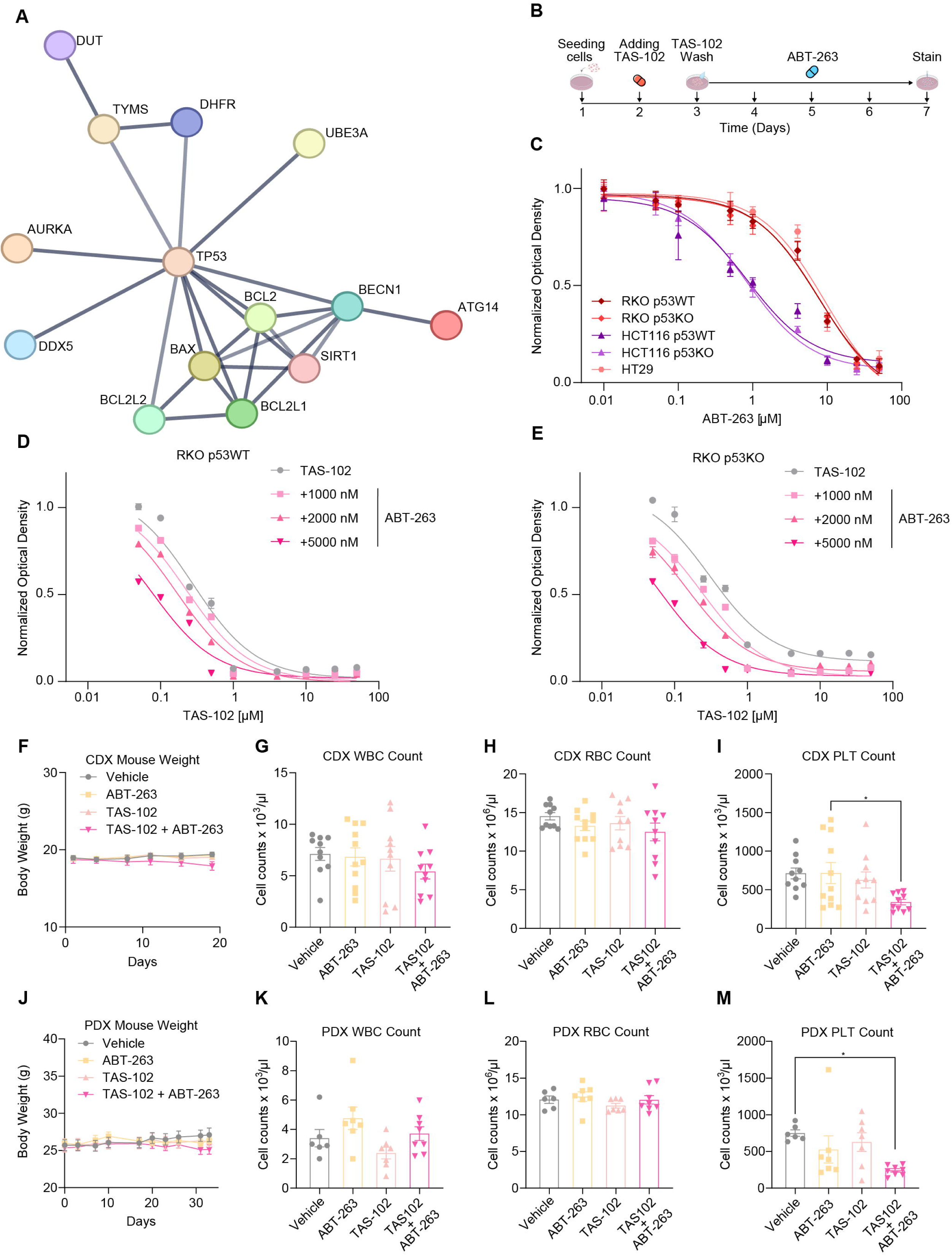

## Notes

https://dataview.ncbi.nlm.nih.gov/object/PRJNA1353565?reviewer=dc3jinggo7a0jli08lpi00boks

### Competing Interest Statement

M.M.A., A.V.B., and D.-K.K. are inventors on a patent (EP4709390A1) relating to this study. K.Y. and K.K.-H.F. are employees of Illumina Inc. The remaining authors declare that they have no competing interests.

### Summary of Updates

Manuscript PDF updated to correct character and symbol rendering issues and improve abstract formatting. Minor copyediting was applied to standardize supplementary table citations and data availability text, with no changes to scientific results or conclusions.

