## Supplementary Note and Supplementary Methods for "A Proteome-Wide Gain-of-Function Atlas Mapping the Landscape of Chemotherapy Response"

To establish the fundamental stability and kinetic properties underlying this reference atlas, we evaluated the engineered HeLa pool's insertion efficiency, longitudinal representation, and expression dynamics. Cells were sorted based on mCherry reporter signal and expanded to form the experimental pool, representing the full hORFeome landscape. Fluorescence-activated cell sorting (FACS) analysis revealed that 18.6% of cells were retrieved with mCherry-positive/mTagBFP2-negative signal (**Additional file 2: Fig. S1G, left**), achieving 150× coverage with over 3 million sorted cells, indicating robust representation of the ORFeome (**Additional file 2: Fig. S1G, right**). To verify expression quality longitudinally, stability tracking over 24 days after ORF insertion in the primary amplification series (PCR1) confirmed high recovery of integrated ORFs and stable distributions across longitudinal time points, retaining 97.5% of initial Day 3 (D3) detected ORFs through Day 24 (D24; 15,503/15,901 ORFs; **Additional file 2: Fig. S1H, left; Additional file 3: Table S2**). Lorenz curve analysis further demonstrated minimal Gini coefficient drift relative to the plasmid pool ($\Delta = 0.11$), consistent with stable representation over 24 days (**Additional file 2: Fig. S1H, right; Additional file 3: Table S2**). Co-transfection confirmed a single landing pad, verifying single-ORF integration (**Additional file 2: Fig. S1I**). Longitudinal sequencing analysis of the 14,506 quality-filtered ORFs revealed that 94.2% of ORFs (13,662/14,506) maintained stable expression relative abundance over time (**Additional file 2: Fig. S1J, left**). A minor fraction displayed longitudinal abundance shifts, segregating into two functionally distinct kinetics (**Additional file 2: Fig. S1J, middle**). A subset of 5.4% exhibited a gradual decrease marked by a continuous, linear reduction in relative abundance, driven by lineage-constrained factors such as *MYOG*, *FOXE1*, *TCF7L1*, *SOX21*, and *MSGN1* that induce partial cell cycle deceleration and mild proliferative slowing, leading to progressive population dilution rather than acute cell death [1–5] (**Additional file 2: Fig. S1J, middle**). Conversely, a distinct 0.4% fraction showed complete dropout, exhibiting a steep drop in relative abundance (**Additional file 2: Fig. S1J, middle**). Ectopic overexpression of these top complete dropout candidates **triggers** acute toxicity, lethal differentiation signaling, or severe mitochondrial complex disruption, driving swift cellular depletion from the pool. These include master developmental transcription factors (*MESP1*, *HOXB8*) [6,7], essential metabolic and signaling adapters (*ENKUR*, *DOC2B*) [8,9], and the pro-apoptotic tumor suppressor *RASSF10* [10] (**Additional file 2: Fig. S1J, middle**). Independently, sequencing of mCherry+ cells and mCherry- cells demonstrated a significant inverse correlation between inserted gene length and the reporter ratio, confirming ORF length-dependent reporter expression characteristics (**Additional file 2: Fig. S1J, right**).

### Supplementary Methods

#### Cell culturing

T-REx™-HeLa Cell Line (HeLa cells) was maintained as previously described [11], using Dulbecco's Modified Eagle's Medium (DMEM, Corning, 10-017-CV). Before the screening, they were tested for mycoplasma contamination (ATCC, 30-1012K), and cells were confirmed using short tandem repeat (STR) profiling at Genome Diagnostics Centre of The Hospital for Sick Children (SickKids) [12]. HeLa Flp-In T-REx cells engineered with the Bxb1-landing pad at the AAVS1 locus with the EF1α promoter (HeLa EF1α cells) were generously provided by the Roth Lab at the University of Toronto, currently moved to the University of Pittsburgh [13].

The human cell lines, colorectal cancer HT-29 (RRID: CVCL_0320), pancreatic cancer MIA PaCa-2 (RRID: CVCL_0428), and pancreatic ductal cell hTERT-HPNE (RRID: CVCL_C466) were cultured as recommended by ATCC. Human colorectal cancer HCT 116 and HCT 116 p53(-/-; KO) cell lines were shared by Dr. Bert Vogelstein at Johns Hopkins University [14]. Human colorectal cancer RKO and RKO p53(-/-; KO) cell lines were shared by Dr. Alessandro Carugo at MD Anderson Cancer Center [15]. Those cell lines were maintained in DMEM and authenticated again using STR profiling by ATCC or the Roswell Park Genomic Shared Resource within the last three years. The cells were routinely screened for mycoplasma; all studies used mycoplasma-free cells. Human cell cultures were maintained in media supplemented with 10% heat-inactivated fetal bovine serum (FBS; Corning, 35-015-CV) and 1% penicillin/streptomycin (P/S, Corning, 30-002-CI) at 37°C with 5-10% $CO_2$ in a humidified incubator.

#### Co-transfection assay for single-integration site validation

To confirm that HeLa EF1α landing-pad cells harbor a single active recombination site and to exclude multi-copy integration, we performed a dual-reporter co-transfection assay adapted from established Bxb1 landing-pad validation protocols [16,17]. After two passages post-thaw, HeLa EF1α cells were seeded in 6-well plates in antibiotic-free DMEM supplemented with 10% FBS. On Day 1, cells in the +Bxb1 integrase group were transfected with the Bxb1 integrase expression vector (pCAG-NLS-HA-coBxb) using Lipofectamine 3000 and P3000 reagents (Invitrogen, L3000015) in Opti-MEM according to the manufacturer's instructions, whereas the negative control group omitted the Bxb1 vector. Culture media were refreshed on Day 2. On Day 3, cells were transfected with an equimolar (1:1) mixture of promoterless attB-EGFP (attB_EGFP_Bgl2) and attB-mCherry (attB_mCherry_Bgl) donor plasmids, alongside individual single-color donor plasmids for spectral compensation, using Lipofectamine 3000, followed by replacement with complete medium on Day 4. Cells were sorted for fluorescent populations one week post-transfection and maintained until two weeks post-transfection to allow full dilution of unintegrated episomal plasmids. Mutually exclusive green (EGFP+) and red (mCherry+) fluorescence populations versus double-positive (EGFP+/mCherry+) events were then quantified by flow cytometry.

#### *En masse* LR cloning of the Bxb1-landing pad ORFeome library

Entry clones in Human Open Reading Frame Collection (ORFeome) v9.1 [18,19] with ~1000 additional clones donated by the Center for Cancer Systems Biology (CCSB) at Dana-Farber Cancer Institute (Delta space) were verified by next-generation sequencing to confirm the authenticity of their coding DNA sequences ("Verified" in column in **Additional file 3: Table S1**). Entry clones were stamped from 384-well plates onto corresponding LB agar plates containing either kanamycin (25 μg/mL; Thermo Fisher, AC611290050) or spectinomycin (50 μg/mL; Thermo Fisher, S05845G). After incubation at 37°C for 48 hours (h), the LB agar plates were scraped and collected for pooled plasmid preparation using the Qiagen Plasmid Mini Prep kit (Qiagen, 27104) - one miniprep per one 384-well plate. The amount of DNA proportional to the number of clones was pooled and mixed into a single tube to create a complete set of entry clones, where each ORF was represented in equivalent ratios. Based on these pools of Entry clones, we performed *En masse* LR cloning as previously described [18,20,21]. 150 ng of the entry pool was incubated with 5× LR Clonase (Invitrogen, 11791020) and 150 ng of destination vector (pDEST-HC-REC, Roth Lab) for 5 days, starting with a 5 μL reaction and supplementing 1 μL of 5× LR Clonase and 150 ng of destination vector each day. We used 1 μL of the LR reaction mix to perform electroporations 8 times (total of 8 μL) with 10-beta electrocompetent *E. coli* (New England Biolabs, C3020K) to generate 20 million colony-forming units, which achieves approximately 1000× library representation. The electroporated bacteria were plated onto LB agar plates containing carbenicillin (Fisher, BP26485, 100 μg/ml) and incubated at 37°C for 18 h. Colonies were collected by gentle scraping, and the final destination library was isolated using a PureLink™ Fast Low-Endotoxin Maxi Plasmid Purification Kit (Invitrogen, A35892).

#### Large-scale overexpression perturbation by constructing a HeLa cell pool with the ORFeome library

HeLa EF1α cells were passaged 3 times after thawing before transfection, and 1.3 million cells were seeded in each of twelve 10 cm dishes. We transfected 10 μg of Bxb1-recombinase expression vector (pCAG-NLS-HA-coBxb, Roth Lab) per dish using Lipofectamine 3000 (Invitrogen, L3000015) according to the manufacturer's instructions. The media was refreshed after 12 h to remove the transfection mix. After 36 h, we transfected 10 µg of the ORFeome destination library prepared in the previous section. The media was refreshed 12 h later and incubated for an additional 24 h to allow mCherry fluorescence expression to appear. On the second day, the transfected cells were pre-sorted for mCherry using fluorescence-activated cell sorting (FACS) by Sony MA900 at Flow & Immune Analysis Shared Resource (FIASR) at Roswell Park Comprehensive Cancer Center (RPCCC), and grown in 15 cm dishes for 3 more days until the residual Blue Fluorescent Protein (mTagBFP2) signal diminished and then were sorted again (post-sort) on the fourth day (5 days post-transfection) by which a 150× representation was achieved (Pre-sort: ~15 million, Post-sort: ~11 million cells). To minimize batch effects from variability within cell pools, we expanded the pool of ORFeome-integrated cells to one hundred 15 cm dishes. The cell pool was then trypsinized, and 3 million ORFeome-integrated HeLa cells (HeLa ORFeome cells) were cryopreserved per vial, with 200 vials for subsequent drug selection screens.

**Drug dose-response assays**

Drug dose-response assays were carried out using the modified protocol from the previous study [22]. HeLa cells were thawed and grown in 10 cm dishes under standard culturing conditions before an assay. We seeded 3,000 cells into 96-well plates with 100 µL of media and incubated them for 24 h to allow for attachment. Each drug was diluted to a 50 or 100 µM concentration and was serially diluted at a 1:3 ratio using culture medium to cover a range of eight concentrations. 100 µL of the diluted drug was added in triplicate and incubated for 72 h. The media were then removed, and viability was measured using AlamarBlue (Invitrogen, A50101) according to the manufacturer's instructions. $OD_{595nm}$ (Absorbance at 595 nm) reading was measured after 2 h of incubation, and the absolute Inhibitory Concentration at 50% ($IC_{50}$) was calculated using GraphPad Prism 9 software (Version 9.1.0).

**Chemotherapeutic screen**

We selected 16 chemically synthesized cytotoxic chemotherapeutic agents that primarily target the cell cycle or exert non-specific effects through DNA damage. These drugs were chosen to represent a broad range of mechanisms of action commonly employed in adult and pediatric cancer treatments, as previously described [23]. To streamline the analysis, we categorized the chemotherapeutics based on their primary mechanisms of action. In some cases, drugs were grouped under broader mechanistic categories. For instance, doxorubicin, a topoisomerase II inhibitor, and bleomycin, an antitumor antibiotic, were classified under agents that induce DNA strand breaks, reflecting their shared outcome despite differing molecular mechanisms. Based on these mechanistic classifications, the 16 chemotherapeutics were grouped into the following four categories:

- *Antimetabolites*: Fluorouracil (5-FU, Sigma, F5130), Trifluridine (TFT, Med Chem Express, HY-A0061), 5-ethynyl-2′-deoxyuridine (EdU, Toronto Research Chem - Sigma, E9386), Floxuridine (FdU, Adooq Bioscience, A10392-500), 6-mercaptopurine (Sigma, 852678), TAS-102 (Trifluridine/Tipiracil, Adooq Bioscience, 733030-01-8)
- *DNA Cross-Linking Agents*: Mitomycin C (Sigma, M5353), Cisplatin (Cayman, 13119), Carboplatin (British Pharmacopeia Reference - Sigma, BP711), Carmustine (European Pharmacopoeia Reference - Sigma, Y0002128)
- *DNA Strand Break Agents*: Doxorubicin (Tocris, 2252), Bleomycin (Sigma, B1141000), Etoposide (Sigma, PHR2717), Irinotecan (Sigma, 341205)
- *Microtubule Inhibitors:* Paclitaxel (Sigma, T7402) and Vinblastine (Sigma, V0300000)

| Chemotherapeutic drug | $IC_{50}$ |
|---|---|
| Cisplatin | 0.333 μM |
| Carboplatin | 3.37 μM |
| Carmustine | 7 μM |
| Mitomycin C | 14 nM |
| TFT | 0.75 μM |
| 5-FU | 0.1925 μM |
| EdU | 0.875 μM |
| FdU | 25 nM |
| 6-mercaptopurine | 2 μM |
| TAS-102 | 0.8 μM |
| Doxorubicin | 5 nM |
| Bleomycin | 3.3 μM |
| Irinotecan | 0.8 μM |

| | |
|---|---|
| Etoposide | 0.625 μM |
| Paclitaxel | 2 nM |
| Vinblastine | 0.3 nM |

Using $IC_{50}$ values obtained through the drug dose-response assay, we determined the appropriate dose to treat the cells in a 15 cm dish format. We seeded at least 3 million cells and treated them with drugs at their $IC_{50}$ dose for 72 h. DMSO was used as the vehicle control. Screens were conducted in five independent batches, each including baseline [24] and a set of vehicle controls to account for each specific batch. To ensure full library representation, we thawed 10 cryopreserved ORFeome-overexpressing pool vials and expanded them for 3 days, recovering a minimum of 3 million viable cells in total to maintain 150x coverage. The cells were then seeded at 3 million per dish and treated with the respective drugs in triplicate, including corresponding vehicle controls in each batch. After 72 h of treatment, cells were trypsinized, and 3 million cells were passaged into fresh dishes with continued drug treatment. After 48 h, the drug-containing medium was removed, and cells were allowed to recover in regular culture medium for 3 more days or until confluency. Recovered cells were trypsinized and pelleted (~10 million cells per pellet) for subsequent genomic DNA extraction.

**Polymerase chain reaction (PCR) amplification of ORFs from genomic DNA**

According to the manufacturer's instructions, genomic DNA was extracted from the cell pellets using the Blood and Cell Culture DNA Midi Kit (Qiagen, 13343). We performed two sets of PCRs (nested PCR) using the Q5 High-Fidelity 2× Master Mix (New England Biolabs, M0544S) and primers (Integrated DNA Technologies) that bind to i) the landing pad locus and ii) the Gateway-cloning sites, to amplify the full-length Open Reading Frame (ORF) from the landing pad with improved selectivity and reduced genomic DNA contamination. A total of 18 μg of DNA was used in the first PCR in twelve 50 μL PCR reactions, which were pooled to represent the full 150x coverage. The second PCR was performed using 2.5 μL of the previous pooled products as the template. Two 50 μL PCR2 reactions were performed and pooled before preparing the sequencing library. For NGS library preparation and to secure the complexity of the first 5 bp during sequencing using the TruSeq platform, the second PCR primers included a 5′ NNNNN sequence [25].

| Primers | Type | Sequences |
|---|---|---|

| | | |
|---|---|---|
| BxbAmp-F | Landing pad locus | 5'-CACCCACACAAAGGAAAAGGG-3' |
| BxbAmp-R | Landing pad locus | 5'-CACATTGCCAAAAGACGGCA-3’ |
| Bxb-ngs-F | Gateway-cloning site | 5'-NNNNNCTCACAAGTTTGTACAAAAAAG-3' |
| Bxb-ngs-R | Gateway-cloning site | 5'-NNNNNTGACCACTTTGTACAAGAAAG-3' |

**Sequencing library preparation**

The quality of the amplicons was confirmed on Tapestation (Agilent, G2991BA) at the Genomics Shared Resource (GSR) of RPCCC. For samples that passed quality control, primer dimers were first removed using an Ampure XP bead clean-up kit (Beckman-Coulter, A63882). Then, amplicons were sheared using a Covaris E220 Focused-ultrasonicator (Covaris, COV-500239) to sizes averaging 250-300 bp. These sheared amplicons were ligated to TruSeq adaptors and were indexed using Kapa hyperprep for sample barcodes (Illumina, 20015963; Roche 0796237100). The resulting sequencing library was finally removed from primer dimers using an Ampure kit. Finally, DNA was quantified using qPCR and loaded into NovaSeq 6000 (Illumina, 20012850), and a minimum of 3 million paired-end reads were generated for each sample.

**ORFeome bioinformatic analysis**

Bioinformatic analysis was processed in a Python Jupyter Notebook script (Python 3.11.0), which is shared as supplementary material. FASTQ files were trimmed using Cutadapt [26] to remove the remaining TruSeq adapter sequence according to Illumina, and further trimmed with Trimmomatic [27] with a sequence Phred score (LEADING:20, TRAILING:20, SLIDINGWINDOW:4:15, MINLEN:25). For the ORFeome sequence, all sequences were checked and assigned appropriate gene symbols with the NCBI gene database [28] (downloaded on August 11th, 2023). The analyzed files were aligned to the reference of human ORFeome sequences v9.1 and delta space using the Bowtie2 tool [29,30] (v2.5.2). After the alignment, post-processed FASTQ files and aligned SAM files were evaluated using FASTQC (v0.12.1) and multiQC (v1.27.1) [31,32]. SAM files were converted to BAM format with a series of post-processing steps using Samtools [33], and the raw read counts were generated using Pysam (0.23.0), including concordant and multiple reads with a MAPQ threshold of 30 [33]. The parameters of the processes were chosen based on the Cut&Run flow used in the peer ORFeome screening publication and best suited for short-read alignment [34,35]. The raw readouts were normalized by Trimmed Mean of M-value (TMM) using EdgeR (v4.0.16) [36,37].

The normalized read counts were selected based on the calculation of sequences with a median $\log_2$TMM higher than -1 to remove noise in read counts. The full dataset was further normalized using the quantile-normalization method and batch-corrected to create an equal distribution of the dataset to analyze the differences between samples using EdgeR (v4.0.16) and Limma (v3.58.1) [38]. For sample-to-sample comparison, a TMM dataset was processed using dimensional reduction analysis of Uniform Manifold Approximation and Projection (UMAP) (UMAP-learn [39]; v0.5.7). Additionally, two-dimensional principal component analysis (PCA) was performed on the transposed batch-corrected normalized expression matrix (Scikit-learn [40]) across the drug-treated samples and DMSO controls, projecting samples along PC1 and PC2 shaped by batch and colored by individual agent, drug category. Pairwise Pearson correlation coefficients (r) were then computed across the 45 drug-treated samples and rendered as a correlation heatmap. Upper-triangle correlation coefficients were partitioned into intra-replicate (same drug) and inter-treatment (different drugs) pairs, compared using a two-sample Student's *t*-test, and visualized via overlaid boxplots and stripplots. The efficiency of ORFeome silencing was assessed by quantifying fluorescence levels in cells expressing mCherry and mTagBFP2. Silencing was evaluated using the silencing ratio, defined as $\log_2(E/S)$, where E represents the number of expressing cells (mCherry$^+$ mTagBFP2$^-$) and S denotes the number of silenced cells (mCherry$^-$ mTagBFP2$^-$). This ratio reflects the difference in relative fluorescence units between cells actively expressing the ORF (mCherry$^+$ mTagBFP2$^-$) and those in which the ORF is likely silenced (mCherry$^-$ mTagBFP2$^-$). ORFs with a Silencing Ratio less than -1 ($\log_2$ fold change ($\log_2$FC) < -1) were classified as successfully silenced ("Silencing" column in **Additional file 3: Table S1**).

For more accurate statistical testing, any samples with an underlying ORF population of less than half of the maximum expressed ORF population in total samples were excluded before TMM normalization [41]. After normalization, the dataset was winsorized using DescTools [42] (0.99.57) with a 0.001% quantile range to remove extreme data points. Then, upper-quantile normalization was performed to normalize the dataset further using EdgeR [37] (v4.0.16). Batch correction was performed using a design matrix, including control and experimental sample information, multiple batches, and altered next-generation sequencing runs, to accurately adjust the batch effect using Limma [38]. Further, $\log_2$FC of TMM counts between the experimental drug condition and the corresponding batch's DMSO in the collection and statistical testing was processed using permutation testing (n=10,000). Integrated *p*-values were computed using two-tailed tests in a combination of t-values, $\log_2$ mean difference, and z-transformed rank-sum statistics, scaled with Liptak-Stouffer's z method [43]. ORFs with fewer than two non-missing measurements in either the drug-treated or batch-matched DMSO group were excluded from differential testing in that condition (**Additional file 3: Table S1**). False discovery rates (FDR) were measured based on integrated *p*-values across detected ORFs using the Benjamini–Hochberg procedure [44] ($\alpha$ = 0.20) implemented in the statsmodels package (multipletests, method='fdr_bh') and FDR q-values were computed across the tested ORFs per drug. To compare multiple-testing behavior across the 16 chemotherapeutic agents, ORFs satisfying $|log_2FC| > 0.5$ and integrated *p*-value < 0.10 were ranked by ascending integrated *p*-value and plotted against their corresponding FDR *q*-values to build drug-specific calibration curves. The data were visualized using Matplotlib [45] and Seaborn [46].

**Library representation, quality filtering, and longitudinal baseline analysis**

Sequencing reads from the input plasmid library and engineered HeLa ORFeome cell pools were aligned to the 18,388 reference ORFs in the human ORFeome library. Before downstream functional filtering, library representation was evaluated at the post-cloning stage (post-LR reaction plasmids) and the post-transfection stage following Bxb1-mediated chromosomal integration into HeLa landing-pad cells (baseline controls). Among the integrated ORFs, 14,506 ORFs were defined as quality-filtered, well-represented ORFs based on a minimum median $log_2$ TMM (Trimmed mean of M-values [36]) expression > -1 [47] across baseline samples, whereas ORFs falling below this threshold were regarded as background noise and excluded from downstream analyses.

To evaluate longitudinal library representation and uniformity in the absence of drug pressure, pooled HeLa ORFeome cells were passaged under untreated baseline conditions, sampled at D3, D7, and D9 (n = 1 each) and at D12 and D24 (n = 3 each) post-transfection, and prepared via first-round PCR amplification for sequencing. Additionally, D24 post-transfection samples (n = 3) were prepared via second-round PCR amplification for sequencing. Sequencing read counts across all longitudinal time points were normalized to counts per million (CPM). Library representation uniformity and distributional drift from the input plasmid pool through day 24 were quantified using Lorenz cumulative read distribution curves and Gini coefficients.

**Global chemoresistance atlas visualization**

To construct the global chemoresistance atlas across all 65 screening samples, including untreated baseline controls (n=5), DMSO vehicle controls (n=15), and drug-treated biological replicates (n=45 across 16 agents grouped into antimetabolites, DNA cross-linking agents, DNA strand break agents, and microtubule inhibitors; except duplicates for 6-mercaptopurine, irinotecan, and vinblastine due to insufficient ORF detection in one replicate), sample-level raw read counts were converted to counts per million (CPM), transformed as $\log_2$(CPM+1), and batch-corrected by subtracting the mean $\log_2$(CPM+1) of batch-matched DMSO controls within each of the five experimental batches to generate the replicate-level matrix. To perform drug-specificity filtering and primary cluster assignment using robust treatment-level expression, the replicate-level matrix was intersected by gene symbol with the $\log_2$FC matrix (containing TMM-normalized, batch-corrected $\log_2$FC and integrated *p*-values across the 16 treatments; n=14,506 matched ORFs). Using the $\log_2$FC matrix, ORFs were evaluated for significance ($|\log_2FC| > 0.5$ and integrated $p < 0.05$), and non-significant fold-change values were masked to zero. To resolve drug-selective response modules and minimize pan-reactive background noise, ORFs significantly altered in 1 to 4 drugs (n=10,628 ORFs) were retained from the $\log_2$FC matrix. Each ORF was assigned to its primary drug cluster based on the highest row-wise *z*-score of significant $|\log_2FC|$ across the 16 treatments in the $\log_2$FC matrix (or by maximum $|\log_2FC|$, followed by descending $|\log_2FC|$ sorting when rendering the 16-drug summary heatmap from the $\log_2$FC matrix). These defined drug clusters were then mapped onto the 65-sample replicate matrix, where ORFs within each drug cluster were ordered via agglomerative hierarchical clustering using Ward's minimum variance method with Euclidean distance in SciPy (scipy.cluster.hierarchy; clusters with ≤ 2 ORFs were ordered by descending $|\log_2FC|$ from the $\log_2$FC matrix) [48]. For heatmap visualization, the clustered matrix was sequentially standardized by column-wise *z*-scores followed by row-wise *z*-scores (display range clamped to [-2.0, 2.0]). For the adjacent pathway enrichment dot plot, representative mechanism-of-action terms derived from Gene Ontology Biological Process, Reactome, and KEGG databases were aligned vertically to the exact row midpoint of each corresponding defined drug cluster in the heatmap using GSEApy (v1.1.9) via the Enrichr API [49].

**Validation of canonical drug-target pairs**

To benchmark biological fidelity across canonical drug-target relationships, literature-curated drug-gene pairs spanning the 16 chemotherapeutics were evaluated using the $\log_2$FC matrix (TYMS with FdU, TAS-102, TFT, and 5-FU; TPMT and HPRT1 with 6-MP; MGMT with BCNU; TUBA1A and TUBB with PTX and VBL; BLMH with BLM; TK1 with TAS-102 and EdU; NQO1 with MMC; HPRT1 with VP-16 and DOX; and SLC29A1 with TFT). $\log_2$FC values were computed by subtracting the mean expression of batch-matched DMSO vehicle controls from each corresponding drug-treated biological replicate. Pairs exceeding $|\log_2FC| > 0.5$ were stratified into resistance ($\log_2FC \geq 0.5$) and sensitization ($\log_2FC \leq -0.5$) groups, ranked in descending order of $log_2FC$, and plotted as mean $\pm$ standard error of the mean (SEM) as error bars.

**Essentiality dropout dynamics and ORF length bias analysis**

To evaluate baseline essentiality and toxicity-induced dropout dynamics over longitudinal culture without drug pressure, sequencing read counts across sequential time points were normalized to counts per million (CPM), with missing values imputed as zero. Group-level mean CPM values were calculated across initial reference (T0) and day 15 (T15, n=3) samples in the primary amplification series (first PCR, PCR1), as well as untreated baseline batches at post-sort day 9 (baseline, n=5; D9), vehicle control replicates at day 14 (DMSO, n=15; D14), and extended culture replicates at day 24 (T15, n=3; D24) in the nested PCR series (second PCR, PCR2). Longitudinal $log_2FC$ values between time points were computed using a pseudocount of 1.0 as $log_2((CPM_{late}+1)/(CPM_{ref}+1))$. Candidates exhibiting consistent depletion across both series ($log_2FC$ of PCR1 T15/PCR1 T0 ≤ -1.5 and $log_2FC$ of PCR2 T15/Baseline ≤ -1.0) were identified, and the 14,506 quality-filtered ORFs were classified based on their longitudinal depletion profiles from D9 to D24 into unchanged (13,662 ORFs, 94.2%), gradual decrease (791 ORFs, 5.4%), and complete dropout (53 ORFs, 0.4%; complete depletion to 0 CPM at D24) categories. To illustrate representative depletion kinetics, $log_2FC$ trajectories across D9, D14, and D24 (normalized relative to D9) were plotted for the top five representative ORFs in both the gradual decrease and complete dropout subsets. Continuous associations between coding sequence length (both raw bp and $log_{10}$-transformed bp) and the silencing ratio were quantified using Pearson's linear correlation (*r*) in SciPy (scipy.stats) [48]. Joint density distributions were visualized using a two-dimensional logarithmic hexbin plot overlaid with an ordinary least squares (OLS) linear regression trendline using Seaborn [46].

**Automated entry clone selections for High Saturation Retesting (HSR)**

Approximately 10% of the ORFeome pool underwent retesting at higher coverage (>1,000-fold) to assess differentially represented ORFs. The selection of these ORFs relied on T-score statistical analysis of $log_2FC$ for individual drugs, grouped into eight baskets. These selections were based on core resistance or sensitizing ORFs across all drugs, individual drugs, and controls. The selection criteria for the HSR ORFeome focused on genes that demonstrated high T-scores, indicative of strong or weak proliferation phenotypes. In total, 2,217 ORFs were selected and cherry-picked from the human ORFeome using a customized robotic platform (S&P Robotics Inc., BM3-BC). These selected ORFs were then pooled and prepared within the

HeLa EF1α cells following the same *en masse* LR cloning [21]. Doxorubicin, TFT, and Cisplatin screening was carried out with matching $IC_{50}$ dosages as in the original screening, employing identical PCR ORF amplification methods and the previously mentioned ORFeome bioinformatics analysis in this method. For comparative validation between the primary and HSR screens, ORFs were classified based on Chemotherapeutic Screen (main) thresholds into “Resistant” ($log_2FC$ of main ≥ 1.0, *p*-value < 0.01), “Sensitizing” ($log_2FC$ of main ≤ -1.0, *p*-value < 0.01), and “Non-hit” ( $|log_2FC|$ ≤ 1.0, p ≥ 0.01) categories. Within each drug condition, HSR $log_2FC$ values were normalized by subtracting the mean HSR $log_2FC$ of the Non-hit group. Directional validation rates were quantified as the percentage of primary Resistant or Sensitizing hits maintaining concordant phenotypic directionality ($log_2FC$ of HSR > 0 or < 0, respectively) in the HSR screen, and cross-drug hit overlaps were visualized using nested donut chart using Matplotlib [45]. To quantify phenotypic signal amplification at higher coverage for representative candidates (*MAGEB2*, *GCNT2*, *TXNDC9*, *SLC28A1*, *TBPL1*, and *GPR21*), replicate-level $log_2FC$ values were calculated relative to matched DMSO controls, and the differential effect size was computed as $\Delta log_2FC$ = mean($log_2FC$ of HSR) - mean($log_2FC$ of main) with propagated SEM as error bars. All statistical comparisons against the Non-hit control group were performed using ordinary one-way ANOVA followed by Dunnett’s multiple comparisons test in SciPy (scipy.stats) [48], and plots were generated using Matplotlib [45].

**mRNA transcriptomics analysis**

The treatment duration and the protocol are adapted from a previous study on finding drug combinations [50]. HeLa Cells were seeded at 3 million cells in 15 cm dishes and treated with an $IC_{50}$ of 5 nM Doxorubicin for 24 h and 120 h. After the treatment, cells were pelleted, and each cell pellet consisted of a minimum of 1 million cells per control and experimental set. Each cell pellet was preserved in TRIzol (Qiagen, #79306) and stored at -20℃ before being sent to Genomics Shared Resources (GSR) at RPCCC. Purified RNA was prepared using the miRNeasy micro kit (Qiagen, 217084), and quality control of the RNA integrity number (RIN) was measured using an Agilent Bioanalyzer 2100 (Agilent, G2939BA). Sequencing libraries were prepared with the KAPA mRNA HyperPrep Kit (Roche, KK8581). The final RNA-seq libraries were sequenced on Illumina NovaSeq 6000. Nextflow version 24.10.4 [51] with nf-core/rna-seq v3.18.0 [52] was utilized using Docker (v27.5.1). The sequencing files were aligned to the GRCh38.p14 human genome[53]. The workflow was configured to perform adapter trimming with Trim Galore [54], followed by alignment using STAR [55] (v2.7.10a) and quantification using RSEM [56] (v1.3.1). Ribosomal RNA reads were removed using

SORTMERNA [57] (v4.3.7) before quantification. Downstream analysis used count-based quantification, matching the configuration of the ORFeome bioinformatic pipeline. Statistical analysis was performed using edgeR's exactTest function (v4.0.16) [37]. Data comparison was visualized using Upset Plot [58].

**Preparation for single-cell RNA-seq**

Cryopreserved HeLa cell stocks after drug selection screen were thawed in a 37 °C water bath for 1 min and transferred to DMEM (Gibco, 11965092) containing 10% FBS (Gibco, 16000044), followed by centrifugation at 300 × g for 5 min at room temperature. Cells were further washed twice with cold DPBS (Gibco, 14190144) containing 0.04% BSA (Miltenyi Biotec, 130-091-376) and filtered through 40 µm cell strainers (Flowmi, BAH136800040). Cell suspensions were then loaded, targeting recovery of 10,000 cells per channel to construct single-cell RNA sequencing libraries using the Chromium Next GEM Single Cell 5' v2 kit (10x Genomics, PN-1000263) according to the manufacturer's instructions. Generated libraries were sequenced on an Illumina NovaSeq 6000 at a paired-end, 150 bp reads configuration, aiming for 400 million reads per library.

**Single-cell RNA-seq bioinformatic analysis**

Reads were aligned to the GRCh38 human genome (2020-A, 10x Genomics) using Cell Ranger (v6.0.2, 10x Genomics) [59] with an include-introns parameter. Filtered count matrices were further processed with Scanpy (v1.9.1) [60]. Low-quality cells with < 2,000 total counts, < 500 detected genes, > 10% mitochondrial gene counts, and > 0.4 Scrublet (v0.2.3) [61] doublet scores were filtered out. Raw counts were normalized per cell to 10,000, and ln(x+1) was transformed and further used as gene expression values. The top 2,000 highly variable genes were selected with the scanpy.pp.highly_variable_genes function with seurat_v3 [62] for each dataset, and principal component analysis (PCA) was performed with the selected highly variable genes. The neighborhood graph was built from the top 50 principal components and visualized using UMAP [63]. Clustering was performed with the Leiden method [64] with 0.5 resolution, and the Wilcoxon rank-sum test [65] was used to calculate marker genes of each cluster.

**Cross-modality comparison of bulk mRNA-seq and single-cell RNA-seq**

To compare endogenous transcriptional shifts with overexpression-driven survival signatures under doxorubicin treatment, differential expression profiles from wild-type HeLa mRNA transcriptomics were merged by gene symbol with single-cell RNA-seq differential expression profiles from HeLa OE cells and ORFeome screen $log_2FC$ values. Genes were stratified across a quadrant scatter plot using an endogenous expression threshold of $|log_2FC| = 0.5$, with point sizes of representative doxorubicin resistance and sensitization markers (*BCL-2*, *BCKDHA*, *DUSP4*, *HDAC4*, and *NUPR1*).

**ORF gene categorization**

Within the quality-filtered ORFs 14,506 ORFeomes, ORFs were categorized into five categorical subsets. Among the categorized ORFs, those with a $log_2FC$ threshold of ±0.5 and an Integrated *p*-value < 0.05 were identified as differentially represented ORFs. Within this set, enriched ORFs (positive $log_2FC$) are interpreted as putative resistance drivers, conferring a survival advantage under treatment, while depleted ORFs (negative $log_2FC$) suggest sensitization drivers, indicating increased drug sensitivity. First, 197 core genes were selected based on ORFs with a filter of at least 7 drugs (≥ 7) (at least half of the maximum observed breadth of 13 agents with more than 1 drug in at least 3 categories). Second, 1,072 categorical genes were assigned based on the recalculation of the median($log_2${drug categories}) - median($log_2${rest of categories}) from the previously mentioned statistical testing and filtered to differentially represented ORFs. Third, 3,832 peripheral genes were identified as ORFs significantly altered ($|log_2FC| \geq 0.5$ and *p*-value < 0.05) in response to specific chemotherapeutic drugs, but not in others. For each drug, genes meeting these criteria were selected and then filtered to exclude any genes that also met the same significance criteria in other drugs, thereby ensuring drug-specificity. Fourth, 6,556 multidrug genes were defined as remaining ORFs significantly altered across 2 to 6 chemotherapeutics ($2 \leq x < 7$), spanning multiple drug classes. Lastly, 2,849 non-respondent genes were picked with ORFs, with a filter for no enriched or depleted ORFs. To avoid any overlapping of ORFs between categories, the selection was prioritized sequentially from "Core" to "Non-respondent" genes.

**Gene properties analysis**

Length of each ORF (ORF Length) and its GC percentage (GC Content) were both derived from human ORFeome v9.1 sequences and delta space references after alignment as described above. We adopted some of the gene properties from our previous study [18,20]. Genotype-Tissue Expression project (GTEx) [66] v6 transcriptome data (downloaded on March 15th, 2016) was processed with R v3.5.1 to extract normalized log-read counts. In this dataset, 16 brain subregions were collapsed into three brain tissues: basal ganglia, cerebellum, and others. The median expression of each gene across all tissue samples was calculated, and a cutoff of >5 was applied. Cell line samples were excluded, and the analysis was restricted to the set of protein-coding genes. We considered these values as a reference for gene expression level (Abundance), which were calculated as mentioned in our previous study [18]. This dataset was also used to calculate the number of tissues each gene was expressed in (Broadness). Genes' evolutionary age information (Age) was adopted from the Protein Historian database [67], with PPODv4_Jaccard as the family database and Dollo parsimony model as the reconstruction algorithm. The database of disordered protein predictions ($D^2P^2$) [68] was used to calculate the fraction of disordered regions relative to full protein length (Disorder). First, all human $D^2P^2$ IDs mapped to Ensembl IDs from "genomes.protein.gz" were extracted, and then IDs were used along with the "consensus_ranges.tsv.gz" file, containing the start and end positions of each predicted disordered region, to compute the total disordered length for each protein. The length of each protein was calculated using the "genomes.fa.bz2" file. The DAVID functional annotation tool [69] was used to retrieve information for other properties. The list of Entrez IDs was submitted, and the Functional Annotation Table for the following databases was downloaded on January 16th, 2024: OMIM_DISEASE, Gene Ontology (GO) - GOTERM_BP_DIRECT (Biological Process), GOTERM_CC_DIRECT (Cellular Components), GOTERM_MF_DIRECT (Molecular Function), PUBMED_ID, and UP_TISSUE. The number of publications (PubCount) was determined by the number of unique PubMed IDs per gene. Similarly, GO term datasets were merged and used to calculate the number of associated terms for each gene (GO Terms), UP_TISSUE [70] to assess the number of tissues in which each gene's expression level was upregulated (Tissue Count), and OMIM [71] to find the number of diseases per gene (OMIM Terms). To assess the extent of each gene's involvement in protein complexes, using the CORUM database version 4.1 [72] (downloaded on January 23rd, 2024), the number of protein complexes associated with each gene (CORUM Complexes) was compared.

**Biomarker analysis**

To assess the clinical relevance of gene subsets identified in our screening, we performed a multi-tiered biomarker overlap analysis using TheMarker database [73], a curated resource of disease, biomarker, and drug associations. Gene matching was based on NCBI Gene identifiers, and only "verified = 1" and "Silencing = No silence" ORFs from the screening dataset of quality-filtered ORFs were included in the analysis. Those ORFs were intersected with cancer-specific biomarker entries in the database, and for each gene subset, the proportion of genes overlapping with cancer-associated biomarkers, classed as "ICD-02: Benign/in-situ/malignant neoplasm," relative to the total ORFs in that subset was calculated to evaluate enrichment for clinically validated cancer-related genes (cancer-associated ORFs). We then examined the association between biomarker-annotated ORFs and chemotherapeutic drugs; 7 out of the 16 drugs profiled in the screening were represented in the database, and we quantified the proportion of ORFs within each subset linked to one or more of these drugs (drug-linked ORFs), connecting screening hits to existing pharmacogenomic knowledge. Finally, we assessed the representation of each gene subset within the total biomarker pool by calculating the proportion of verified, non-silenced cancer biomarker annotations relative to the total biomarkers mapped to each respective subset in the database. All overlap proportions were normalized using *z*-scores to enable comparisons across subsets and highlight differential enrichment patterns. The *z*-scores were visualized as heatmaps, and the numbers of four biomarker types within each ORF category were calculated as relative percentages (%) within each ORF category and displayed in stacked bar graphs. The stacked bars represent percentages for each category separately and do not correspond to the sum of total biomarker numbers.

**Proliferation-associated ORFs using *z*-score ranking**

To assess the clinical relevance of cancer proliferation-related ORF populations, we employed a combined growth *z*-score ranking approach. For each experimental batch, changes in ORF abundance were calculated as the $\log_2$FC between mean expression under DMSO treatment and baseline expression, following a strategy adapted from CRISPR screening methodologies [74]. These differences, representing perturbation responses, were computed for each batch independently. The resulting difference matrix was then growth *z*-score normalized batch-wise, transforming each batch's distribution. To aggregate signals across multiple experimental batches, an inverse *z*-score transformation method from a previous study [75] was applied per ORF. Specifically, each *z*-score was converted back to an approximated original scale using the Inverse *z*-score Transformation. The combined growth *z*-score for each ORF was then calculated by averaging these inverse-transformed values across all batches. ORFs were ranked in descending order based on their combined growth *z*-scores.

**Comparative benchmarking analysis**

To benchmark baseline proliferation phenotypes, combined HeLa OE growth *z*-scores (DMSO versus baseline) were compared against lentiviral ORF proliferation *z*-scores in MCF-7 cells across 364 overlapping ORFs [76]. Length-dependent phenotypic dynamic range was evaluated by comparing absolute $\log_2$FC distributions across 1kb ORF length intervals (up to 9 kb) between our BOGO platform and published lentiviral ORF screens [76,77]. For cross-modality drug response comparisons, HeLa OE and NALM-6 CRISPR-Cas9 KO profiles [78] were independently $z$-score standardized within each of the eight shared chemotherapeutics, and top quadrant candidates were prioritized by Euclidean distance from the origin ($\sqrt{Z_{OE}^2 + Z_{KO}^2}$). All datasets were integrated via NCBI gene symbols and visualized using Matplotlib [45] and Seaborn [46].

**Prognostic evaluation using precision–recall metrics and survival analysis**

The ORF list was integrated with prognostic annotations from the Human Protein Atlas (HPA) [79] (v24.0) through a two-step merge using Entrez gene IDs and gene symbols. Genes annotated with a favorable and unfavorable prognosis (defined as being linked to poor outcome in at least one cancer type) were designated as "Hits". Based on the first hit, the first rank was computed. To assess performance, precision was calculated as the proportion of Hits among the top-ranked genes, and recall as the proportion of total Hits identified up to a given rank. In addition, the maximum precision at or beyond each rank was computed using a cumulative maximum function applied in reverse order. Both the maximum and recall curves were visualized to evaluate the model's ability to prioritize the proliferation-related ORF associated with poor prognosis. A parallel analytical pipeline was applied to assess the association between ranked ORFs and drug resistance, using manually curated resistance data from The Cancer Genome Atlas (TCGA), following criteria from prior work [80]. For survival analysis, we used RNA-seq data from HPA. Patients were stratified into high and low expression groups for each gene using the best expression cut-off, provided by HPA, defined as the Fragments Per Kilobase of transcript per Million mapped reads (FPKM) threshold that yields the lowest log-rank $p$-value between the two groups.

**Drug-response similarity network and functional neighborhood enrichment analysis**

For mapping global functional relationships across the 16 chemotherapeutic treatments, we constructed an ORF-ORF drug-response similarity network encompassing all 11,595 drug-responsive ORFs ($|\log_2 FC| \geq 0.5$, integrated $p$-value < 0.05 in at least one individual drug; 62 Categorical ORFs that were significant only at the pooled drug-class level had all-zero single-agent response vectors and were excluded). For each ORF pair sharing at least 8 measured drug conditions, we computed the pairwise-complete Pearson correlation coefficient ($r$) across their signed categorical response profiles (sign($\log_2$FC) for significant hits; 0 otherwise). Each ORF was connected to its $k$ = 10 most correlated neighbors (retaining the maximum positive weight w = max($r$, 0.01) for mutual edges), yielding an undirected graph of

11,595 nodes and 100,347 edges. Network coordinates were generated using a deterministic Fruchterman-Reingold force-directed layout (60 iterations; seeded with PCG64 random generator seed = 0), aligned along principal axes via singular value decomposition (SVD), and radially equalized onto a circular disc.

To identify spatially concentrated biological pathways on the similarity network without layout-coordinate artifacts, we performed a correlation-rank neighborhood enrichment analysis modeled from Spatial Analysis of Functional Enrichment (SAFE) [81]. For each node, its local neighborhood was defined as the 100 most correlated ORFs by pairwise-complete Pearson (*r*). Gene Ontology (GO) annotations were compiled from Gene Ontology resources using a script from SAFE, restricted to terms containing 10 to 600 library ORFs. Local neighborhood enrichment per node was evaluated by a one-sided hypergeometric test ($p$-value < 0.001). Domain spatial concentration was quantified by the number of enriched nodes and benchmarked against an empirical null distribution of 3,000 size-matched random gene sets to compute empirical $p$-values (with within-domain Šidák correction and across-domain Benjamini-Hochberg FDR adjustment).

**Pathway enrichment grid, bipartite network, and endolysosomal cascade analysis**

To evaluate pathway convergence across all 16 chemotherapy agents without network clique amplification, 14 pre-specified apoptotic, autophagic, and endolysosomal gene sets were tested across all 16 chemotherapeutics in both resistance $\log_2$FC ≥ 0.5, $p$-value < 0.05, and sensitization $\log_2$FC ≤ -0.5, $p$-value < 0.05 directions using one-sided hypergeometric tests relative to the measured library per drug (444 tests total). To control for baseline proliferation bias, headline enrichments were benchmarked against 20,000 random gene sets matched to the combined growth $z$-score decile distribution of the query set.

For the gene-drug bipartite network, we extracted all 15 Core ORFs (≥ 7 drugs) comprising nine ORFs annotated to apoptotic signaling GO terms or the canonical *BCL-2* family together with the six Core ORFs of the *BCL-2* family-mediated autophagy-endolysosomal axis (*MARCHF1, HEBP2, FUNDC2, BCL2L2, UGCG*, and *DEDD2*; 113 significant gene-drug interactions). Each Core ORF node was positioned at the |$\log_2$FC|-weighted centroid of its interacting drugs and colored by its net response direction across treatments. For the endolysosomal cascade analysis, $\log_2$FC profiles were mapped across the 33 responsive subunits of the obligate ESCRT-0, ESCRT-I, ESCRT-II, ESCRT-III, HOPS, and V-ATPase complexes and contrasted with the nine TAS-102-responsive Core vacuolar and autophagic ORFs; directional concordance under TAS-102 across the cascade was evaluated by a one-sided exact binomial sign test.

**Lysosomal functional screen using Lysotracker**

The FACS method followed the protocol from the previous study [82]. HeLa ORFeome cells were thawed and pooled to maximize ORF library coverage, then allowed to recover for three days before the experiment. All sorting experiments were performed with three independent biological replicates (n =3), where each replicate was processed independently. For the screening, 3 million cells were seeded per 15 cm dish for the DMSO control while cells were still in suspension, before full adherence. Following 72 h of DMSO treatment, lysosomal activity was assessed by staining cells with 50 nM LysoTracker Green for 20 minutes at 37°C. After two DPBS washes, cells were sorted on a Sony MA900 flow cytometry cell sorter, gating for mCherry positivity and then for enhanced lysosomal activity (High green fluorescence, upper 5%) and diminished lysosomal activity (Low green fluorescence, lower 5%) populations alongside the pre-sorted input control. Sorted cells were collected, seeded in 6-well plates, and expanded for one week before being cultured to full confluence in 15 cm dishes. Genomic DNA extraction, PCR amplification, and sequencing were performed as previously described for chemotherapeutic screening. Sequencing data from the sorted populations were analyzed using the ORFeome bioinformatics pipeline to calculate $\log_2$FC and *p*-values by comparing the pre-sort control or lysosomal activity. Data visualization was done using UMAP [63] and Upset Plot [58]. Functional and pathway enrichment analyses focusing on the TGF-beta signaling pathway were conducted using DAVID [69]. Survival analysis and patient stratification based on the selected genes were performed using data from the TCGA PanCancer Atlas [83] via cBioPortal [84] (accessed on July 7th, 2025), and visualized using Matplotlib [45]. Because GHSR transcript was undetectable in most TCGA tumors (66.1% of samples; 6,540 of 9,892), precluding a conventional median split, tumors were stratified into detectable (High, n = 3,352) versus undetectable (Low, n = 6,540) GHSR groups, whereas CCDC86 was quantifiable across all samples and stratified by the pan-cancer median into High (n = 4,946) and Low (n = 4,946) expression groups. Overall survival associations were evaluated using Kaplan–Meier curves alongside cancer-type-stratified log-rank tests and stratified Cox proportional hazards regression models (across 28 cancer types with complete survival follow-up [n = 9,610] for GHSR and 26 cancer types [n = 9,065] for CCDC86).

**Protein-Protein interaction network analysis**

To investigate the overlap between lysosomal modulators identified in the LysoTracker screen and regulators of TAS-102 response, significant genes ($|\log_2FC| \geq 0.5$, *p*-values < 0.05) from the upper 5% LysoTracker population, lower 5% LysoTracker population, and TAS-102 drug screen were combined into a single dataset for downstream network analysis. Genes that did not meet the significance criteria, as well as genes significant in only one dataset, were excluded, retaining only those with significant values present in at least two of the three datasets. The list of gene symbols was imported into Cytoscape (v3.10.4) [85] to generate a protein-protein interaction network using the STRING database [86] (accessed September 29th, 2026), as described above, using a confidence score threshold of 0.7. The largest connected subnetwork was isolated and visualized using a grid layout. GO enrichment analysis was performed using DAVID [69], and the network was reorganized using the attribute grid layout by incorporating an additional column that positioned genes associated with the most relevant/selected enriched GO terms in the first row. The Therapeutic Target Database (TTD) [87] was used to identify genes associated with approved or investigational drugs and chemical compounds, which were visualized using dashed borders mapped to a corresponding binary attribute. The Omics Visualizer [88] in Cytoscape was used to generate pie chart visualizations displaying the $log_2FC$ from each dataset. To illustrate the canonical literature-curated bridge connecting the TAS-102 target TYMS to the *BCL-2* family and autophagy regulators, a protein-protein interaction network was constructed in STRING [86] (confidence score > 0.7; <5 interactors in the 1st and 2nd shells).

**Western blot analysis for BCL-2 and lysosomal pathway protein detection**

For BCL-2 protein detection across cell lines, HeLa (0.8 μM), RKO (0.6 μM), and HCT 116, MIA PaCa-2, and HT-29 (2 μM) cells were treated with DMSO or TAS-102 for 24 h. For lysosomal pathway analysis, HeLa cells were treated with DMSO, 1 μM TAS-102, 1 μM ABT-263 (MedChemExpress, HY-10087), or 1 μM TAS-102 + 1 μM ABT-263 in combination, and HCT 116 cells were treated with DMSO, 1 μM TAS-102, 0.5 μM ABT-263, or 1 μM TAS-102 + 0.5 μM ABT-263 in combination for 48 h. After 24 h or 48 h of incubation, cells were lysed in sample buffer (60 mM Tris-HCl (pH 6.8), 2% SDS, 10% glycerol, 5% β-mercaptoethanol, and 0.01% bromophenol blue) or EBC lysis buffer (50 mM Tris-HCl (pH 8.0), 200 mM NaCl and 0.5% NP-40), supplemented with a protease inhibitor cocktail (Sigma-Aldrich, 11836153001) for 30 min on ice. Total cell lysates were sonicated three times for 10 sec each and centrifuged at 15,000 × g at 4˚C for 15 min. The supernatants were collected, and protein concentration was measured using the BCA protein assay (Thermo Fisher Scientific, Inc., 23227). Equal amounts of protein were separated by 8- 15% SDS–PAGE and transferred to a PVDF membrane (Bio-Rad Laboratories, Inc., 1620177). The membranes were blocked in 1X Tris-buffered saline with 0.1% Tween 20 and 5% skim milk (CellNest, CNS109) at room temperature for 30 min and incubated with primary antibodies at 4˚C overnight or at room temperature for 1 h and with secondary antibodies at room temperature for 1 h. The signal was visualized with ImageQuant™ 800 (Cytiva™, 29399481) using an enhanced chemiluminescence detection system (GenDEPOT, LLC, W3652-020). The following antibodies were used:

| Antibodies (Ab) | Ab type | Vendor | Catalog | 1st Ab Dilution | 2nd Ab Dilution |
|---|---|---|---|---|---|
| BCL-2 (D17C4) | 1st | Cell Signaling | 3498; RRID: AB_1903907; Lot 6 | 1:500 | 1:2500 |
| TFEB | 1st | Cell Signaling | 4240; RRID: AB_11220225; Lot 2 | 1:1000 | 1:2500 |
| SQSTM1/p62 | 1st | Cell Signaling | 5114; RRID: AB_10624872; Lot 6 | 1:1000 | 1:2500 |
| LAMP-1 (H4A3) | 1st | Santa Cruz | Sc-20011; RRID: AB_626853; Lot C0620 | 1:500 | 1:3000 |
| LC3B | 1st | Novus Biologicals | NB100-2220; RRID: AB_10003146 | 1:2000 | 1:5000 |
| β-Actin | 1st | Sigma-Aldrich | A1978; RRID: AB_476692; Lot 0000124305 | 1:10000 | 1:10000 |
| Peroxidase AffiniPure Goat Anti-Rabbit IgG (H+L) | 2nd | Jackson ImmunoResearch | 111-035-003; RRID: AB_2313567 | | |
| Peroxidase AffiniPure Goat Anti-Mouse IgG, Fcγ fragment specific | 2nd | Jackson ImmunoResearch | 111-035-071; RRID: AB_2338506 | | |

#### Immunofluorescence assay

HCT 116 cells were seeded at a density of 1 × $10^5$ cells per well in 12-well plates. On the following day, cells were treated with DMSO, 2 μM TAS-102, 0.5 μM ABT-263, or 2 μM TAS-102 + 0.5 μM ABT-263 in combination. After 24 h of incubation, the cells were fixed with 2% paraformaldehyde at room temperature for 10 min and permeabilized with 0.5% Triton X-100 for 5 min. Next, the cells were treated with 2% bovine serum albumin (A0100-010; GenDEPOT) for 1 h and then incubated with primary antibodies in 2% BSA at 4°C overnight. The next day, the cells were treated with Alexa Fluor 488 goat IgG antibody (A-11034; Thermo Fisher Scientific Inc.) in 2% BSA at room temperature for 1 h. Nuclei were stained with 1 μg/mL Hoechst 33342 (B2261; Sigma) for 5 min at room temperature. Finally, the coverglass slips were then mounted onto glass slides using an anti-fade fluorescence mounting medium (ab104135; Abcam). Confocal imaging (LSM800; CLSM) was subsequently performed. The primary antibodies used were against TFEB (1:500; 4240; Cell Signaling). To quantify nuclear TFEB fluorescence using ImageJ software (v1.53K; National Institutes of Health), nuclear masks were delineated based on the Hoechst 33342 channel to measure the average nuclear fluorescence intensity across five independent fields of view per condition. To correct for differences in cell number across microscopic fields, the average nuclear TFEB fluorescence intensity of each image was normalized by dividing by the total number of nuclei counted within that field. Statistical analysis was performed using two-way ANOVA (with TAS-102 and ABT-263 as factors) with Tukey's multiple comparisons test.

**DQ-BSA assay**

To assess lysosomal proteolytic activity, the DQ-BSA assay was performed as previously described [89]. HCT 116 cells were seeded at 1 × $10^5$ cells per well on coverglass slips in a 12-well plate. The following day, cells were treated with DMSO, 2 μM TAS-102 or 0.5 μM ABT-263 or a combination of both. Following 24 h of drug treatment, cells were incubated with 10 μg/mL DQ™ Red-BSA (D12051; Thermo Fisher Scientific Inc.) at 37°C for 24 h in the dark. After incubation, cells were washed with DPBS and fixed in 2% formaldehyde for 10 min at room temperature. Nuclei were stained with 1 μg/mL Hoechst 33342 for 5 min at room temperature. The coverglass slips were then mounted onto glass slides using an anti-fade fluorescence mounting medium. Confocal imaging (LSM800; Carl Zeiss) was subsequently performed, and fluorescence intensity was quantified using ImageJ (v1.53K). Statistical analysis was performed using two-way ANOVA (with TAS-102 and ABT-263 as factors) with Tukey's multiple comparisons test.

**Measurement of lysosomal pH**

To assess lysosomal acidification, the lysosomal pH was measured as previously described [90]. HCT 116 cells were seeded at $1 \times 10^4$ cells per well into a 96-well plate. The following day, cells were incubated in complete media containing DMSO, 2 μM TAS-102, 0.5 μM ABT-263, or 2 μM TAS-102 + 0.5 μM ABT-263 in combination. Following 24 h of drug treatment, cells were incubated with 1 μM LysoSensor™ Yellow/Blue DND-160 (L7545; Thermo Fisher Scientific Inc.) for 5 min in the dark. After incubation, cells were washed with DPBS, and then the media was replaced with Hanks' Balanced Salt Solution (HBSS) (Gibco, 14025092) supplemented with 10% FBS for 10 min for test wells. To generate a standard curve, cells were incubated with a buffer (25 mM HEPES, 115 mM KCl, 1.2 mM $MgCl_2$, 10 mM glucose, 10 μM nigericin) calibrated to pH 4.5, 5.0, 5.5, 6.0, 6.5, or 7.0 as described previously. Fluorescence was measured using a PerkinElmer EnSpire multimode plate reader with excitation/emission filters set to 329/440 nm and 380/540 nm. Lysosomal pH values (n = 5 per group) were determined by linear interpolation against the standard curve generated from the pH 4.5–7.0 calibration buffers. Statistical analysis was performed using two-way ANOVA (with TAS-102 and ABT-263 as factors) with Tukey's multiple comparisons test.

**Cell proliferation assay**

To assess the cell proliferation difference after drug treatment, a method was adopted from the previous protocol [91]. HCT 116 cells were seeded in a 12-well plate at a density of $5 \times 10^4$ cells per well. The following day, cells were treated with DMSO, 2 μM TAS-102, 100 nM Bafilomycin A1, or 2 μM TAS-102 + 100 nM Bafilomycin A1 in combination. Cells were harvested and counted at 0, 24, and 48 h after drug treatment. Each count was performed in six independent replicates at each time point, and the mean values were used to generate the proliferation graphs. Statistical analysis was performed using two-way ANOVA (with treatment and time as factors) with Tukey's multiple comparisons test.

**Cytotoxicity assay**

To examine drug cytotoxicity in human cells, we followed the protocol from the previous publication [92]. Cells were plated at a density of 5,000 cells/well in a 96-well plate and treated with TAS102 at 0, 0.01, 0.05, 0.1, 0.5, 1, 4, 10, 25, and 50 μM concentrations for 24 h. The media was replenished with media containing either 0, 25, 50, 100, 200, 500, 1000, 2000, or 5000 nM concentration of ABT-263, followed by a 96-hour incubation. Cells were stained with 0.5% Methylene Blue for 30 minutes, rinsed with water, dried, solubilized in 5% SDS in DPBS, and read at 650 nm. The average of the control was used to normalize the raw absorbance. Combinatorial synergy scores were computed across the dose–response matrices using the

Bliss independence, Highest Single Agent (HSA), and Zero Interaction Potency (ZIP) reference models. The graph was generated using GraphPad Prism 9, and statistical analysis was performed using two-way ANOVA followed by Tukey's multiple comparisons test.

**Cell Line-Derived Xenografts (CDX) and Patient-Derived Xenograft (PDX) Models**

Female SCID/CB17 mice, aged 6 to 8 weeks, were sourced from a colony bred and maintained at RPCCC. Mice were inoculated subcutaneously into the left flank with exponentially growing HCT 116 tumor cells ($1×10^6$ per mouse) suspended in a 1:1 mixture of DPBS and Matrigel. Mice were monitored daily, while body weight and tumor size were measured twice a week. Tumor dimensions were measured with digital calipers and calculated using the formula: (length × width$^2$)/2. When the tumor volume reached 50 to 100mm$^3$, mice were randomly divided into four treatment groups (n = 10-11 in each group in CDX and n = 6-8 in each group in PDX): (1) Vehicle Control, (2) 50 mg/kg ABT-263, (3) 50 mg/kg TAS-102 (MedChemExpress, Cat. No. HY-16478), and (4) Combination (50 mg/kg TAS-102 + 50 mg/kg ABT-263). TAS-102 was dissolved in 10% 2-hydroxypropyl-β-cyclodextrin (HPCD) in DPBS, while ABT-263 was prepared in 10% DMSO, 40% PEG300, 5% Tween-80, and 45% saline. For the Combination group, TAS-102 was given 4 h before ABT-263. All drugs were freshly prepared and given via oral gavage on a 5-day-on, 2-day-off schedule.

Donor and experimental colorectal patient-derived xenografts (CRC PDX-03-26-4p) were grown in male SCID/CB17 mice from RPCCC. Tumors were grown in donor mice to a volume of around 1500 mm$^3$, harvested rapidly after euthanasia, cut into 2×2×2 mm$^3$ pieces under sterile conditions, and surgically implanted into the left flank. Surgical staples were removed about ten days post-implantation. Tumor growth and mouse weights were measured twice a week. Once the average tumor volume reached 50-100 mm$^3$, mice were randomized into four groups: PDX model (1) Vehicle Control, (2) 50 mg/kg ABT-263, (3) 50 mg/kg TAS-102, and (4) Combination (50 mg/kg TAS-102 + 50 mg/kg ABT-263). The drugs were prepared as in the CDX experiment and administered via the same route and dosing schedule.

Investigators were not blinded to treatment group allocation due to the staggered sequential dosing schedule; all tumor dimensions, endpoint tumor weights, and hematologic parameters were measured using objective quantitative instruments (digital calipers, analytical balance, and automated Heska HemaTrue Analyzer).

**Mouse Model Monitoring**

Mice were housed in micro-isolator units and were provided with food and water ad libitum.

Daily monitoring of animal health, body weight, and tumor growth, as well as endpoint procedures, was performed following the protocol and guidelines approved by the Institutional Animal Care and Use Committee (IACUC). The facility is certified by the Association for Assessment and Accreditation of Laboratory Animal Care (AAALAC) International and complies with regulations and standards of the US Department of Agriculture and the US Department of Health and Human Services.

At the endpoint, when the tumor diameter reached 20 mm, mice were euthanized and subjected to necropsy and organ collection. Tumor growth assessment was stopped once a mouse reached the endpoint (typically in a vehicle-treated cohort), while the survival data were still collected in the study. Intracardiac blood was collected and stored in EDTA-coated tubes at 4 °C for a complete blood count test [92]. Analysis was performed with 20 μL blood samples using the Heska HemaTrue Analyzer and HeskaView Integrated Software version 2.5.2.

Statistical analyses were performed in GraphPad Prism 9 using two-way ANOVA with Tukey's multiple comparisons test for tumor growth curves, one-way ANOVA for CBC parameters, and Welch's two-tailed *t*-test for endpoint tumor weights.

6. Chan SS-K, Shi X, Toyama A, Arpke RW, Dandapat A, Iacovino M, et al. Mesp1 patterns

mesoderm into cardiac, hematopoietic, or skeletal myogenic progenitors in a context-dependent manner. Cell Stem Cell. 2013;12:587–601.

16. Matreyek KA, Stephany JJ, Chiasson MA, Hasle N, Fowler DM. An improved platform for functional assessment of large protein libraries in mammalian cells. Nucleic Acids Res.

2020;48:e1.

29. Langmead B, Salzberg SL. Fast gapped-read alignment with Bowtie 2. Nat Methods.

2012;9:357–9.

41. Abbas-Aghababazadeh F, Li Q, Fridley BL. Comparison of normalization approaches for gene expression studies completed with high-throughput sequencing. PLoS One.

2018;13:e0206312.

42. Signorell A. DescTools: Tools for descriptive statistics. R package version 0.99. 20. CRAN. 2017;

54. Krueger F. Trim Galore: a wrapper tool around Cutadapt and FastQC to consistently apply quality and adapter trimming to FastQ files. Babraham Bioinformatics. 2019;

75. Salkovic E, Sadeghi MA, Baggag A, Salem AGR, Bensmail H. OutSingle: a novel method of detecting and injecting outliers in RNA-Seq count data using the optimal hard threshold for singular values. Bioinformatics. 2023;39:btad142.

82. Kim M, Pyo Y, Hyun S-I, Jeong M, Choi Y, Kim VN. Exogenous RNA surveillance by proton-sensing TRIM25. Science. 2025;388:eads4539.
