## Supplementary Figures for "A Proteome-Wide Gain-of-Function Atlas Mapping the Landscape of Chemotherapy Response"

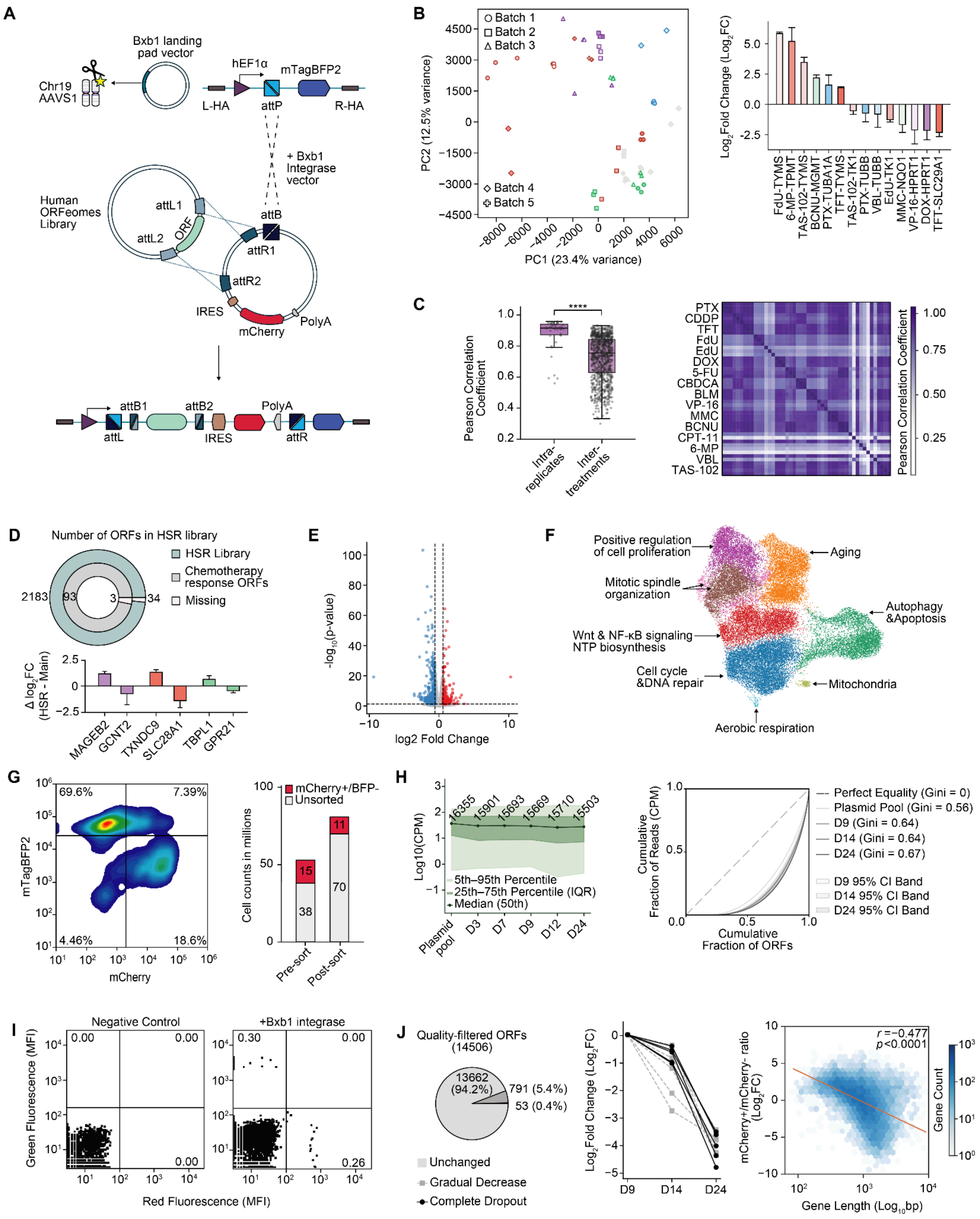


**Fig. S1. Technical validation and quality control of the proteome-wide GOF screening platform. (A)** Molecular schematic of the Bxb1-landing pad human ORFeome-integrated system for a proteome-wide Gene Overexpression (BOGO) architecture. **(B)** Global profiling validation and batch reproducibility. Left: PCA evaluating variance distribution across five independent experimental sequencing batches. Right: Validation of canonical drug-gene response. **(C)** Screen reproducibility and inter-treatment distinctiveness. Left: Comparison of Pearson correlation coefficients between intra-replicates and inter-treatment conditions.

Right: Pairwise Pearson correlation heatmap across all 16 chemotherapeutic modalities. **(D)** High-saturation retesting (HSR) library composition (top) and differential response validation by $log_2FC$ of HSR screening minus original chemotherapy screening (main) for validating genes. **(E)** Volcano plot highlighting differentially expressed genes in mRNA-seq of DOX 120-hour treatment. **(F)** Single-cell transcriptomic UMAP projection annotated with major biological pathways and functional modules driven by GOF perturbations. **(G)** Efficiency of ORF integration in HeLa cells measured by mCherry and mTagBFP2 (BFP) fluorescence. Left: FACS gating strategy. Right**:** Stacked bar graph showing the number of sorted and unsorted cells via mCherry fluorescence. **(H)** Library representation metrics over longitudinal baseline culture. Left: Abundance distributions from plasmid pool through day 24 post-transfection in PCR1-prepared samples. Right: Lorenz curves depicting cumulative read distributions and Gini coefficients across time points in nested PCR samples. **(I)** Co-transfection assay validates the single integration site. **(J)** Essentiality dropout dynamics and length bias analysis. Left: Pie chart showing that around 6% of quality-filtered ORFs show a gradual decrease or complete dropout. Middle: Trajectory line plot showing normalized $log_2FC$ to D9 across time points for the top 5 genes in the gradual decrease and complete dropout ORFs. Right: Inverse correlation between gene length and mCherry+/mCherry- ratio. Statistical significance in panel C was evaluated by a two-tailed unpaired Student's t-test; **** $p < 0.0001$.

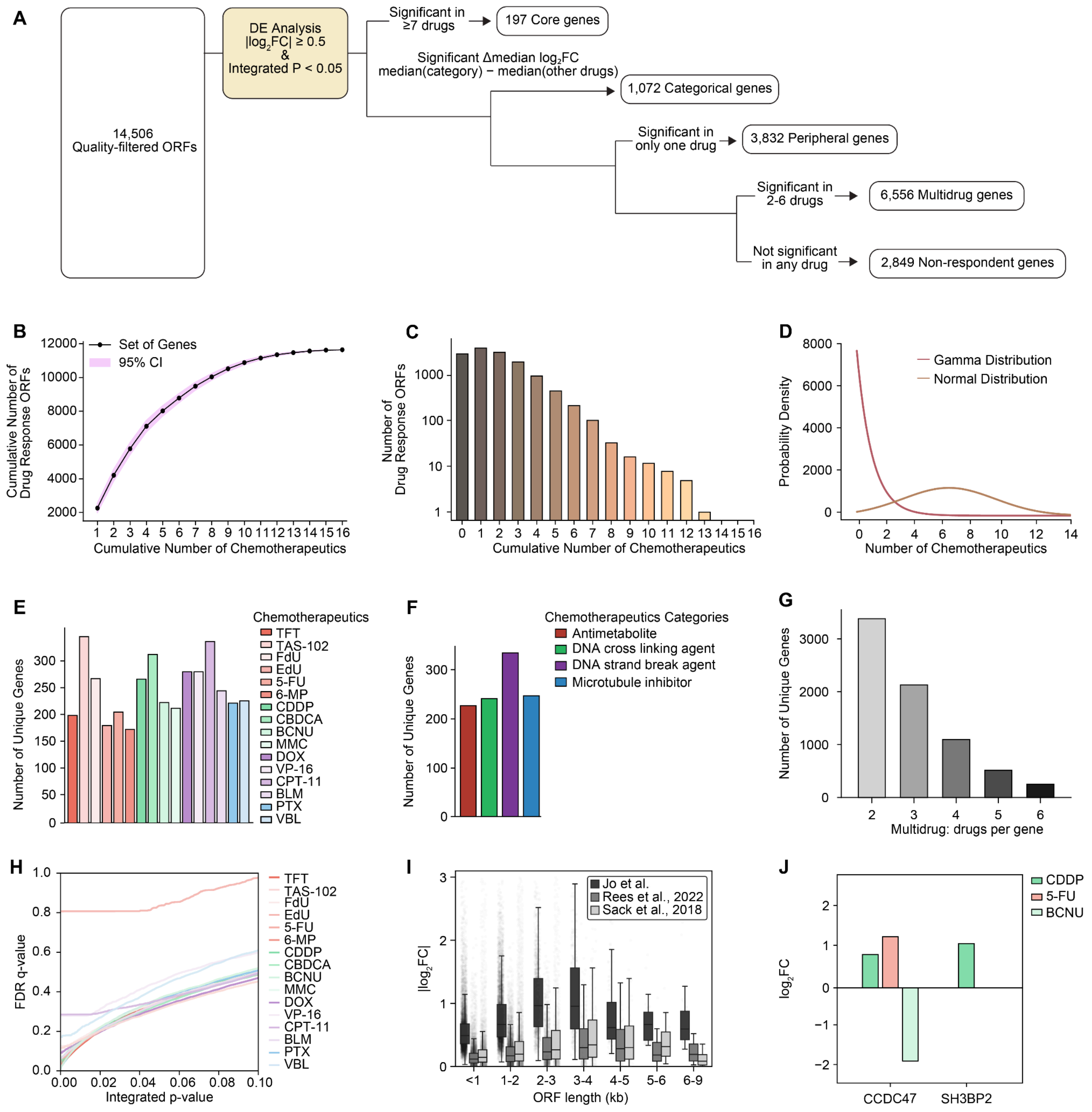


**Fig. S2. Methodological workflow, distribution modeling, and functional characterization of screened ORFs. (A)** Step-by-step pipeline flowchart illustrating the systematic filtering and classification workflow from 14,506 quality-filtered ORFs into functional categories (Core, Categorical, Peripheral, Multidrug, and Non-respondent genes) based on differential expression analysis ($|\log_2 FC| \geq 0.5$ and integrated *p*-values < 0.05) and intersection thresholds. **(B)** Cumulative distribution of unique drug-response-modulating ORFs across an increasing number of tested chemotherapeutic agents, compared with a 95% confidence interval model. **(C)** Histogram showing the distribution trends of candidate counts according to the cumulative number of chemotherapeutic agents. **(D)** Mixture modeling of the distribution of responsive drug counts per ORF using gamma and normal distributions. **(E)** Distribution profile highlighting the number of unique ORFs identified across individual chemotherapeutic agents. **(F)** Distribution of unique ORF counts across drugs grouped by common mechanisms of action (Antimetabolite, DNA cross-linking agent, DNA strand break agent, and Microtubule inhibitor). **(G)** Distribution of unique ORFs per drug count in the Multidrug category (ranging from 2 to 6 drugs). **(H)** Comparison curves of integrated *p*-values versus FDR *q*-values across individual chemotherapeutic agents. **(I)** Boxplot evaluating $|\log_2 FC|$ across 1-kb ORF length intervals (up to 9 kb), compared across independent datasets (Sack et al., 2018 and Rees et al., 2022). **(J)** Bar chart displaying $\log_2 FC$ values for representative candidates (*CCDC47* and *SH3BP2*) across selected chemotherapeutic treatments (CDDP, 5-FU, and BCNU).

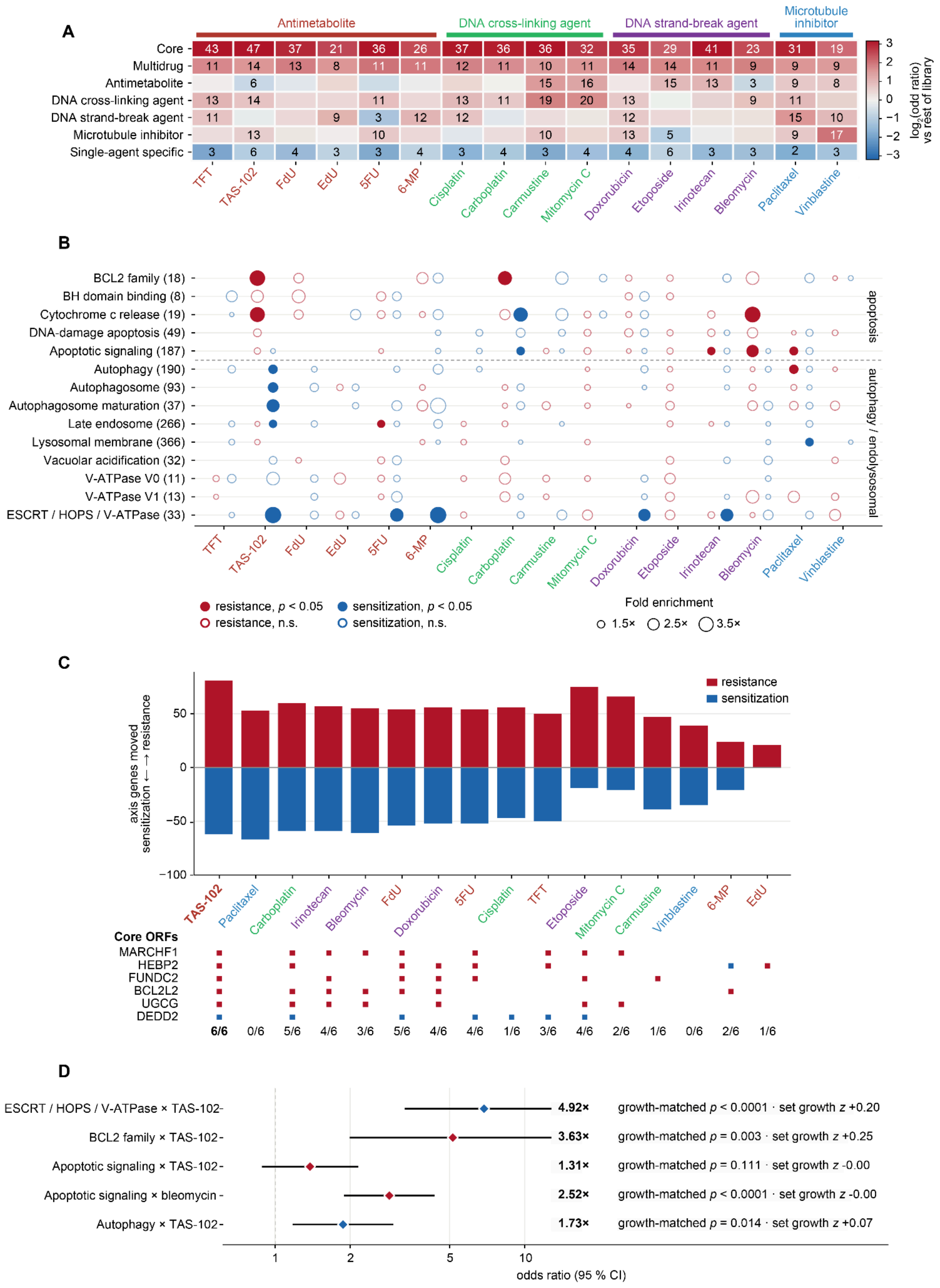


**Fig. S3. Chemotherapy response and ORF GO enrichment dynamics in the *BCL-2* family-associated autophagy-endolysosomal axis. (A)** Heatmap showing resistance enrichment ($\log_2$ odds ratio versus the rest of the measured library) across functional ORF categories and 16 chemotherapeutic agents grouped by mechanism of action category, with cell numbers indicating the percentage (%) of measured ORFs conferring significant resistance ($\log_2 FC \geq 0.5$ and integrated $p$-values < 0.05). **(B)** Dot-plot matrix of 444 one-sided hypergeometric enrichment tests across 14 pre-specified apoptotic, autophagic, and endolysosomal gene sets (rows; number of library ORFs per set in parentheses) and 16 chemotherapeutic agents (columns) in both response directions (red, resistance; blue, sensitization; filled circles, $p < 0.05$; open circles, non-significant; circle size, fold enrichment). **(C)** Bidirectional distribution of resistance (red, upper bars) and sensitization (blue, lower bars) hits across the 842 ORFs annotated to the combined BCL-2 family, autophagy, and lysosome axis across all 16 chemotherapeutic agents (top), alongside the response direction and engagement fraction of the six Core ORFs of the axis (*MARCHF1*, *HEBP2*, *FUNDC2*, *BCL2L2*, *UGCG*, and *DEDD2*). **(D)** Forest plot of odds ratios (diamonds; red, resistance; blue, sensitization) and 95% confidence intervals for headline pathway enrichments, annotated with fold enrichment, empirical proliferation-matched permutation $p$-value (20,000 draws matched by DMSO growth z-score), and median set growth z-scores.

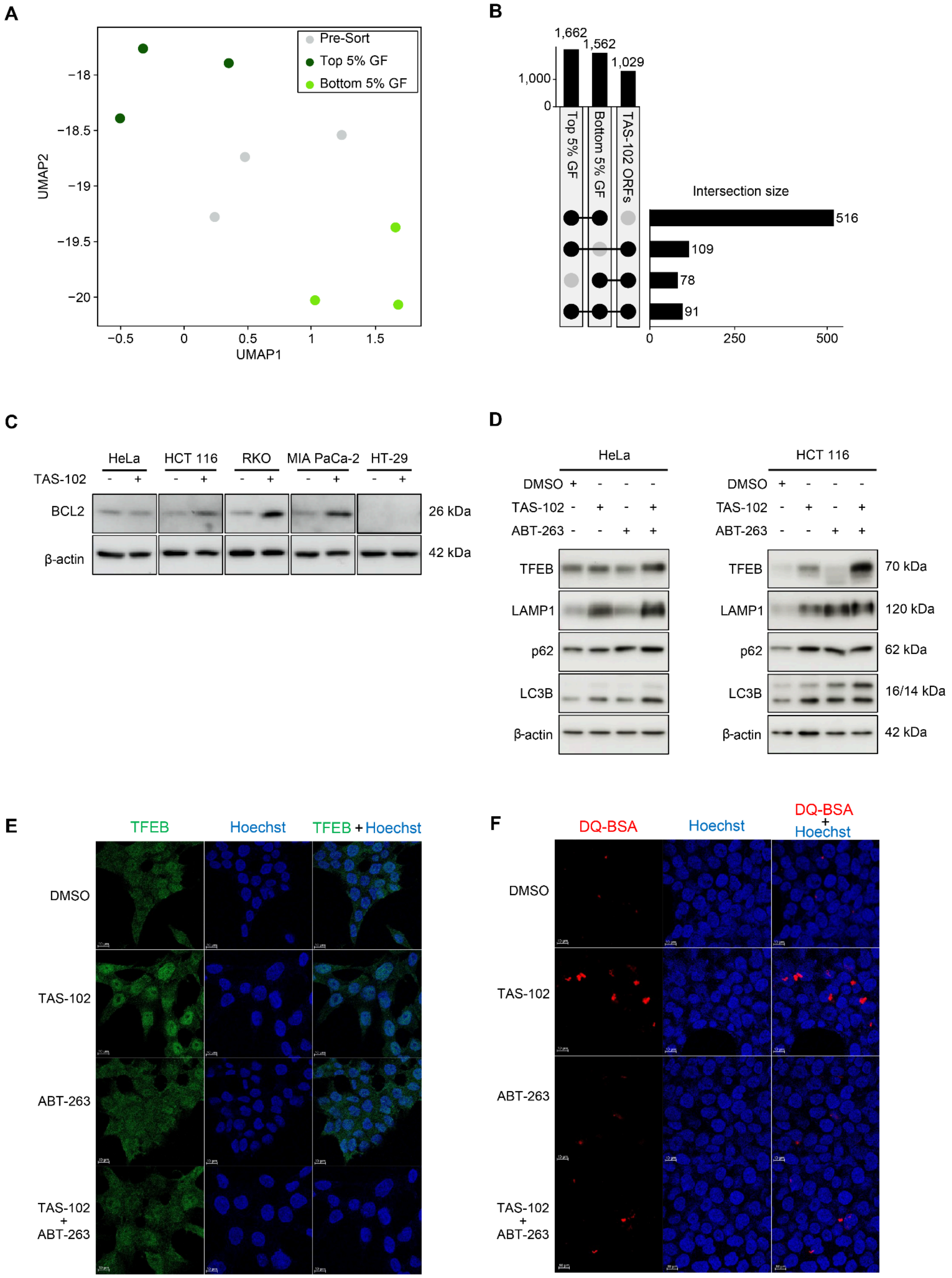


**Fig. S4. Characterization of lysosomal modulation. (A)** UMAP clustering plot displaying the distribution and reproducibility of sorted populations (top 5% high green fluorescence, bottom 5% low green fluorescence, and pre-sort samples) from the LysoTracker screen. (**B**) UpSet intersection plot showing the overlap between identified lysosomal activators, repressors, and TAS-102-specific drug response profiles. (**C**) Western blot analysis of BCL-2 protein expression in HeLa (0.8 μM), HCT 116 (2 μM), RKO (0.6 μM), MIA PaCa-2 (2 μM), and HT-29 (2 μM) cells following 24 h of DMSO (–) or TAS-102 (+) treatment, with β-actin as a loading control. (**D**) Western blot analysis of lysosomal and autophagy pathway proteins (TFEB, LAMP1, SQSTM1/p62, and LC3B, with β-actin as a loading control) in HeLa and HCT 116 cells treated with DMSO, TAS-102 (1 μM), ABT-263 (1 μM for HeLa; 0.5 μM for HCT 116), or their combination for 48 h. (**E**) Representative confocal immunofluorescence images (scale bars: 10 μm) of TFEB nuclear translocation (green) co-stained with Hoechst 33342 (blue) in HCT 116 cells treated with DMSO, TAS-102 (2 μM), ABT-263 (0.5 μM), or their combination for 24 h. (**F**) Representative confocal fluorescence images (scale bars: 10 μm) of DQ-BSA proteolysis assays (red) co-stained with Hoechst 33342 (blue) assessing lysosomal hydrolytic activity in HCT 116 cells treated with DMSO, TAS-102 (2 μM), ABT-263 (0.5 μM), or their combination for 24 h.

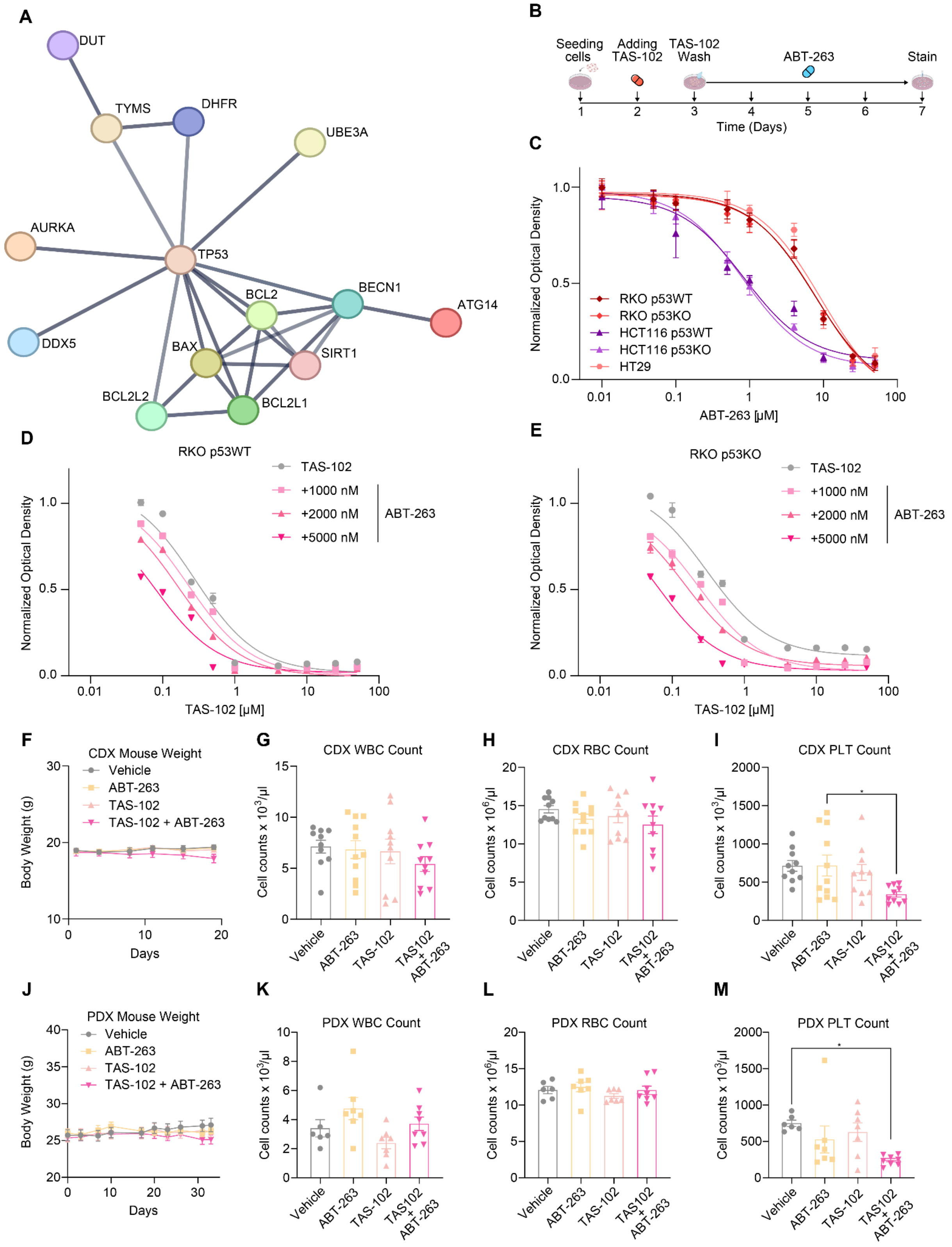


**Fig. S5. Synergistic efficacy and safety profile of TAS-102 and ABT-263 combination. (A)** STRING PPI network illustrating the canonical TP53-centered interaction bridge connecting thymidylate synthase (TYMS) to the BCL-2 family and core autophagy regulators (*BECN1*, *ATG14*; high confidence, > 0.7 protein interaction score from STRING; < 5 interactors in the first and second shells). **(B)** Schematic timeline of the experimental design and treatment schedule for the combination therapy. **(C)** Representative dose-response curves for ABT-263 in colorectal cancer cell lines (HCT 116, RKO p53 WT and KO, and HT-29) across concentrations ranging from 0 to 50 μM. **(D–E)** Cytotoxicity evaluation of TAS-102 and ABT-263 combination therapy in **(D)** RKO p53 WT and **(E)** RKO p53 KO cells. (**F–I**) Longitudinal mouse body weight tracking (**F**) and endpoint complete blood count (CBC) parameters, including white blood cells (WBC; **G**), red blood cells (RBC; **H**), and platelets (PLT; **I**) in the HCT 116 CDX model. (**J–M**) Longitudinal mouse body weight tracking (**J**) and endpoint WBC (**K**), RBC (**L**), and PLT (**M**) counts in the colorectal PDX model. Data in panels C–E are presented as mean ± SD, whereas panels F–M show mean ± SEM. Statistical analysis was performed using one-way or two-way ANOVA with Tukey's multiple comparisons test; *$p < 0.05$.
